# DeepD enables efficient genome modeling and controllable long-context sequence generation across domains of life

**DOI:** 10.64898/2026.10.13.758170

**Authors:** Beibei Xin, Zijian Wang, An Yu, Ziyang Chen, Yucheng Guo, Mingpei Zhang, Bozhen Hu, Chonghang Ye, Jian Chen, Qiujie Liu, Haiming Zhao, Weibin Song, Nanqing Dong, Fan Yang, Qirong Yang, Hainan Zhao, Jinsheng Lai

## Abstract

Genome foundation models hold great potential for advancing biological research, but modeling diverse complex genomes and generating controllable genome-scale sequences remain challenging. We present DeepD, a phylogeny-informed sparse mixture-of-experts model operating at single-nucleotide resolution over a 1-million-token context while reducing active computation. Pretrained on unselected whole-genome sequences across all domains of life, DeepD learns genic and intergenic sequence features and supports broad sequence-to-function prediction across species. DeepD performs well in human protein fitness and disease variants effect prediction, such as *BRCA1* and *BRCA2*, with particular strength in regulatory regions, such as promoter and transcription-factor binding sites. Expert routing captures exon–intron boundaries, transposable elements, and distal *cis*-regulatory elements. DeepD also generates experimentally validated core promoters with specified transcriptional strengths. Finally, the Context–Anchor–Body strategy enables generation of complete yeast chromosome sequences and megabase-scale *Arabidopsis* segments, preserving gene structure and order while introducing variations within natural ranges. Together, DeepD is an efficient genome foundation model that unifies accurate sequence-to-function prediction with controllable genome-scale generation, advancing predictive and generative design of complex genomes.

## Introduction

Genomes encode the instructions of life and reflect its evolutionary diversity. Genome models have developed with distinct emphases.^1–6^ Self-supervised DNA language models learn sequence constraints from unlabeled genomes: masked-language models such as Nucleotide Transformer (NT) generate transferable representations for downstream prediction and variant prioritization,^4^ whereas autoregressive models such as Evo and Evo2 support nucleotide-likelihood scoring and sequence generation to some extent.^3,5^ In parallel, sequence-to-function models such as AlphaGenome focus on human and mouse functional genomics to predict molecular phenotypes and regulatory variant effects.^7^ An ideal genome model would intrinsically unify accurate sequence-to-function prediction and controllable sequence generation across different domains of life.

However, training and inference of models spanning all domains of life over long-context sequences require substantial computational resources, particularly at single-nucleotide resolution.^3^ These computational demands hinder the scalable modeling of phylogenetically diverse genomes over long genomic contexts, further limiting their broad portability. Sparse mixture-of-experts (MoE) architectures, which are successful in natural large language modeling, as exemplified by DeepSeek,^8,9^ expand model capacity while limiting active computation by routing each token to only a subset of experts.^10,11^ Their potential for modeling highly complex genomes across different domains of life remains unexplored. A phylogeny-informed MoE architecture could allocate model capacity across evolutionary lineages and long genomic contexts, enabling efficient training and inference at a broad phylogenetic scale.

The success of a foundation model depends not only on efficient architecture, but also heavily depends on data composition and volume.^12–14^ So far, most of the genome foundation models were pretrained either on sampled genomic sequences^15,16^ or on gene-centric and particular regulatory regions.^2,3,6,17–19^ To fully represent the complexity of genome-wide features, it is preferable to train a model on unselected whole-genome sequences encompassing all genic regions and entire intergenic regions, without downsampling intergenic (regulatory and repetitive) regions. Such comprehensive pretraining could strengthen prediction in non-coding or regulatory regions, capture features ranging from exon–intron boundaries to DNA transposons, LTR retrotransposons, and distal *cis*-regulatory elements (CREs), and support coherent long-sequence generation in complex eukaryotic genomes.

Beyond prediction and interpretation, genome foundation models offer a route towards programmable DNA design across different scales. Several earlier generative models have produced proteins, genes, and *cis*-regulatory elements around a few kilobases in length,^20–25^ while recent models have generated longer DNA sequences, including multicomponent genetic systems, bacteriophage genomes, and yeast chromosomes.^3,5,26–28^ Yet early errors in autoregressive generation can propagate with sequence length, progressively reducing long-range coherence and controllability.^29,30^ This limitation is particularly consequential for genomic sequences over hundreds of kilobases, where genomic grammar remains incompletely understood. Existing approaches based on continuation from short prompts or predictor-guided search provide limited control over the placement and organization of internal genomic elements.^5^ Moreover, eukaryotic chromosome-scale design must coordinate gene structure, gene order, and intergenic variations.^31,32^ Natural language modeling has addressed analogous long-sequence challenges through conditional infilling and recurrent context strategies.^33,34^ Adapting these principles, we developed a Context–Anchor–Body (CAB) strategy that generates a Body between a downstream Context and an upstream Anchor and iteratively advances these constraints beyond a single context window.

Here, we present DeepD, a phylogeny-informed genome foundation model that combines efficient sparse computation, unselected whole-genome pretraining, and CAB-based controllable generation. DeepD was trained at two scales: a 1.8-billion-parameter model on 1 trillion tokens (DeepD-2B) and a 63-billion-parameter model on 5.69 trillion tokens (DeepD-60B). The DeepD-60B pretraining corpus comprised approximately 5.90 Tb of nucleotide sequence spanning the three domains of life together with viruses, including 5,434 eukaryotic, 6,932 archaeal, 136,485 bacterial, and 2,917,521 viral genomes. DeepD integrates broad sequence-to-function prediction and interpretable genomic representations with experimentally validated regulatory-element design, chromosome-scale generation in yeast, and megabase-scale generation in *Arabidopsis*. Together, these capabilities establish DeepD as an efficient foundation model for accurate sequence-to-function prediction and coherent genome-scale generation across all domains of life.

## Results

### DeepD enables efficient long-context modeling

We developed DeepD to model genomes as they occur in nature: long, heterogeneous, evolutionarily diverse, and dominated by both coding and non-coding sequence. DeepD was designed to meet four biological requirements for genome-scale modeling: efficient long-context computation, phylogenetic representation, unselected whole-genome pretraining, and controllable generation (**Figure 1A**). DeepD was trained at two scales: a smaller version with 1.8 B parameters trained on 1 trillion (T) tokens (DeepD-2B), and a larger version with 63 B parameters trained on 5.69 T tokens (DeepD-60B). DeepD-60B was trained on nucleotide sequences spanning eukaryotes, archaea, bacteria, and viruses, including 5.39 Tb from 5,434 eukaryotic species, 12.09 Gb from 6,932 archaeal species, 452.60 Gb from 136,485 bacterial species, and 45.22 Gb from 2,917,521 viral species (**Figure 1A**). The eukaryotic partition includes 3,600 animal genomes, 928 plant genomes, 767 fungal genomes, and 139 protist genomes (**Figure S1A**), providing broad phylogenetic coverage across major kingdoms. Rather than enriching for genic regions, we designed the training corpus as unselected whole-genome data. As a result, DeepD was exposed to both annotated and unannotated sequence classes, including coding sequence (CDS), introns, repeats, other intergenic regions, and viral sequences (**Figure 1A, 1C** and **S1B**). Non-coding and repetitive sequences constitute a substantial fraction of the corpus, especially in eukaryotes, ensuring that the model learns genome-wide organization rather than genes alone (**Figure S1B**).

**Figure 1.**
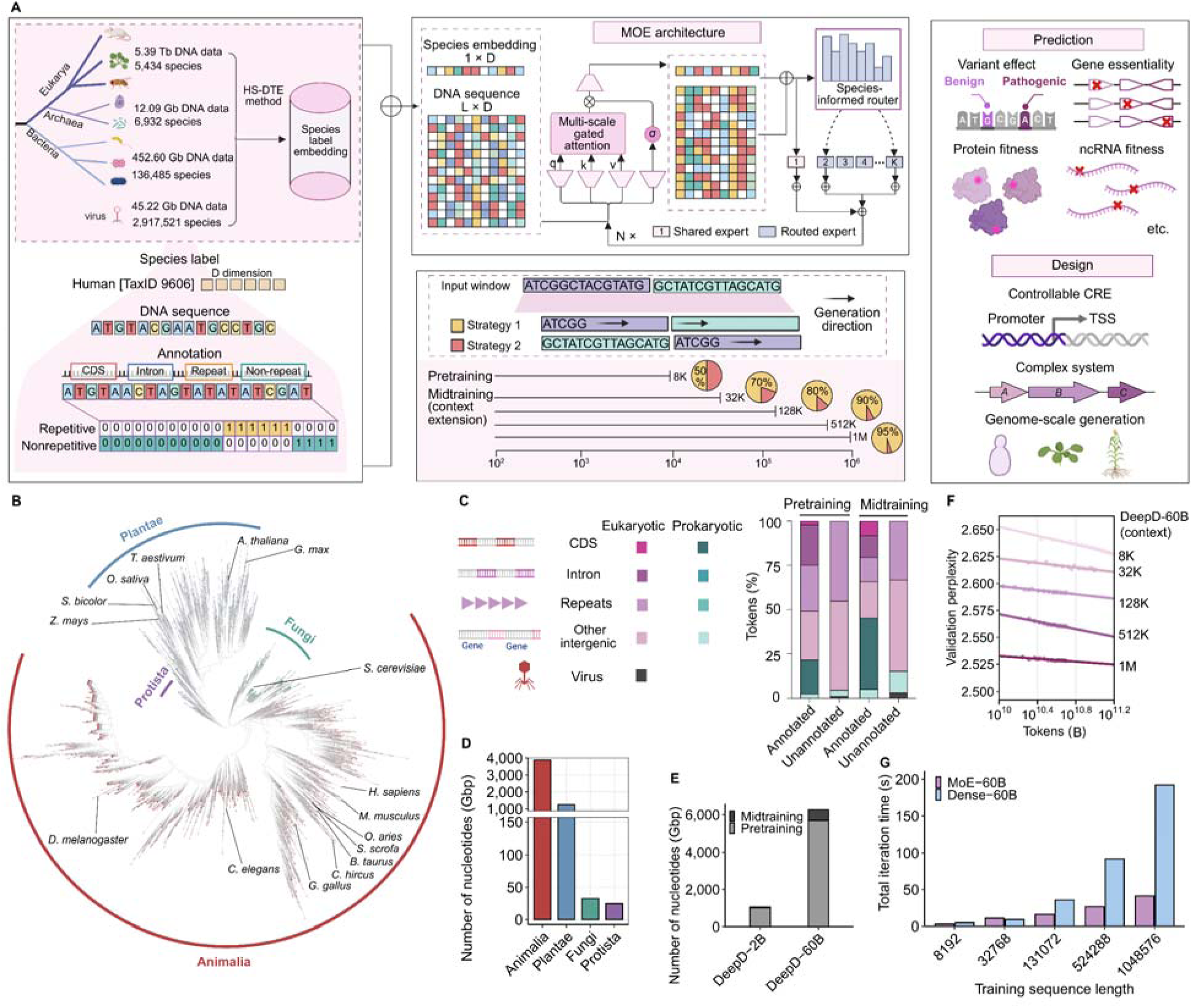
Overview of DeepD model architecture, pretraining corpus, and long-context scaling. (A) Schematic of the DeepD genome foundation model. DeepD was trained on 5.39 Tb from 5,434 eukaryotic species, 12.09 Gb from 6,932 archaeal species, 452.60 Gb from 136,485 bacterial species, and 45.22 Gb from 2,917,521 viral species. The model integrates DNA sequence, species label embeddings, and genome annotation features through a sparse MoE architecture. Multi-scale gated attention combines 24 sliding-window attention heads (SWA) with 8 full-attention heads (FA), while a species-informed router dynamically activates experts conditioned on phylogeny. (B) Phylogenetic tree of 5,434 eukaryotic genomes in the pretraining corpus. Major eukaryotic species in the following four kingdoms are highlighted: Animalia, Plantae, Fungi, and Protista. (C) Distribution of tokens in pretraining and midtraining stages, including the percentage of CDS, introns, repeats, and other intergenic regions in both annotated and unannotated genomes. (D) Number of tokens in each kingdom of NCBI genomes. (E) Number of tokens for pretraining and midtraining of DeepD-2B and DeepD-60B. (F) Validation perplexity of DeepD-60B midtraining across context lengths, showing benefits with increasing context length. (G) Comparison of iteration time on 64 GPUs for DeepD-60B versus a dense 60B model, showing that the sparse MoE architecture reduces computation time.

DeepD integrates DNA sequence, species label embeddings, and annotation features within a sparse MoE architecture (**Figure 1A**). The model uses a species-informed router and a multi-scale gated attention module. In each block, 24 of 32 attention heads use sliding-window attention, whereas 8 heads use full attention (**Figure S1C**). This design captures both local sequence patterns and longer-range dependencies. To extend modeling across genomic length scales, we trained DeepD in two stages. Pretraining was performed at 8K short context. Midtraining then progressively extended the context window from 8K to 32K, 128K, 512K, and 1M tokens (**Figure 1A** and **S1D**). The token distribution remained broadly stable across these stages, preserving representation of both annotated and unannotated genomic features as context length increased (**Figure S1D**).

We next examined optimization and scaling behavior. Perplexity decreased steadily during training for both the 2B and 60B models, with the larger model consistently achieving lower perplexity (**Figure S1E**). In the long-context training pipeline, validation perplexity continued to improve as context length increased from 8K to 1M (**Figure 1F**), indicating that larger context windows provide measurable gains for genome language modeling. We then compared the computational cost of MoE-60B with a dense model at matched parameter scale. Iteration time increased sharply with sequence length in both cases, but the increase was substantially smaller for the sparse MoE model, and the gap widened further at longer contexts (**Figure 1G**). Thus, ultra-long context imposes a steep computational burden on dense architectures, whereas sparse routing makes long-context genome modeling more practically accessible. Together, these results establish DeepD as an efficient, unselected whole-genome foundation model for learning across species and genomic scales.

### Phylogeny-informed modeling captures species-specific coding rules

Genomic GC content is a major genomic signature shaped by evolutionary pressure and serves as a primary feature for distinguishing diverse species and lineages.^35^ To investigate whether DeepD learns the species-specific genomic rules and whether these rules can be used to guide programmable sequence generation, we selected a representative set of species spanning a wide genomic GC spectrum for mixture-of-experts (MoE) interpretability analysis (**Figure 2A** and **S2A**).

**Figure 2.**
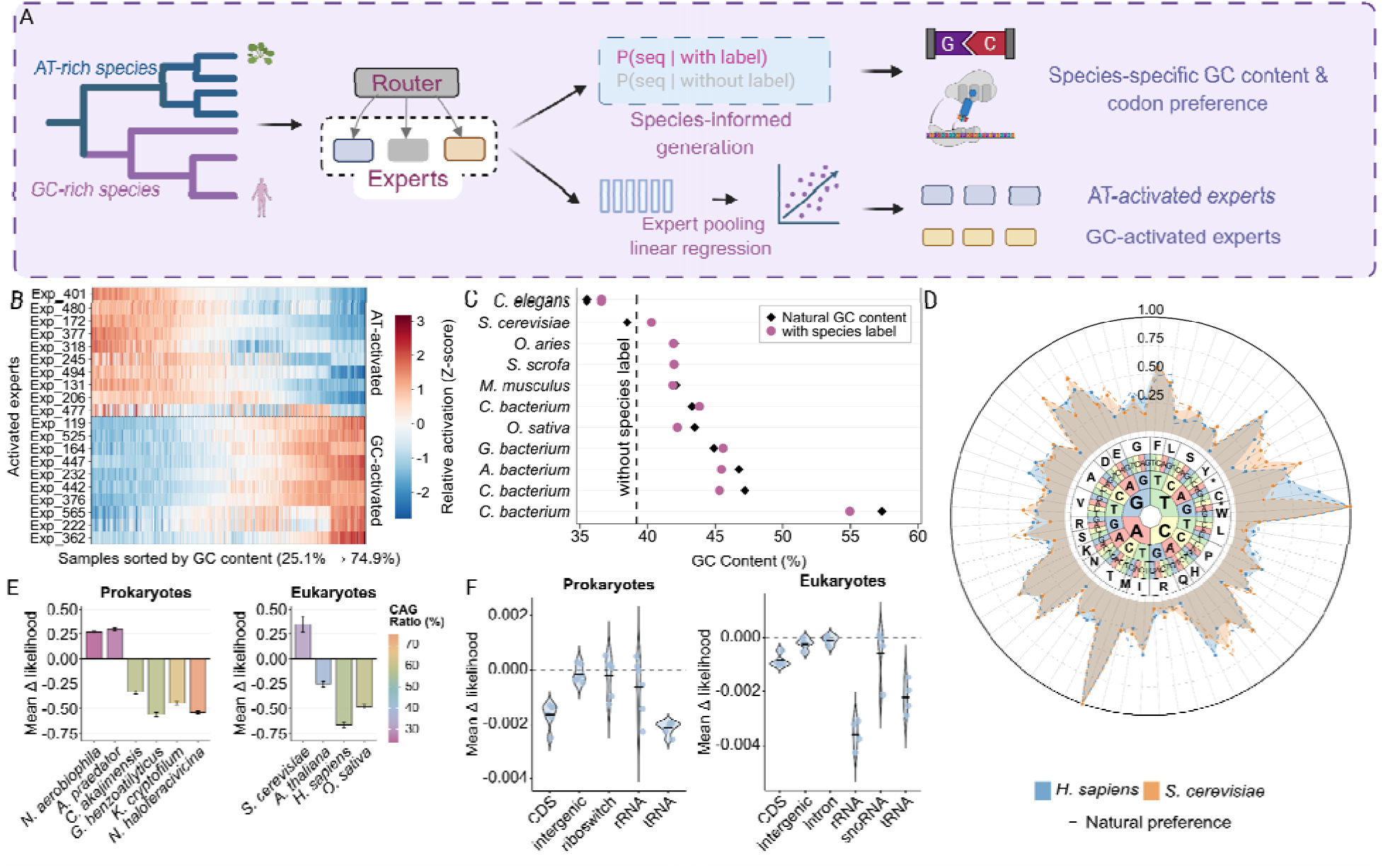
Phylogeny-informed modeling and zero-shot mutational probing of species-specific genomic constraints. (A) Schematic overview of the DeepD framework for phylogeny-guided sequence generation and interpretability analysis. The routing mechanism within the Mixture-of-Experts (MoE) architecture guides sequence processing across diverse phylogenetic levels (left). Two primary downstream analyses are depicted: species-conditioned sequence generation targeted at recovering taxon-specific GC content and codon preferences (top right), and the identification of composition-sensitive expert clusters (e.g., AT-activated and GC-activated experts) using Ridge regression-based expert pooling (bottom right). (B) Spatial activation patterns of Layer 2 experts across a genomic GC gradient. The heatmap represents the relative activation (Z-score) of experts in Layer 2, with input sequence samples sorted in ascending order of genomic GC content (from 0.25 to 0.75). The expert network spontaneously partitions into distinct AT-activated (top) and GC-activated (bottom) clusters in an unsupervised manner. (C) Genomic GC content of natural versus model-generated sequences across diverse taxa. Comparison of GC content (%) among natural genomic sequences (black diamonds), sequences generated using species-specific tags (pink circles), and sequences generated using the baseline “unlabeled” tag (dashed vertical line, representing a reference baseline of approximately 40%). (D) Recapitulation of species-specific codon usage profiles in generated sequences. Radar chart comparing the RSCU profiles of natural references (solid and dashed black lines) with sequences generated by DeepD under *H. sapiens* (blue shaded area) and *S. cerevisiae* (orange shaded area) conditioning across the 64-codon landscape. (E) Zero-shot mutational probing of codon-specific evolutionary constraints. Mean change in log-likelihood (Mean △likelihood) resulting from a simulated G-to-A transition at the third position of CAG codons in prokaryotes (left) and eukaryotes (right). Bar colors correspond to the natural genomic CAG preference (CAG Ratio, %) of each respective species. Error bars indicate standard deviation (s.d.). (F) Model sensitivity to simulated deletions across diverse functional genomic elements. Violin plots illustrating the mean change in log-likelihood (Mean △likelihood) following simulated 10-bp deletions in various functional genomic regions (including CDS, intergenic regions, riboswitches, rRNA, tRNA, introns, and snoRNAs) for prokaryotes (left) and eukaryotes (right). Dashed lines indicate zero baseline change.

We predicted sequence-level GC content layer by layer using Ridge regression with nucleotide-specific weight aggregation across experts (**Figure 2B** and **Figure S2B**). The low-level experts spontaneously act as efficient compositional sensors in shallow layers (**Figure S2B**). Spatial activation analysis of Layer 2 experts confirmed this physical basis. DeepD decodes genomic heterogeneity through the differential activation of distinct “AT-activated” and “GC-activated” expert clusters (**Figure 2B**). A clear transition in expert activation is observed as the GC content of input sequences progressively increases. These results suggest that DeepD encodes genomic compositional features at early network layers within its internal expert structure.

We next evaluated whether these learned compositional rules could be used for controlled generation with phylogeny-informed species labels. With the phylogenetic tags, DeepD generated sequences whose GC compositions closely matched the natural genomic backgrounds (**Figure 2C**). Generated GC contents were strongly correlated with that of natural genomes and thus recovered diverse compositions spanning a wide GC spectrum (R^2^ = 0.706 and Spearman’s ρ = 0.816, **Figure S2C**). By contrast, sequences generated without taxonomic information resulted in a consistent baseline GC content of approximately 40%. These results suggest DeepD uses phylogenetic tags to recover species-specific GC contents.

We then asked whether phylogeny-conditioned generation also preserves finer-scale codon usage bias. Given a specific species tag and a minimal 50-bp upstream prompt, DeepD generated 1,000-bp coding sequences. The Relative Synonymous Codon Usage (RSCU) analysis showed that synthetic sequences generated with *H. sapiens* and *S. cerevisiae* tags closely matched their natural codon usage across the full 64 codon space (**Figure 2D**). The model captured both the more GC-rich codon preferences of human coding sequences and the more AT-biased codon usage of yeast, indicating that species-conditioned generation extends beyond global nucleotide composition to synonymous codon preference.

To determine whether these codon constraints are encoded during zero-shot inference, we performed zero-shot likelihood perturbation analyses. We first conducted a mutational probing analysis focusing on the third position (G) of CAG codons within CDS regions (**Figure 2E**). Substitution from G to A produces the synonymous CAA codon and therefore directly illustrates species-specific codon preference. By quantifying the likelihood difference between A and G, we found that DeepD’s contextual predictions mirrored species-specific codon usage biases. In species with low CAG preference (∼30%), DeepD assigned higher likelihoods to the G-to-A transition. Conversely, in high-CAG species (∼60–70%), the model imposed a substantial likelihood penalty on A. Thus, DeepD captures species-dependent synonymous codon preference at single-nucleotide resolution without task-specific fine-tuning.

We further tested whether DeepD recognizes basic genomic grammar by introducing premature termination codons (PTCs) into coding sequences. The PTC analysis showed consistently reduced sequence log-likelihood across all evaluated prokaryotic and eukaryotic taxa (**Figure S2D**), indicating that DeepD identified premature stops as detrimental mutations. Finally, we extended this analysis using 10-bp deletions across diverse genomic elements (**Figure 2F**). In both eukaryotes and prokaryotes, the model predicted higher likelihood penalties in coding sequences and structured RNA elements such as rRNA and tRNA than in intronic or intergenic regions (**Figure 2F**). Deletions in introns and intergenic regions elicited minimal likelihood penalties, reflecting their relaxed evolutionary pressures. Together, these findings suggest that DeepD integrates global taxonomic context, local functional topography, and strict genomic grammar into a unified, interpretable sequence representation.

### DeepD enables broad sequence-to-function prediction across species

Functionally important sequence patterns are often conserved during evolution. By learning these conserved patterns from phylogenetically diverse genomes, genome foundation models acquire a likelihood landscape in which mutations that disrupt functional constraints tend to receive lower probabilities, providing a basis for zero-shot functional prediction without task-specific fine-tuning or supervision.^3–5^ We then asked whether DeepD could support zero-shot sequence-to-function prediction for three biological modalities (DNA, RNA, and protein) across species. We therefore evaluated DeepD on zero-shot benchmarks spanning bacterial proteins, eukaryotic proteins, non-coding RNAs (ncRNAs), and phage genes (see **Methods**; **Figure 3A**). Across seven bacterial protein fitness datasets, DeepD achieved a mean absolute Spearman correlation of 0.551, outperforming the evaluated Evo-series models^3,5^ and ProGen2 models,^24,36^ while remaining competitive with leading protein language models such as the ESM series.^37,38^ Across five eukaryotic protein datasets, DeepD achieved a mean absolute Spearman correlation of 0.478, comparable to the strongest genome and protein language models despite being trained on genomic sequences rather than protein-specific corpora. DeepD showed a particularly clear advantage for ncRNAs, achieving the highest mean correlation across seven mutational datasets spanning rRNAs, tRNAs, and ribozymes (0.297), outperforming both the Evo series and the dedicated RNA language model RiNALMo.^39^ DeepD also achieved an area under the receiver operating characteristic curve (AUROC) of 0.603 for phage essential-gene prediction, comparable to the best-performing Evo2 models. Together, these results demonstrate that DeepD recovers functional constraints across biological modalities and generalizes zero-shot sequence-to-function prediction across bacterial, phage, and eukaryotic systems.

**Figure 3.**
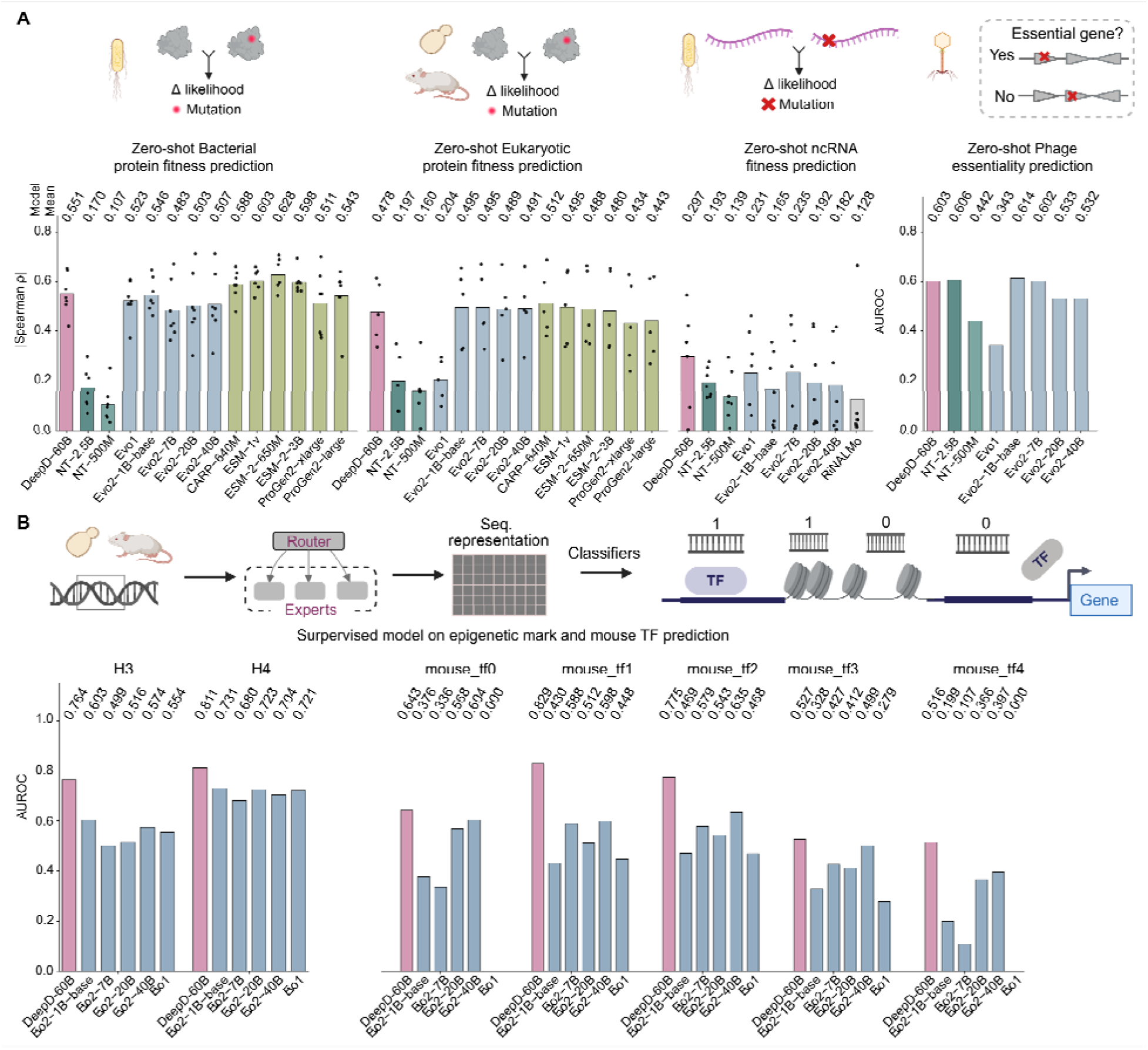
DeepD exhibits broad generalization across sequence-to-function prediction tasks. (A) Zero-shot sequence-to-function prediction across species and molecular modalities. DeepD-60B and baseline models were evaluated on bacterial protein fitness (seven datasets), eukaryotic protein fitness (five datasets), ncRNA fitness (seven datasets), and phage gene essentiality prediction. For the protein and ncRNA benchmarks, performance was measured by the absolute Spearman correlation (|Spearman ρ|) between experimentally measured mutational effects and model-derived changes in sequence likelihood (Δ likelihood). Dots represent individual datasets, bars indicate mean performance across datasets, and values above the bars denote model means. Phage essentiality prediction was evaluated using AUROC. Red bars indicate DeepD-60B. Blue bars indicate other genome language models, green bars indicate protein language models, and the gray bar indicates the RNA language model RiNALMo. (B) Performance of supervised classifiers built on top of genome model embeddings across non-coding regulatory genome tasks, evaluated by AUROC. Tasks include epigenetic mark prediction in yeast (H3 and H4) and TF binding prediction in mouse. Colored circles indicate the performance of individual genome foundation models, with DeepD-60B highlighted in red, Evo series models shown in blue.

Beyond likelihood-based zero-shot prediction, we next asked whether DeepD representations could support the supervised decoding of regulatory functions across species. Using model embeddings as input features, we trained downstream classifiers for histone-mark (H3 and H4) prediction in yeast and five transcription factor (TF) binding prediction tasks in mouse, and evaluated performance by AUROC (see **Methods**; **Figure 3B**). DeepD achieved AUROCs of 0.764 and 0.811 for H3 and H4 prediction, respectively, exceeding the strongest Evo-series comparators, which achieved 0.600 and 0.741. Across the five mouse TF binding tasks, DeepD ranked first on every task, with AUROCs ranging from 0.516 to 0.823 and improvements of 0.033 to 0.225 over the strongest comparator for each task. DeepD therefore showed its exceptional advantage in non-coding regulatory prediction, consistent with pretraining on unselected whole-genome data that retain full-space intergenic sequences.

Collectively, these analyses demonstrate the high capability of DeepD as a sequence-to-function model generalizing across species. DeepD recovered functional constraints across DNA, RNA, and protein sequences from bacterial, phage, and eukaryotic systems, while its embeddings consistently supported regulatory prediction in yeast and mouse. Its particularly strong performance on ncRNAs and non-coding regulatory elements further indicates that DeepD learns transferable representations beyond protein-coding regions.

### Human regulatory and variant-effect prediction

Predicting human variant effects is central to interpreting disease-associated genomes and prioritizing variants for experimental or clinical investigation, yet direct functional measurements are unavailable for most variants, particularly non-coding and non-SNV variants. DeepD addresses this gap by scoring the change in sequence likelihood between reference and alternative alleles, enabling zero-shot prediction without task-specific labels. We evaluated DeepD using human protein fitness, pathogenic and benign variants from ClinVar, splice-altering variants from SpliceVarDB, and functional effects of *BRCA1* and *BRCA2* variants (see **Methods**; **Figure 4A–D**).^40–44^ To contextualize the performance of DeepD, we compared it with currently published general genome language models from the Evo series,^3,5^ the multispecies DNA foundation models from NT version 2,^4^ and AlphaGenome,^7^ a human-focused sequence-to-function model trained on functional genomic readouts. Across five human variant-effect benchmarks, DeepD achieved overall AUROCs ranging from 0.814 to 0.965 and remained competitive with leading general and human-focused genome models.

**Figure 4.**
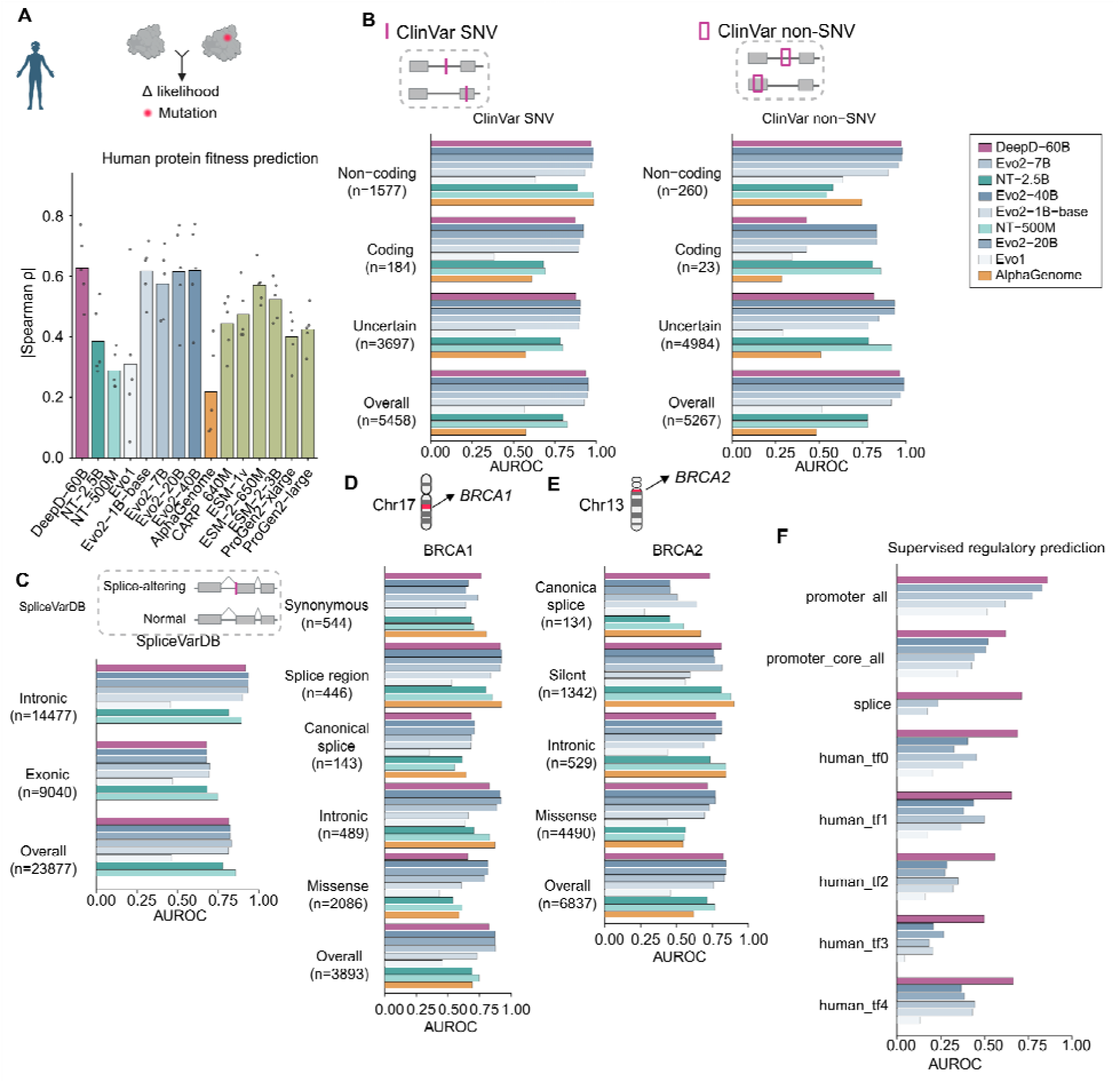
DeepD generalizes to human variant-effect and regulatory prediction. (A) Zero-shot prediction of human protein fitness from mutation-induced changes in sequence likelihood. Bars show the mean absolute Spearman correlation across datasets, and points represent individual datasets. (B) Zero-shot prioritization of pathogenic ClinVar SNVs (left) and non-SNVs (right), evaluated overall and stratified by coding annotation. (C) Zero-shot discrimination of experimentally validated splice-altering and control variants from SpliceVarDB, evaluated overall and for exonic and intronic variants. (D and E) Zero-shot prediction of functional effects for *BRCA1* (D) and *BRCA2* (E) variants, evaluated overall and across the indicated consequence classes. (F) Supervised prediction of human promoters, core promoters, splice sites, and five transcription factor-binding datasets using model-derived representations. Performance is reported as AUROC in (B–F). Models are colored as indicated in (B), and sample sizes in (B–E) denote the numbers of evaluated variants. NT, Nucleotide Transformer; SNV, single-nucleotide variant.

We first assessed zero-shot protein fitness prediction (**Figure 4A**). DeepD achieved a mean absolute Spearman correlation of 0.627 across the evaluated datasets. This substantially exceeded the NT-2.5B (0.385), NT-500M (0.289), and AlphaGenome (0.219). DeepD also slightly exceeded the strongest Evo-series model (Evo2-40B, 0.618) and the best-performing dedicated protein language model evaluated here (ESM-2-650M, 0.571). Thus, although DeepD was trained on genomic DNA rather than a protein-specific corpus, its sequence likelihoods retain information about the functional consequences of coding variation.

We next evaluated whether this capability extended to clinically annotated variants using ClinVar (**Figure 4B**). For single-nucleotide variants (SNVs), DeepD achieved an overall AUROC of 0.934, compared with 0.798 for NT-2.5B, 0.824 for NT-500M, and 0.576 for AlphaGenome. DeepD achieved AUROCs of 0.873 and 0.965 for coding and non-coding SNVs, respectively; on non-coding SNVs, its performance was comparable to NT-500M (0.980) and AlphaGenome (0.985). DeepD also performed strongly on non-SNVs, achieving an overall AUROC of 0.965, substantially exceeding NT-2.5B (0.779), NT-500M (0.779), and AlphaGenome (0.485). This advantage was particularly pronounced for non-coding non-SNVs, for which DeepD achieved an AUROC of 0.974, remaining close to the strongest Evo2 models (0.978–0.981). Area under precision-recall curve (AUPRC) analyses showed a similar pattern (**Figure S3**). These results indicate that DeepD generalizes across clinically annotated variant classes, with particular strength in non-coding non-SNVs.

Because many pathogenic variants disrupt RNA splicing, we further evaluated DeepD on experimentally validated splice-altering variants from SpliceVarDB (**Figure 4C**). DeepD achieved an overall AUROC of 0.814, comparable to the Evo2 models (0.812–0.833) and NT-500M (0.856), and exceeding NT-2.5B (0.779) and AlphaGenome (0.713). DeepD achieved AUROCs of 0.677 for exonic variants and 0.919 for intronic variants. Its intronic performance approached that of the strongest Evo2 model (0.936) and exceeded NT-2.5B (0.814), NT-500M (0.894), and AlphaGenome (0.722), whereas its exonic performance was comparable to AlphaGenome (0.682) (**Figure S3B**). Notably, AlphaGenome was trained with extensive supervision from human alternative-splicing readouts, whereas DeepD received no task-specific splicing supervision during pretraining.

We then focused on *BRCA1* and *BRCA2*, two clinically important tumor-suppressor genes with comprehensive experimental measurements of variant effects (**Figures 4D and 4E**). DeepD achieved overall AUROCs of 0.825 for both genes, exceeding NT-2.5B (0.693 and 0.712), NT-500M (0.747 and 0.765), and AlphaGenome (0.695 and 0.618), while remaining competitive with the strongest Evo2 models (0.877 and 0.844). For *BRCA1*, DeepD performed strongly on splice-region variants (AUROC = 0.915) and achieved the highest performance among general genome language models on synonymous variants, compared with 0.737 for the strongest Evo2 model and 0.707 for the strongest NT model. For *BRCA2*, DeepD achieved an AUROC of 0.813 for silent variants and the highest performance on canonical splice variants (0.730), exceeding AlphaGenome (0.668), the strongest NT model (0.548), and the strongest Evo2 model (0.507). Overall AUPRCs were 0.575 for *BRCA1* and 0.560 for *BRCA2*, further supporting the AUROC-based comparisons (**Figures S3C** and **S3D**). These results demonstrate that likelihoods learned through general whole-genome pretraining can distinguish functionally consequential variation within clinically important human genes.

Finally, we evaluated whether DeepD representations support supervised prediction of human regulatory functions (see **Methods**, **Figure 4F**). We trained downstream classifiers for human transcription factor (TF) binding, core-promoter and promoter detection, and splice-site prediction. Across five TF-binding tasks, DeepD achieved AUROCs of 0.497–0.688, outperforming the strongest general genome language model on each task. DeepD also led general genome language models in core-promoter, promoter, and splice-site prediction, achieving AUROCs of 0.620, 0.860, and 0.714, respectively. AlphaGenome performed better on TF binding (0.695–0.918) and splice-site prediction (0.815), consistent with its direct training on human functional genomic tracks, whereas DeepD performed better in both core-promoter (0.620 versus 0.601) and promoter detection (0.860 versus 0.692; **Figure S3E**). These results demonstrate that unselected whole-genome pretraining yields transferable representations of human regulatory and intergenic sequences.

Together, these results show that DeepD was competitive with leading published genome language models in zero-shot human protein-fitness and disease-associated variant-effect prediction, with particular strength in non-coding, splice-region, and intronic variants; AlphaGenome was generally less competitive in these zero-shot settings. In supervised classification of intergenic regulatory sequences, DeepD substantially outperformed all other general genome language models, demonstrating the transferability of representations learned from unselected whole-genome pretraining.

### Expert routing interprets coding, regulatory, and repetitive genomic features

Prediction alone does not explain how a genome model represents sequence organization. To examine whether DeepD’s sparse expert routing reflects genomic organization, we used MoE gating weights as internal features for annotation probing (**Figure 5A**). For each input position, we extracted 576-dimensional gating vectors from 23 layers, spanning layers 2 to 24. DeepD-60B (8K) was used for compact annotation tasks, whereas DeepD-60B (1M) was used for annotations expected to depend on long-range sequence context. For each annotation class, we trained a binary classifier on the gating vectors and selected the best-performing layer (see **Methods**). We applied this framework to three genomes with increasing structural complexity: the compact *E. coli* genome, the human genome, and the repeat-rich maize genome.

**Figure 5.**
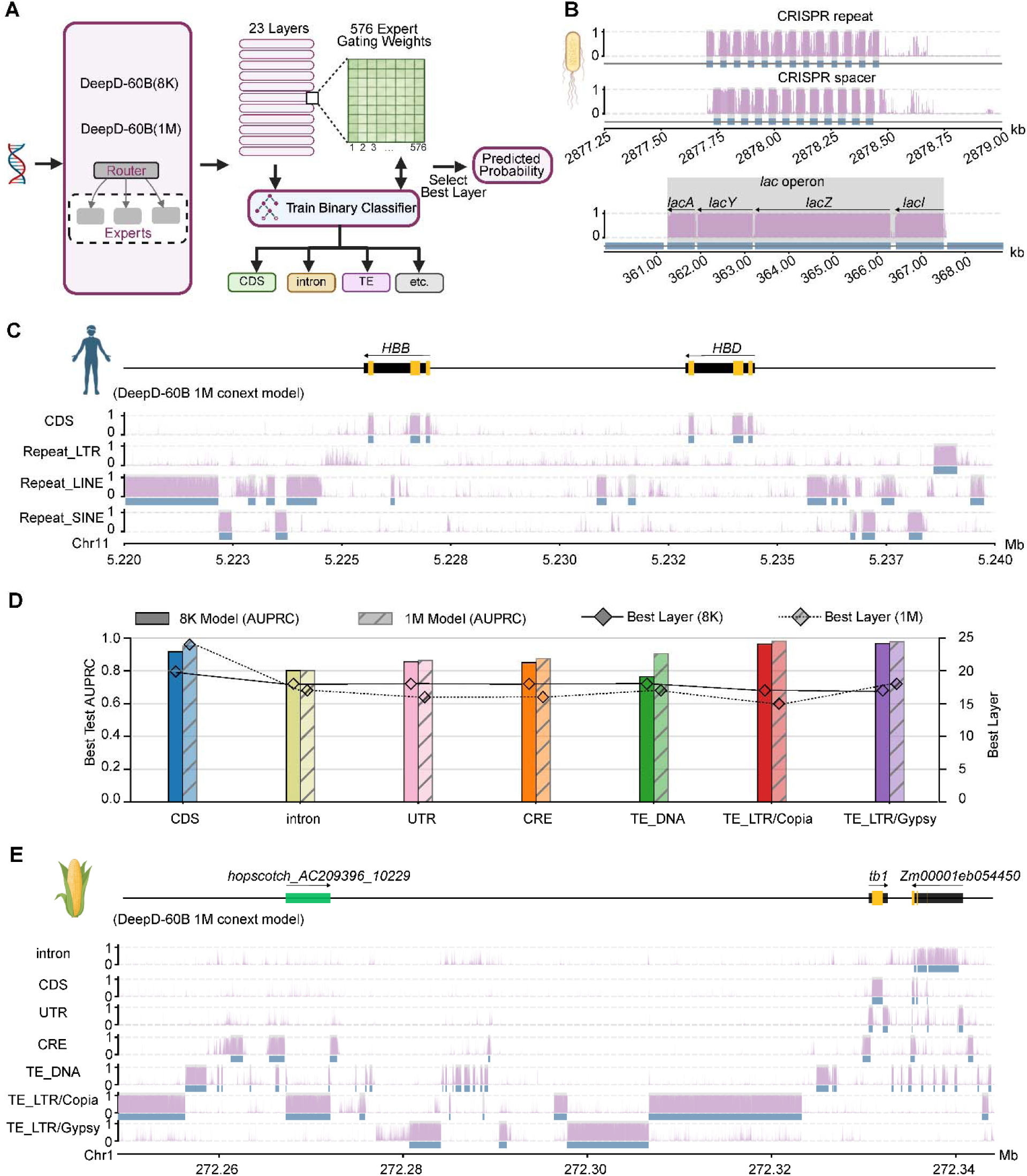
Expert routing in DeepD encodes functionally discrete genomic features across domains of life. (A) Schematic of the probing framework. Genomic sequences from *E. coli*, human, and maize are processed through DeepD-60B (8K) or DeepD-60B (1M). Expert gating weight vectors (576-dimensional) are extracted across 23 layers (2–24) and used to train annotation-specific binary classifiers. The best-performing layer per annotation category is selected to generate continuous predicted probability tracks across genomic loci. (B) High-resolution recovery of compact functional elements in *E. coli*. Predicted probability tracks (purple) with DeepD-60B (8K) routing weights are shown at two representative loci. Top: at a CRISPR array locus, probabilities for CRISPR repeat and spacer alternate in precise register with annotated structural boundaries (blue boxes), capturing the periodic internal organization of the array. Bottom: At the *lac* operon, gene-associated probability signals align accurately with the individual boundaries of *lacA*, *lacY*, *lacZ*, and *lacI* (grey shading, blue boxes). (C) High-resolution decoding of coding and repetitive features in the human genome with routing weights with DeepD-60B (1M). Predicted probability tracks (purple) from DeepD-60B (1M) are shown across the *HBB* gene cluster on chromosome 11. The model accurately reconstructs exon and CDS boundaries. Annotated regions are indicated by blue boxes below each track. (D) Layer-wise decoding of maize genomic features across context lengths. Bar plot (left y-axis): AUPRC of linear classifiers trained on representations with DeepD-60B (8K) (solid bars) and DeepD-60B (1M) (striped bars) across diverse annotation categories including CDS, intron, UTR, CRE, and TE families. Line plot with diamond markers shows the optimal extraction layer per annotation category, with numbers shown on the right y-axis. (E) Decoding of maize genomic features with routing weights with DeepD-60B (1M). Predicted probability tracks (purple) alongside reference annotations (blue boxes) are shown across the *tb1* region. The *Hopscotch* LTR/Copia retrotransposon insertion is indicated by a light green box.

In *E. coli*, classifiers trained on DeepD-60B (8K) routing weights generated probability tracks that resolved annotated elements at high positional resolution (**Figure 5B**). At a CRISPR array locus, the predicted CRISPR repeat and spacer signals alternated along the array and matched the annotated repeat–spacer pattern. This indicates that DeepD routing encodes the internal periodic structure of the array, rather than merely detecting repetitive sequence. At the *lac* operon, gene-associated probabilities aligned with the boundaries of *lacA*, *lacY*, *lacZ*, and *lacI*, with reduced signal in adjacent intergenic regions. These results indicate that sparse routing contains fine-scale information about discrete prokaryotic genomic elements.

We next examined whether the same routing representation resolves coding and repetitive organization in the human genome. At the *HBB* gene cluster on chromosome 11, routing-based classifiers recovered CDS signals at the annotated *HBB* and *HBD* genes and produced class-specific tracks for LTR, LINE, and SINE elements that aligned with their annotated boundaries (**Figure 5C** and **S4B**), indicating that the routing representation distinguishes major repeat classes rather than generating a generic repeat signal. Genome-wide probing further confirmed high decoding performance across human CDS and major repeat categories, with a mean area under the precision–recall curve (AUPRC) of 0.92 and particularly strong performance for CDS and SINE elements (AUPRC = 0.96 and 0.95, respectively) (**Figure S4A**). Models with 8K and 1M contexts showed similarly high AUPRCs for these compact genomic features, suggesting that coding boundaries and major repeat classes are already robustly encoded in local-to-intermediate routing representations. The best-performing layers differed among annotation classes, with CDS features peaking at deeper layers and repeat classes generally decoded from middle-to-deep layers. Thus, in the human genome, sparse routing provides an interpretable representation of both gene structure and repeat architecture, with layer-specific organization of genomic features.

We then tested DeepD in maize, a repeat-rich genome in which more than 85% of the sequence is repetitive and many CREs act over tens to hundreds of kilobases.^45^ Compared with the compact human CDS and repeat features, maize provides a more stringent test for long-context routing. Across gene structure annotations, abundant transposable element (TE) families, and CREs, DeepD-60B (1M) showed stronger decoding performance than the 8K model, consistent with the long-range context required for distal regulatory elements (**Figure 5D**). As in human, maize features were best decoded at different depths, with coding features peaking in deeper layers and repeats generally decoded from middle-to-deep layers.

At the maize *tb1* locus, a major domestication gene that suppresses axillary branch growth and promotes apical dominance,^46,47^ the routing-based CDS track accurately recovered the annotated gene region (**Figure 5E** and **S5A**). It also identified a prominent signal approximately 60 kb upstream, coinciding with the *Hopscotch* LTR/Copia retrotransposon, a functionally validated enhancer that increases *tb1* expression and contributed to the reduced branching architecture of domesticated maize.^47^ At *gt1*, another domestication gene that restricts axillary growth and reduces ear prolificacy,^48^ DeepD similarly recovered the annotated intron, CDS, UTR, CRE, and TE elements (**Figure S5B**). Together, these examples show that DeepD routing features capture not only canonical gene structures but also known distal functional elements associated with maize domestication.

### Controllable design of maize core promoters with specified strength

Core promoter sequences encode regulatory features that determine the magnitude of transcriptional output. Promoter strength depends on the identity, position, and spacing of multiple sequence motifs, making the generation of promoters with predefined activity a stringent test of whether a genomic model has learned regulatory grammar rather than memorized natural sequences. Recent generative approaches, including generative adversarial networks, autoregressive language models, and diffusion models, have enabled the design of synthetic regulatory DNA with specified activity levels, some of which has been experimentally validated.^20,22,25^ We therefore asked whether DeepD could generate novel maize core promoters conditioned for high or low transcriptional activity.

We fine-tuned DeepD using maize core promoters assigned Low, Med, and High activity labels based on STARR-seq data in maize protoplasts (**Figure 6A** and **S6A,** see **Methods**).^49^ Using these labels as prompts, we conditioned DeepD to generate 170-bp synthetic core promoters (SCPs) corresponding to High and Low activity classes (**Figure 6B**). The SCPs showed low similarity to maize sequences, suggesting that DeepD generates novel sequences rather than memorizing natural sequences (**Figure S6B**). We further evaluated the SCPs using a separately trained regression model based on DeepD sequence embeddings. Predicted promoter activity followed the intended conditioning direction, and High and Low SCPs formed distinct distributions in the embedding space (**Figure S6C**), indicating that the conditioning labels were translated into sequence features recognized by the activity predictor.

**Figure 6.**
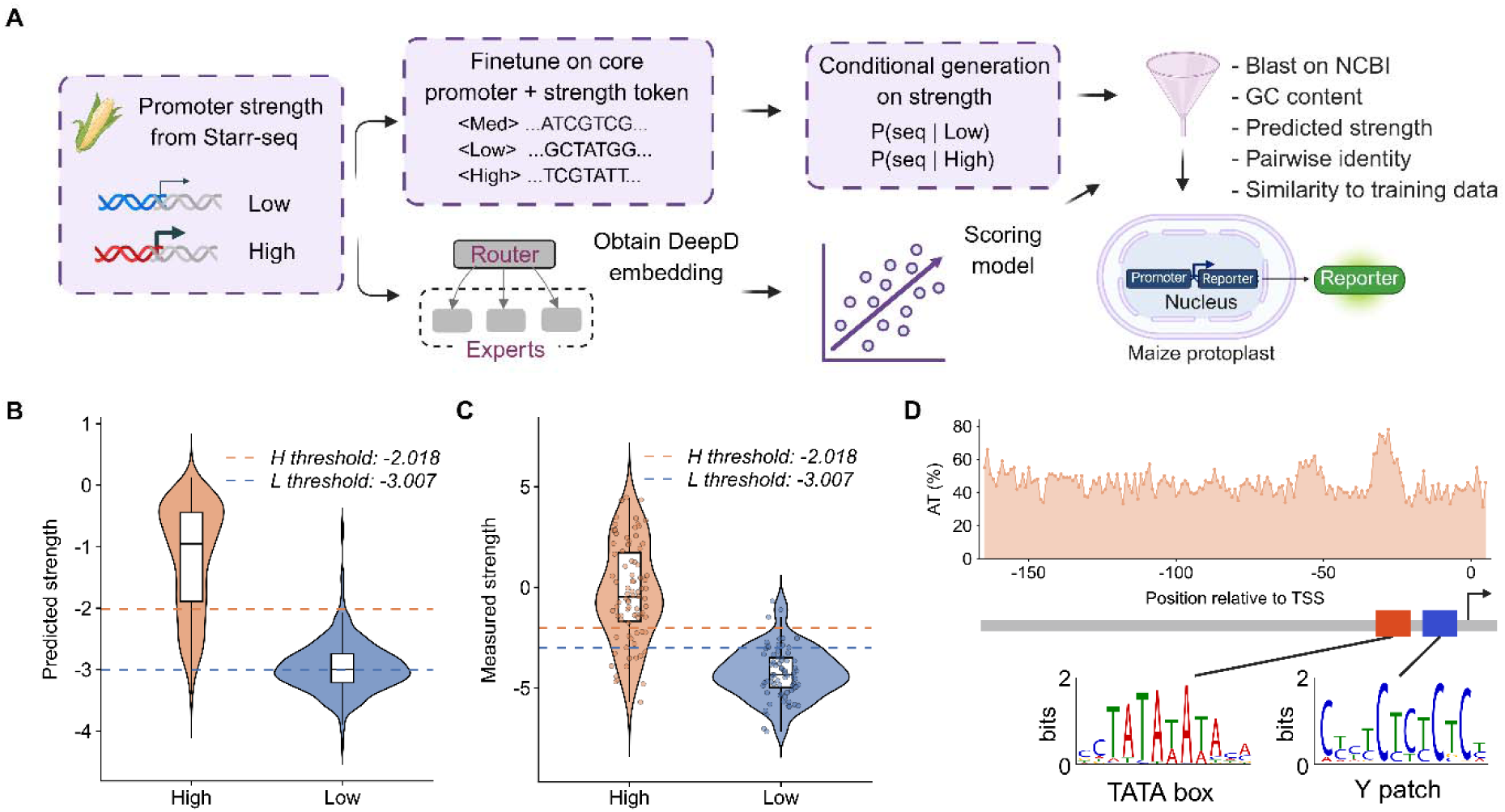
DeepD enables controllable design of functional maize core promoters. (A) Pipeline for the conditional generation and multi-step computational filtering of maize core promoters using DeepD. The model was fine-tuned on enhancerless STARR-seq data (−165 to +5 bp relative to the transcription start site, TSS) from maize protoplasts. Filtering criteria included BLAST against the NCBI nt database, identity × coverage < 0.6, full-length homology < 80%, GC content between 40% and 65%, predicted promoter strength, pairwise candidate similarity, and similarity to training sequences. (B) Predicted promoter strength distributions of the conditionally generated High (n = 450) and Low (n = 450) candidate sequence pools (Wilcoxon rank-sum test, P < 2.2 × 10^-16^). Dashed lines indicate the High (−2.018) and Low (−3.007) activity thresholds. (C) In vivo validation using dual-luciferase reporter assays of the generated sequences in maize protoplasts. Box plots show the measured promoter strength (log2-transformed Luc/Ren ratio normalized to the 35Smini promoter) for the evaluated High (n = 83) and Low (n = 71) candidates. Data points represent the mean of three biological replicates (Wilcoxon rank-sum test, P < 2.2 × 10^-16^). Dashed lines represent the High (−2.018) and Low (−3.007) baselines established from the native dataset. (D) Positional distribution of AT content (upper) and representative enriched plant regulatory motifs across the selected High candidates (n = 100).

We next experimentally evaluated the activity of these SCPs using transient dual-luciferase reporter assays in maize protoplasts. The measured promoter activities were broadly consistent with the design direction, with High-conditioned SCPs driving stronger reporter expression than Low-conditioned SCPs (**Figure 6C**). Using the thresholds that defined the native High and Low conditioning labels as baselines, about 76% of the evaluated High SCPs exhibited activity above the High-activity threshold, whereas approximately 83% of the evaluated Low SCPs fell below the Low-activity threshold (**Figure 6C**). Sequence analysis identified canonical core promoter motifs in High SCPs, including the TATA box and Y patch (see **Methods** and **Figure 6D**). The positions of these motifs in SCPs were similar to those observed in natural core promoters.^49^

Together, these results demonstrate that DeepD can translate coarse activity labels into experimentally measurable differences in promoter strength. The generated sequences are distinct from natural maize promoters yet retain recognizable plant regulatory grammar, supporting the use of DeepD for controllable de novo design of functional regulatory elements.

### Chromosome-scale sequence generation in yeast

The design of synthetic eukaryotic chromosomes provides a powerful route for investigating genome organization and constructing programmable cellular systems. The Sc2.0 project established that yeast chromosomes can be extensively refactored according to predefined engineering principles while retaining cellular viability.^31,50^ Genome language models provide a data-driven alternative and have generated chromosome-scale yeast sequences; however, current prompt-based generation incompletely recapitulates native chromosome architecture, including gene and tRNA content, and provides limited control over chromosome-wide gene organization.^3^ Such limitations are particularly important for eukaryotic chromosomes, where coding, regulatory, and repetitive sequences must remain coordinated over long distances.^32^ Inspired by conditional infilling and recurrent context strategies in natural language modeling,^33,34^ we developed a Context–Anchor–Body (CAB) strategy in which downstream Context and upstream Anchor sequences condition the generation of an intervening Body segment (**Figure 7A**). During inference, DeepD generated fixed 8-kb Body chunks through an iterative sliding-window relay, using each newly generated segment to anchor the next round. This strategy decoupled final output length from the training context window and enabled full-length chromosome generation while maintaining local continuity and broad chromosomal organization.

**Figure 7.**
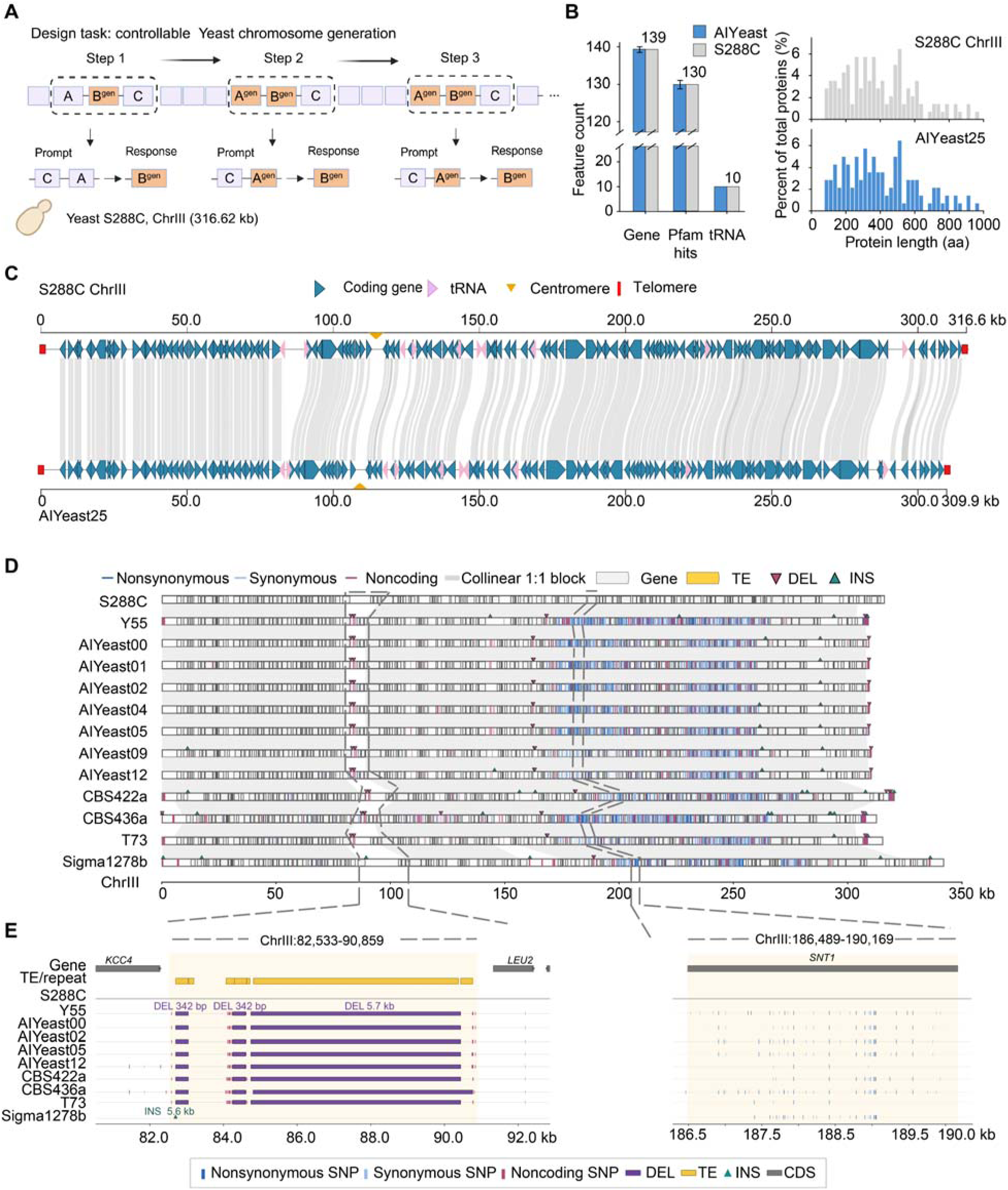
DeepD generates coherent eukaryotic chromosomes with natural-like population diversity. (A) Context–Anchor–Body (CAB) strategy for controllable chromosome-scale generation. DeepD iteratively generates 8-kb Body segments conditioned on upstream Anchor and downstream Context sequences, enabling long-sequence generation beyond a single model window. (B) The left panel shows generated *S. cerevisiae* ChrIII sequences preserve reference-like numbers of coding genes, Pfam-supported genes, and tRNAs. The right panel shows protein-length distributions for the S288C ChrIII reference (gray) and a representative DeepD-generated chromosome, AIYeast25 (blue). Protein lengths are shown as the percentage of all encoded proteins in each sequence. (C) Whole-chromosome alignment between S288C ChrIII and a representative generated chromosome shows extensive gene-level collinearity, indicating preservation of chromosome-scale continuity and gene organization. (D) Variant landscapes of generated and natural ChrIII sequences relative to S288C, showing synonymous, nonsynonymous, and noncoding variants across collinear 1:1 blocks. Generated chromosomes display population-like variant distributions rather than collapsing into a single artificial cluster. Y55, T73, CBS422a, CBS436a, and Sigma1278b are natural accessions. (E) Enlarged view of the variable region at ChrIII:82,533–90,859 bp. A transposable element present in S288C is absent from the generated chromosomes, matching its absence from 54.99% of natural pangenome accessions. (F) Enlarged view of the coding region at ChrIII:186,489–190,169 bp. The approximately 3.5-kb gene contains both synonymous and nonsynonymous substitutions in generated chromosomes, indicating that DeepD samples coding-sequence diversity observed among natural yeast isolates.

To test whether DeepD could generate S288C chromosome III (ChrIII)-scale variants that preserve coherent chromosome organization while closely mimicking natural yeast pangenomic diversity,^51^ we fine-tuned DeepD-60B at a 32,768-token context length on 762 high-quality *S. cerevisiae* genomes representing species-wide pangenomic diversity,^52^ with the S288C genome explicitly excluded from the fine-tuning set (see **Methods**). Using S288C ChrIII held-out backbone, DeepD generated 29 full-length ChrIII sequences, ranging from 309.86 to 316.62 kb in length, which is close to the 316.62-kb S288C reference. The generated chromosomes closely matched genomic features of S288C, including coding-gene number (138 to 140 versus 139 in S288C), tRNA number (10 versus 10 in S288C), protein-length distribution (median range from 392.50 to 400.50 aa versus 392.60 aa in S288C), and the number of genes with Pfam hits (129 to 131 versus 130 in S288C) (see **Methods** and **Figure 7B**). Notably, natural *S. cerevisiae* isolates are highly similar at the gene level. For example, T73 from the Wine/European clade and S288C from the M3 clade^53^ showed a median protein-level similarity of 99.9%, with 99.60% and 99.70% similarity in upstream and downstream flanks, respectively (**Figure S7A**). DeepD-generated ChrIII sequences were slightly more diverged from S288C, with an average gene-body protein similarity of 96.8% (**Figure S7A**). Whole-chromosome alignment revealed extensive gene-level collinearity: of the 140 genes in AIYeast25, 138 were collinear with their counterparts in S288C (**Figure 7C** and **S8**). Thus, DeepD coherently generated chromosome-scale yeast sequences, preserving the great majority of gene structure and gene order.

We next used comparative genomics to examine whether variations introduced in generated chromosomes could reflect yeast pangenomic diversity. We mapped 29 generated sequences against S288C with AnchorWave,^54^ and discovered eight to ten structural variants (SVs) and 3,288 to 4,025 single nucleotide polymorphisms (SNPs) per generated sequence (see **Methods**). The variant-density landscape was similar to that of the natural yeast population. For example, AIYeast25 contained 3,490 SNPs, of which 3,490 (100.00%) were present in at least one natural accession. Among all AIYeast25 SNPs, 1,511 were nonsynonymous, none of which were stop-gain SNPs. Of the eight SVs in AIYeast25, seven (87.50%) did not overlap annotated genes. Five deletions and two insertions were supported by at least one natural accession, and only one deletion was absent from all natural accessions. Phylogenetic analysis based on synonymous SNPs placed generated chromosomes across different yeast clades rather than in a single artificial cluster, supporting recapitulation of population-level diversity (**Figure S7B**).

We further closely investigated the nature of variations in generated sequences. Near 84 kb, a 5,958-bp S288C TE was absent in all generated ChrIIIs, consistent with its absence from 54.99% of natural accessions (**Figure 7E**). Around 150 kb, a ∼5-kb non-coding region showed variable SNPs and insertion patterns, consistent with natural diversity at this locus (**Figure S7C**). Lastly, a ∼3.5-kb gene near 190 kb carried 2–10 synonymous and 2–15 missense substitutions relative to S288C, but no stop-gain mutations, 94.12% SNPs appear in natural population, indicating the generation function of DeepD could largely reflect genetic diversity in genic regions (**Figure 7F**). Together, these results show that DeepD preserves chromosome-scale organization while introducing variants that mirror natural yeast pangenomic diversity.

### Controllable generation to megabase-scale plant genomes

Having established that CAB-based generation produces coherent chromosome-scale *S. cerevisiae* chromosomes, we next tested this strategy in a more complex multicellular eukaryotic genome. Higher eukaryotic genomes harbor longer intergenic regions and more complex variations, making long-context generation more challenging. We therefore applied CAB to a 1-Mb interval on chromosome 1 (0.13–1.13 Mb) and generated three independent copies (see **Methods** and **Figure 8A**). As a result, of the 299 reference protein-coding genes annotated in the corresponding TAIR12 segment, 245 were classified as core genes according to the *A. thaliana* pangenome analysis.^55^ Among these core genes, 243 were recovered in one of the generated sequences AIAtRegion1_01, corresponding to a recovery rate of 99.2% (**Figure 8B**). AIAtRegion1_01 recovered 296 of the 299 reference protein-coding genes (see **Methods**). Among the three not fully recovered genes, two of them had disrupted open reading frames, the third one was depleted due to its location at the very end of the segment. In addition, AIAtRegion1_01 contained a 225-bp gene fragment insertion, corresponding to part of the CDS of *AT1G04625*, located at approximately 1.29 Mb on chromosome 1 in TAIR12. The other two independent generations displayed similar patterns (**Figure S9A**). Moreover, AIAtRegion1_01 contained 266 versus 269 Pfam-annotated proteins in TAIR12, 1,753 versus 1,760 exons, and the same number of tRNA genes. Protein-length distributions were also highly similar between the TAIR12 and AIAtRegion1_01 (**Figure 8D**).

**Figure 8.**
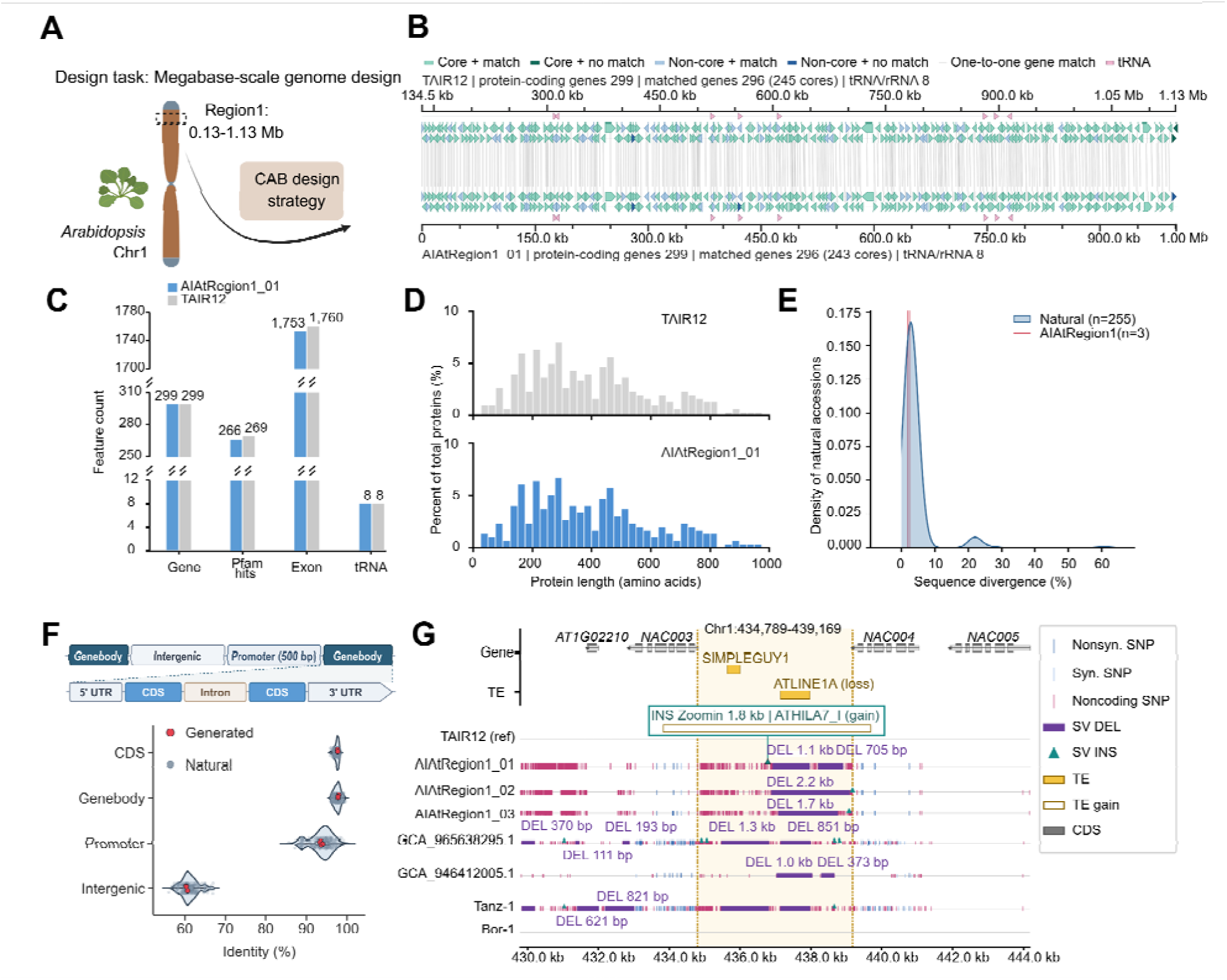
Megabase-scale *Arabidopsis* genome design by DeepD. (A) Schematic of CAB-based long-context generation of a 1-Mb region on *Arabidopsis* chromosome 1. Three independent sequences were generated from the same genomic interval. (B) Comparison of gene organization between TAIR12 and the representative generated sequence AIAtRegion1_01. Core protein-coding genes, CDS structures, unmatched genes, and newly introduced gene fragments are indicated. The 225-bp insertion near 550 kb corresponds to the first CDS of *AT1G04625*, located at ∼1.29 Mb on chromosome 1 in TAIR12. (C) Comparison of genomic composition between TAIR12 and AIAtRegion1_01, including gene number, Pfam-annotated proteins, CDS features, and tRNA genes. (D) Protein-length distributions in TAIR12 and AIAtRegion1_01. (E) Sequence divergence from TAIR12 for the corresponding 1-Mb interval across 255 natural *A. thaliana* accessions and three independently generated sequences. (F) Sequence conservation of CDSs, gene bodies, 500 bp promoter regions, and intergenic regions in natural accessions and DeepD-generated sequences relative to TAIR12. (G) Representative SVs in AIAtRegion1_01. The generated sequence contains an ∼1.8-kb fragment derived from the ∼10-kb Ty3-type LTR retrotransposon ATHILA7, while lacking part of bases in ∼1-kb Harbinger-type transposon SIMPLEGUY1 and ∼2.2-kb LINE element ATLINE1A. Gene and TE annotations surrounding the locus are shown. GCA 965638295.1, GCA 946412005.1, Tanz-1, and Bor-1 are natural accessions.

We then compared the extent of sequence divergence relative to TAIR12, between 255 natural *A. thaliana* accessions and the three generated copies (see **Methods**). The level of sequence divergence of the three generated sequences fell in the middle of observed divergence (**Figure 8E**). In addition, DeepD-generated variation is structured by genomic functional constraint rather than distributed uniformly across the sequence. As a result, sequence conservation across different functional categories could recapitulate the pattern observed in natural genomes (**Figure 8F**). CDSs were the most conserved, followed by gene bodies and promoter regions, whereas intergenic sequences showed substantially greater divergence.

We next examined the composition and potential source of newly-introduced variants in generated sequences. Joint variant calling with 255 natural accessions produced a population catalogue containing 5,493 synonymous, 10,141 nonsynonymous, and 48,332 noncoding variants. AIAtRegion1_01 carried 5,061 SNPs (**Figure S9B** and **S9C**), including 178 synonymous (3.52%), 427 nonsynonymous (8.44%), and 4,456 noncoding variants (88.05%). Among these SNPs, 1,985 (39.22%) were present in at least one natural accession, whereas 3,076 (60.78%) were specific to AIAtRegion1_01. Among the nonsynonymous changes, all 15 stop-gain SNPs were AIAtRegion1_01-specific and affected four genes. In two of these genes, premature termination truncated more than 30% of the reference protein sequence, such that the resulting proteins could no longer be matched to their corresponding reference protein models. SVs were more strongly confined to noncoding genomic compartments. AIAtRegion1_01 carried 12 SVs, comprising five insertions and seven deletions, all located in intergenic or intronic regions. Nine SVs were specific to AIAtRegion1_01, whereas three deletions were supported by natural accessions. Together, the generated variants were dominated at noncoding regions and a large proportion of them were newly-introduced.

TE-associated loci further distinguished the recovery of standing variation from the generation of new local sequence configurations (**Figure 8G**). The ATLINE1A-associated deletion overlapped deletion alleles in 236 natural accessions, although their breakpoints and lengths varied, indicating extensive population-level variations of this TE locus. By contrast, no natural accession carried an insertion within ±100 bp of the ATHILA7_I-associated insertion breakpoint in AIAtRegion1_01. However, searches of the inserted sequence against TAIR12 and the natural accession genomes identified homologous sequences near 10 Mb on chromosome 5 in 240 accessions, with a mean identity of 0.89. The insertion therefore happened in a brand new location using sequences similar to an endogenous TE found elsewhere in the *Arabidopsis* genome, not inserted by completely unconstrained foreign sequence. Together, these results show that CAB-based long-context generation could be to megabase-scale plant genomic sequence generation, with coherent genic regions while introducing some extent of new insertions and deletions in intergenic regions.

## Discussion

Genome foundation models support sequence prediction, interpretation, and generation, yet modeling various genomes of high complexity remains constrained.^3–5,22^ We trained DeepD-60B model on 5.69 Tb DNA sequence, activating approximately 1 billion parameters per token, with active computation substantially reduced. Its 5.69 Tb pretraining corpus spans the all domains of life together with viruses and retains whole-genome genic and full-space intergenic sequences. DeepD supports broad sequence-to-function prediction, strong zero-shot variant-effect prediction, and transferable representations of intergenic DNA, while expert routing captures features ranging from exon–intron boundaries to transposable elements and distal CREs. DeepD has unique capability of integrating accurate sequence-to-function prediction with coherent chromosome-scale sequence generation.

The sparse MoE architecture enables DeepD to increase model capacity without activating the entire network for each nucleotide, an approach that has substantially improved computational efficiency in natural language models.^8–10^ DeepD extends this principle to single-nucleotide modeling of phylogenetically diverse genomes over a one-million-token context window. Explicit species label and phylogenetic representation capture lineage-associated nucleotide composition, codon usage, and coding constraints, while expert-routing patterns provide interpretable representations of genomic features. In addition to exon–intron boundaries previously identified by dense genome models,^3^ DeepD interprets DNA transposons, LTR retrotransposons, distal CREs, and CRISPR repeat–spacer units. Thus, sparse routing provides both computational efficiency and a biologically informative organization of model capacity.

DeepD is also distinct in terms of data composition during pretraining. Rather than selecting gene-enriched regions, DeepD uses all genic sequences and full-space intergenic regions. Soft-masked repetitive bases remain in their native sequence context, although their contribution to the training loss is reduced. This strategy preserves the continuity and natural composition of whole genomes while preventing abundant repeats from dominating optimization. Consistent with this design, DeepD achieves broad zero-shot performance with its exceptional strengths in regulatory and non-coding tasks. Its routing representations capture TEs and distal CREs, extending beyond the primarily gene-centered features learned by many existing models.^3,4,6^ The generation of a TE-derived insertion within an otherwise collinear *Arabidopsis* region further suggests that genome-wide pretraining enables SV in intergenic sequences rather than arbitrary sequence insertion.

For biological and clinical applications in human, DeepD and AlphaGenome can be ideally complementary to each other. Across benchmarks of localized variant effects—including both SNVs and non-SNVs—DeepD was competitive against currently published Evo-series and NT models, and outperformed AlphaGenome on *BRCA1*, *BRCA2*, ClinVar, and protein-fitness prediction, with particular strength for synonymous, splice-related, and non-coding variants. In supervised human regulatory tasks, DeepD ranked first among general genome language models, indicating strong representations of intergenic DNA. AlphaGenome nevertheless retained advantages on several task-aligned supervised outputs, particularly TF-binding and splice-site prediction, for which extensive human functional genomic data were used for training. Notably, comparisons involving AlphaGenome on SpliceVarDB and supervised regulatory tasks should be interpreted in light of its direct training on closely aligned functional genomic tracks, potentially providing somewhat overestimation of its performance. Nevertheless, owning to its strong zero-shot variant-effect prediction and transferable representation for intergenic sequences, DeepD will be one of the top choices for sequence-to-function prediction of many other species with limited functional genomic data.

Our CAB strategy addresses a major limitation of existing genome generators. Previous models have generated long genomic sequences mainly through unconstrained autoregressive continuation from an initial prompt.^3,5,27^ As generation further extends, earlier sequence eventually falls outside the available model context and can no longer accurately guide subsequent output, limiting control over a long stretch of genomic organization. Using CAB, DeepD generated entire yeast chromosomes coherently that can hardly be distinguished from sequences of the natural yeast population. Similarly, generated megabase-scale *Arabidopsis* segments preserve coherent genic regions while introducing some extent of new insertions and deletions in intergenic regions. CAB therefore advances genome-scale generation from open-ended sequence continuation to controllable chromosome construction. Furthermore, since CAB decouples total output length from the fixed generation window through iterative sequence extension, DeepD therefore provides a reasonable path towards ultimately genome-scale sequence design.

Taken together, comparison with current published models positions DeepD as a leading genome foundation model that unifies sequence-to-function prediction with coherent genome-scale generation. DeepD combines strong predictive performance across protein-coding, regulatory, and non-coding sequences with the capacity to generate functional regulatory elements and coherent chromosome-scale sequences. This integration moves genome modeling beyond task-specific prediction or unconstrained sequence generation toward a unified framework for interpreting natural variation and designing genomes with specified properties.

### Limitations of the study

Although DeepD advances substantially in unifying accurate sequence-to-function prediction with long-context coherent genomic sequence generation, several limitations remain. First, DeepD primarily models DNA sequences without explicitly incorporating other potentially related modalities, future incorporation of functional genomics, epigenomics, transcriptomics, proteomics, three-dimensional genome organization could further enhance DeepD into a more comprehensive foundation model for predictive and generative biology. Second, for sequence generation, DeepD generates coherent yeast chromosomes and relatively gene-rich segments of *Aradopisis*, its applicability to more complex, repeat-rich regions—particularly those containing long and nested retrotransposon insertions—remains untested and may require additional longer-context fine-tuning and optimization of CAB generation parameters. Finally, although DeepD has the capacity in generating chromosome-scale sequences that can hardly be distinguished from that of the natural population, the functional performance of the generated sequences has not been extensively tested experimentally, awaiting further confirmation in specific designing projects in the future.

## Lead contact

Requests for resources and further information should be addressed to the lead contact, Jinsheng Lai.

## Code and data availability

Custom analysis scripts, 29 generated *S. cerevisiae* chrIII sequences, and three generated megabase-scale *A. thaliana* genomic sequences are currently available at https://github.com/biomap-research/DeepD.git. During peer review, DeepD will be available to editors and reviewers through anonymous APIs. The source code, model weights, and inference pipeline will be released through the same Github repository no later than the time of publication.

## Supporting information

Method Details

## Acknowledgements

This work was supported by the National Natural Science Foundation of China, the National Key Research and Development Program of China. Thanks to High-performance Computing Platform of China Agricultural University for its support of large-scale computation.

## Author contributions

J.L. and H.Z. conceived the project. B.X., Q.Y., J.L. and H.Z. supervised the study. B.X., Y.G. and Q.Y. designed the DeepD framework and model architecture. Y.G. and Q.Y. developed the training and inference code. Y.G. and Q.Y. contributed to model engineering, distributed training, and API deployment. Z.W., A.Y., Z.C. and C.Y. curated and processed the pretraining genome dataset, species labels, phylogenetic information, and genome annotations. Y.G. performed model pretraining, context extension, checkpoint evaluation, and scaling analyses. Z.C. performed phylogeny-informed modeling analyses. B.X. and C.Y. performed zero-shot and supervised sequence-to-function prediction analyses. Z.W. designed and performed expert-routing interpretability analyses. A.Y. and B.H. designed and performed *cis*-regulatory element generation and validation. M.Z. performed *cis*-regulatory element activity validation. B.X., Y.G., B.H. and Q.Y. designed the chromosome-scale generation framework. Z.W. and Y.G. generated and analyzed yeast ChrIII-scale sequences. Z.W. and Y.G. generated and analyzed megabase-scale *Arabidopsis* genomic sequences. N.D., F.Y., J.C., Q.L., H.Z. and W.S. assisted with manuscript positioning and conceptual improvement. B.X., Z.W., A.Y., Z.C., and Y.G. wrote the first draft. All authors revised and approved the final manuscript.

## METHODS

Detailed experimental and computational procedures are provided in the METHODS DETAILS including the following:

- DeepD model architecture and training
- DeepD pretraining data
- Sequence-to-function prediction evaluations
- Analysis of phylogeny-informed genome modeling
- Biological interpretation of expert routing
- Controllable design and validation of maize core promoters
- Controllable long-context design of eukaryotic chromosomes

## References

1. Benegas, G., Albors, C., Aw, A.J., Ye, C., and Song, Y.S. (2025). A DNA language model based on multispecies alignment predicts the effects of genome-wide variants. Nature Biotechnology 43, 1960–1965. 10.1038/s41587-024-02511-w.

2. Benegas, G., Batra, S.S., and Song, Y.S. (2023). DNA language models are powerful predictors of genome-wide variant effects. Proceedings of the National Academy of Sciences 120, e2311219120. 10.1073/pnas.2311219120.

3. Brixi, G., Durrant, M.G., Ku, J., Naghipourfar, M., Poli, M., Sun, G., Brockman, G., Chang, D., Fanton, A., Gonzalez, G.A., et al. (2026). Genome modelling and design across all domains of life with Evo 2. Nature 652, 1349–1361. 10.1038/s41586-026-10176-5.

4. Dalla-Torre, H., Gonzalez, L., Mendoza-Revilla, J., Lopez Carranza, N., Grzywaczewski, A.H., Oteri, F., Dallago, C., Trop, E., de Almeida, B.P., Sirelkhatim, H., et al. (2025). Nucleotide Transformer: building and evaluating robust foundation models for human genomics. Nature Methods 22, 287–297. 10.1038/s41592-024-02523-z.

5. Nguyen, E., Poli, M., Durrant, M.G., Kang, B., Katrekar, D., Li, D.B., Bartie, L.J., Thomas, A.W., King, S.H., Brixi, G., et al. (2024). Sequence modeling and design from molecular to genome scale with Evo. Science 386, eado9336. 10.1126/science.ado9336.

6. Zhai, J., Gokaslan, A., Schiff, Y., Berthel, A., Liu, Z.-Y., Lai, W.-Y., Miller, Z.R., Scheben, A., Stitzer, M.C., Romay, M.C., et al. (2025). Cross-species modeling of plant genomes at single-nucleotide resolution using a pretrained DNA language model. Proceedings of the National Academy of Sciences 122, e2421738122. 10.1073/pnas.2421738122.

7. Avsec, Ž., Latysheva, N., Cheng, J., Novati, G., Taylor, K.R., Ward, T., Bycroft, C., Nicolaisen, L., Arvaniti, E., Pan, J., et al. (2026). Advancing regulatory variant effect prediction with AlphaGenome. Nature 649, 1206–1218. 10.1038/s41586-025-10014-0.

8. Dai, D., Deng, C., Zhao, C., Xu, R., Gao, H., Chen, D., Li, J., Zeng, W., Yu, X., and Wu, Y. (2024). Deepseekmoe: Towards ultimate expert specialization in mixture-of-experts language models. pp. 1280–1297.

9. Guo, D., Yang, D., Zhang, H., Song, J., Wang, P., Zhu, Q., Xu, R., Zhang, R., Ma, S., Bi, X., et al. (2025). DeepSeek-R1 incentivizes reasoning in LLMs through reinforcement learning. Nature 645, 633–638. 10.1038/s41586-025-09422-z.

10. Fedus, W., Zoph, B., and Shazeer, N. (2022). Switch transformers: Scaling to trillion parameter models with simple and efficient sparsity. Journal of Machine Learning Research 23, 1–39.

11. Shazeer, N., Mirhoseini, A., Maziarz, K., Davis, A., Le, Q., Hinton, G., and Dean, J. (2017). Outrageously large neural networks: The sparsely-gated mixture-of-experts layer.

12. Ashish, V. (2017). Attention is all you need. Advances in neural information processing systems.

13. Brown, T., Mann, B., Ryder, N., Subbiah, M., Kaplan, J.D., Dhariwal, P., Neelakantan, A., Shyam, P., Sastry, G., and Askell, A. (2020). Language models are few-shot learners. Advances in neural information processing systems 33, 1877–1901.

14. Hoffmann, J., Borgeaud, S., Mensch, A., Buchatskaya, E., Cai, T., Rutherford, E., de Las Casas, D., Hendricks, L.A., Welbl, J., and Clark, A. (2022). An empirical analysis of compute-optimal large language model training. Advances in neural information processing systems 35, 30016–30030.

15. Mendoza-Revilla, J., Trop, E., Gonzalez, L., Roller, M., Dalla-Torre, H., de Almeida, B.P., Richard, G., Caton, J., Lopez Carranza, N., Skwark, M., et al. (2024). A foundational large language model for edible plant genomes. Communications Biology 7, 835. 10.1038/s42003-024-06465-2.

16. Wang, A., Li, J., Dong, H., Xu, B., Yin, Q., Xu, Y., Fu, J., and Zhao, J. (2025). Omnireg-gpt: a high-efficiency foundation model for comprehensive genomic sequence understanding. Nature Communications 16, 10139. 10.1038/s41467-025-65066-7.

17. Aspidova, A., Kuratov, Y., Shadskiy, A., Burtsev, M., and Fishman, V. (2026). Back to BERT in 2026: ModernGENA as a Strong, Efficient Baseline for DNA Foundation Models. bioRxiv, 2026.2004. 2021.719816. 10.64898/2026.04.21.719816.

18. Chen, K., Zhou, Y., Ding, M., Wang, Y., Ren, Z., and Yang, Y. (2024). Self-supervised learning on millions of primary RNA sequences from 72 vertebrates improves sequence-based RNA splicing prediction. Briefings in Bioinformatics 25, bbae163. 10.1093/bib/bbae163.

19. Karollus, A., Hingerl, J., Gankin, D., Grosshauser, M., Klemon, K., and Gagneur, J. (2024). Species-aware DNA language models capture regulatory elements and their evolution. Genome Biology 25, 83. 10.1186/s13059-024-03221-x.

20. DaSilva, L.F., Senan, S., Kribelbauer-Swietek, J.F., Patel, Z.M., Louis, L.K., Reddy, A.J., Gabbita, S., Rosen, J.D., Nussbaum, Z., Córdova, C.M.V., et al. (2026). Designing synthetic regulatory elements using the generative AI framework DNA-Diffusion. Nature Genetics 58, 180–194. 10.1038/s41588-025-02441-6.

21. de Almeida, B.P., Schaub, C., Pagani, M., Secchia, S., Furlong, E.E.M., and Stark, A. (2024). Targeted design of synthetic enhancers for selected tissues in the Drosophila embryo. Nature 626, 207–211. 10.1038/s41586-023-06905-9.

22. Lal, A., Garfield, D., Biancalani, T., and Eraslan, G. (2024). Designing realistic regulatory DNA with autoregressive language models. Genome Research 34, 1411–1420. 10.1101/gr.279142.124.

23. Li, C., Zhang, Q., Chen, H., Lin, K., Luo, C., Yang, M., Xu, W., Yao, F., Yan, J., Yang, Q.-Y., and Hu, X. (2026). PlantGFM: A Genomic Foundation Model for Discovery and Creation of Plant Genes. Advanced Science 13, e75772. 10.1002/advs.75772.

24. Madani, A., Krause, B., Greene, E.R., Subramanian, S., Mohr, B.P., Holton, J.M., Olmos, J.L., Xiong, C., Sun, Z.Z., Socher, R., et al. (2023). Large language models generate functional protein sequences across diverse families. Nature Biotechnology 41, 1099–1106. 10.1038/s41587-022-01618-2.

25. Zrimec, J., Fu, X., Muhammad, A.S., Skrekas, C., Jauniskis, V., Speicher, N.K., Börlin, C.S., Verendel, V., Chehreghani, M.H., Dubhashi, D., et al. (2022). Controlling gene expression with deep generative design of regulatory DNA. Nature Communications 13, 5099. 10.1038/s41467-022-32818-8.

26. Faure, A.J., Domingo, J., Schmiedel, J.M., Hidalgo-Carcedo, C., Diss, G., and Lehner, B. (2022). Mapping the energetic and allosteric landscapes of protein binding domains. Nature 604, 175–183. 10.1038/s41586-022-04586-4.

27. Merchant, A.T., King, S.H., Nguyen, E., and Hie, B.L. (2026). Semantic design of functional de novo genes from a genomic language model. Nature 649, 749–758. 10.1038/s41586-025-09749-7.

28. Shao, B., and Yan, J. (2024). A long-context language model for deciphering and generating bacteriophage genomes. Nature Communications 15, 9392. 10.1038/s41467-024-53759-4.

29. Bengio, S., Vinyals, O., Jaitly, N., and Shazeer, N. (2015). Scheduled sampling for sequence prediction with recurrent neural networks. Advances in neural information processing systems 28.

30. Guan, J., Mao, X., Fan, C., Liu, Z., Ding, W., and Huang, M. (2021). Long text generation by modeling sentence-level and discourse-level coherence. pp. 6379–6393.

31. Annaluru, N., Muller, H., Mitchell, L.A., Ramalingam, S., Stracquadanio, G., Richardson, S.M., Dymond, J.S., Kuang, Z., Scheifele, L.Z., Cooper, E.M., et al. (2014). Total Synthesis of a Functional Designer Eukaryotic Chromosome. Science 344, 55–58. 10.1126/science.1249252.

32. Goold, H.D., Kroukamp, H., Erpf, P.E., Zhao, Y., Kelso, P., Calame, J., Timmins, J.J.B., Wightman, E.L.I., Peng, K., Carpenter, A.C., et al. (2025). Construction and iterative redesign of synXVI a 903 kb synthetic Saccharomyces cerevisiae chromosome. Nature Communications 16, 841. 10.1038/s41467-024-55318-3.

33. Bavarian, M., Jun, H., Tezak, N., Schulman, J., McLeavey, C., Tworek, J., and Chen, M. (2022). Efficient training of language models to fill in the middle. arXiv preprint arXiv:2207.14255.

34. Xiao, G., Tian, Y., Chen, B., Han, S., and Lewis, M. (2024). Efficient streaming language models with attention sinks. pp. 21875–21895.

35. Plotkin, J.B., and Kudla, G. (2011). Synonymous but not the same: the causes and consequences of codon bias. Nature Reviews Genetics 12, 32–42. 10.1038/nrg2899.

36. Nijkamp, E., Ruffolo, J.A., Weinstein, E.N., Naik, N., and Madani, A. (2023). ProGen2: Exploring the boundaries of protein language models. Cell Systems 14, 968–978.e963. 10.1016/j.cels.2023.10.002.

37. Hayes, T., Rao, R., Akin, H., Sofroniew, N.J., Oktay, D., Lin, Z., Verkuil, R., Tran, V.Q., Deaton, J., Wiggert, M., et al. (2025). Simulating 500 million years of evolution with a language model. Science 387, 850–858. 10.1126/science.ads0018.

38. Rives, A., Meier, J., Sercu, T., Goyal, S., Lin, Z., Liu, J., Guo, D., Ott, M., Zitnick, C.L., Ma, J., and Fergus, R. (2021). Biological structure and function emerge from scaling unsupervised learning to 250 million protein sequences. Proceedings of the National Academy of Sciences 118, e2016239118. 10.1073/pnas.2016239118.

39. Penić, R.J., Vlašić, T., Huber, R.G., Wan, Y., and Šikić, M. (2025). RiNALMo: general-purpose RNA language models can generalize well on structure prediction tasks. Nature Communications 16, 5671. 10.1038/s41467-025-60872-5.

40. Findlay, G.M., Daza, R.M., Martin, B., Zhang, M.D., Leith, A.P., Gasperini, M., Janizek, J.D., Huang, X., Starita, L.M., and Shendure, J. (2018). Accurate classification of BRCA1 variants with saturation genome editing. Nature 562, 217–222. 10.1038/s41586-018-0461-z.

41. Huang, H., Hu, C., Na, J., Hart, S.N., Gnanaolivu, R.D., Abozaid, M., Rao, T., Tecleab, Y.A., Ambrosone, C.B., Yao, S., et al. (2025). Functional evaluation and clinical classification of BRCA2 variants. Nature 638, 528–537. 10.1038/s41586-024-08388-8.

42. Landrum, Melissa J., Chitipiralla, S., Kaur, K., Brown, G., Chen, C., Hart, J., Hoffman, D., Jang, W., Liu, C., Maddipatla, Z., et al. (2025). ClinVar: updates to support classifications of both germline and somatic variants. Nucleic Acids Research 53, D1313–D1321. 10.1093/nar/gkae1090.

43. Landrum, M.J., Lee, J.M., Benson, M., Brown, G.R., Chao, C., Chitipiralla, S., Gu, B., Hart, J., Hoffman, D., Jang, W., et al. (2018). ClinVar: improving access to variant interpretations and supporting evidence. Nucleic Acids Research 46, D1062–D1067. 10.1093/nar/gkx1153.

44. Sullivan, P.J., Quinn, J.M.W., Wu, W., Pinese, M., and Cowley, M.J. (2024). SpliceVarDB: A comprehensive database of experimentally validated human splicing variants. The American Journal of Human Genetics 111, 2164–2175. 10.1016/j.ajhg.2024.08.002.

45. Chen, J., Wang, Z., Tan, K., Huang, W., Shi, J., Li, T., Hu, J., Wang, K., Wang, C., Xin, B., et al. (2023). A complete telomere-to-telomere assembly of the maize genome. Nature Genetics 55, 1221–1231. 10.1038/s41588-023-01419-6.

46. Doebley, J., Stec, A., and Hubbard, L. (1997). The evolution of apical dominance in maize. Nature 386, 485–488. 10.1038/386485a0.

47. Studer, A., Zhao, Q., Ross-Ibarra, J., and Doebley, J. (2011). Identification of a functional transposon insertion in the maize domestication gene tb1. Nature Genetics 43, 1160–1163. 10.1038/ng.942.

48. Wills, D.M., Whipple, C.J., Takuno, S., Kursel, L.E., Shannon, L.M., Ross-Ibarra, J., and Doebley, J.F. (2013). From many, one: genetic control of prolificacy during maize domestication. PLoS Genet 9, e1003604. 10.1371/journal.pgen.1003604.

49. Jores, T., Tonnies, J., Wrightsman, T., Buckler, E.S., Cuperus, J.T., Fields, S., and Queitsch, C. (2021). Synthetic promoter designs enabled by a comprehensive analysis of plant core promoters. Nature Plants 7, 842–855. 10.1038/s41477-021-00932-y.

50. Richardson, S.M., Mitchell, L.A., Stracquadanio, G., Yang, K., Dymond, J.S., DiCarlo, J.E., Lee, D., Huang, C.L.V., Chandrasegaran, S., Cai, Y., et al. (2017). Design of a synthetic yeast genome. Science 355, 1040–1044. 10.1126/science.aaf4557.

51. Peter, J., De Chiara, M., Friedrich, A., Yue, J.-X., Pflieger, D., Bergström, A., Sigwalt, A., Barre, B., Freel, K., Llored, A., et al. (2018). Genome evolution across 1,011 Saccharomyces cerevisiae isolates. Nature 556, 339–344. 10.1038/s41586-018-0030-5.

52. Loegler, V., Thiele, P., Teyssonnière, E., Tsouris, A., Brach, G., Cruaud, C., Payen, E., Engelen, S., Dunham, M.J., Hou, J., et al. (2025). From genotype to phenotype with 1,086 near telomere-to-telomere yeast genomes. Nature 648, 649–658. 10.1038/s41586-025-09637-0.

53. Esberg, A., Muller, L.A.H., and McCusker, J.H. (2011). Genomic Structure of and Genome-Wide Recombination in the Saccharomyces cerevisiae S288C Progenitor Isolate EM93. PLOS ONE 6, e25211. 10.1371/journal.pone.0025211.

54. Song, B., Marco-Sola, S., Moreto, M., Johnson, L., Buckler, E.S., and Stitzer, M.C. (2022). AnchorWave: Sensitive alignment of genomes with high sequence diversity, extensive structural polymorphism, and whole-genome duplication. Proceedings of the National Academy of Sciences 119, e2113075119. 10.1073/pnas.2113075119.

55. Lian, Q., Huettel, B., Walkemeier, B., Mayjonade, B., Lopez-Roques, C., Gil, L., Roux, F., Schneeberger, K., and Mercier, R. (2024). A pan-genome of 69 Arabidopsis thaliana accessions reveals a conserved genome structure throughout the global species range. Nature Genetics 56, 982–991. 10.1038/s41588-024-01715-9.

