## Supplementary Figures for "DeepD enables efficient genome modeling and controllable long-context sequence generation across domains of life"

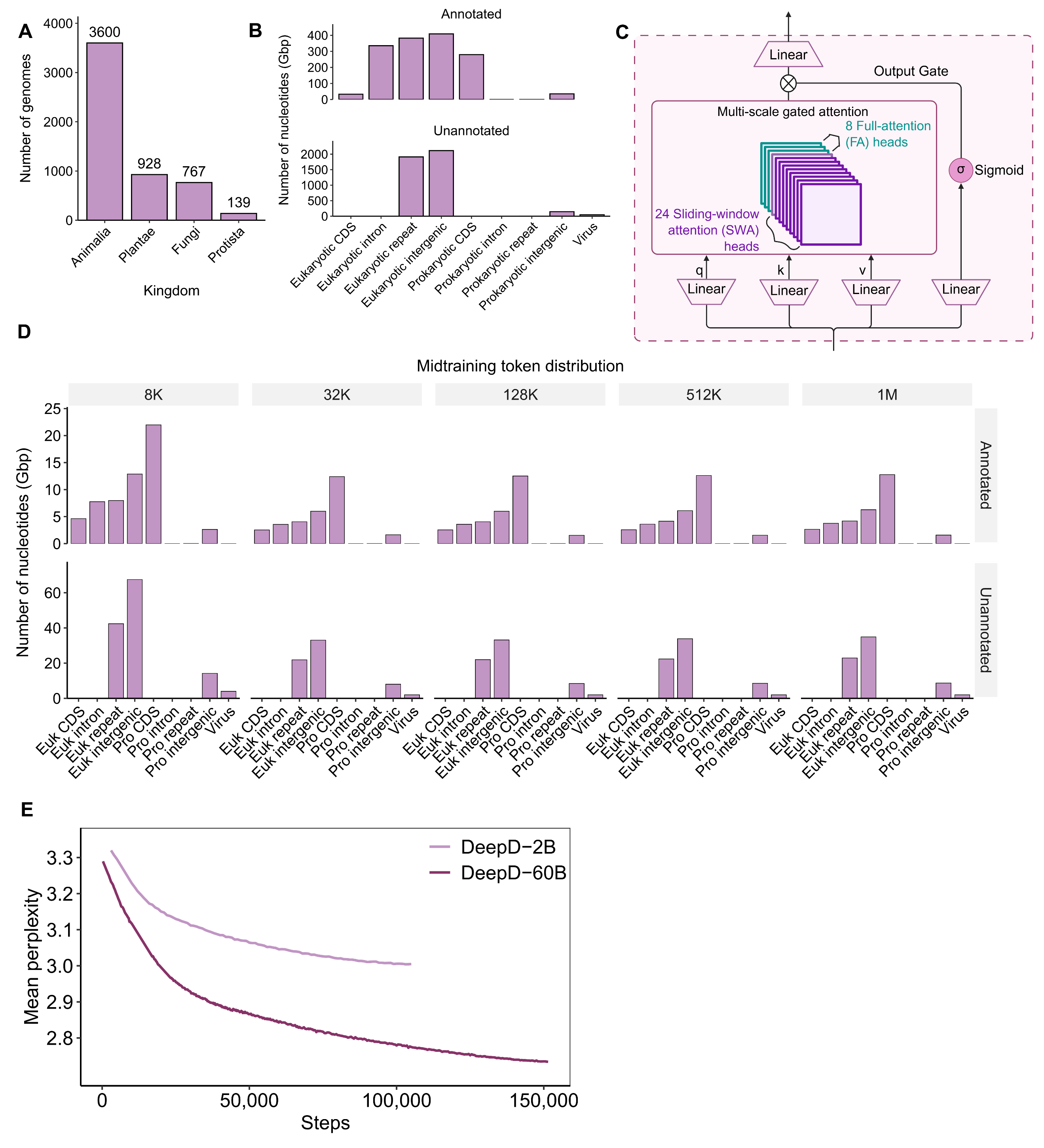

**Figure S1. Composition of the DeepD training corpus, attention design, midtraining token distribution, and pretraining dynamics.** (A) Taxonomic breakdown of the eukaryotic genomes included in the DeepD training corpus, showing representation from Animalia (3,600 genomes), Plantae (928 genomes), Fungi (767 genomes), and Protista (139 genomes). (B) Token distribution during pretraining in annotated and unannotated genomes. Annotated tokens include eukaryotic CDS, introns, repeats, and intergenic regions, as well as prokaryotic CDS and viral sequences. Unannotated tokens are dominated by eukaryotic repeats and eukaryotic intergenic regions, reflecting the native composition of genome-scale sequence space. (C) Schematic of the multi-scale gated attention module used in DeepD. Each block contains 24 sliding-window attention (SWA) heads and 8 full-attention (FA) heads, followed by a sigmoid-gated output projection, enabling simultaneous modeling of local sequence patterns and longer-range dependencies. (D) Token distribution during midtraining across progressively increasing context lengths (8K, 32K, 128K, 512K, and 1M). For each context stage, both annotated and unannotated tokens retain broad representation of CDS, introns, repeats, intergenic regions, and viral sequences, indicating that genomic diversity is preserved during long-context extension. (E) Mean perplexity during pretraining for DeepD-2B and DeepD-60B. Perplexity decreases steadily throughout training for both models, with the 60B model consistently achieving lower perplexity, indicating stable optimization and improved modeling capacity with scale.

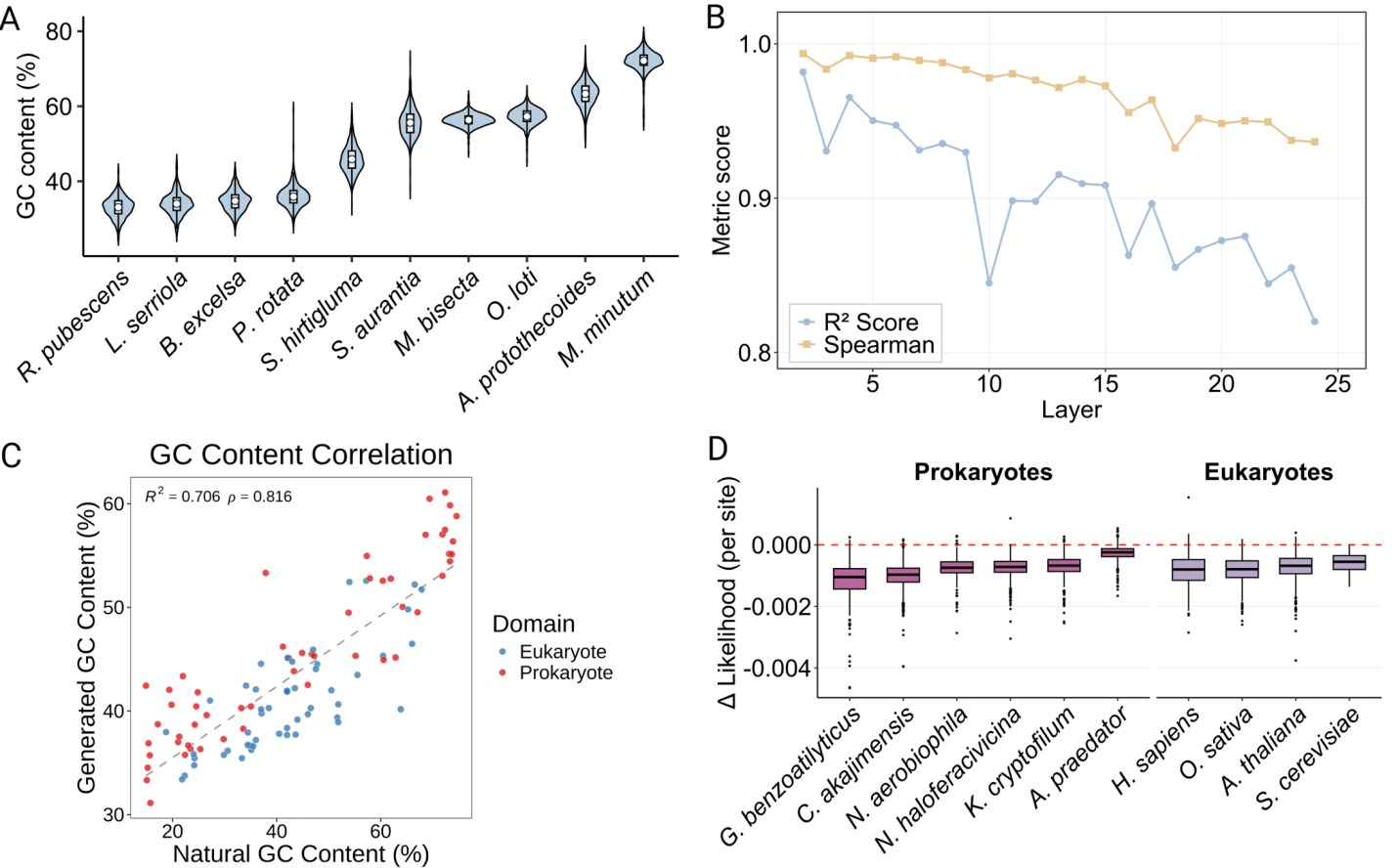

**Figure S2. Supplementary validation of phylogeny-guided sequence generation, interpretability, and structural probing.** (A) GC content distribution of representative species spanning a wide genomic spectrum. Violin plots representing the genomic GC content (%) distributions for the ten selected reference species, chosen to validate model adaptiveness across a broad compositional spectrum. White squares indicate the median, and internal boxplots show the interquartile range and whiskers. (B) Layer-wise performance of GC content prediction from expert activations. Line plot showing the prediction accuracy—measured by R^2^ score (blue circles) and Spearman correlation (yellow squares)—of Ridge regression models trained layer by layer (Layers 2 to 24) using expert activation weights to predict sequence-level GC content. (C) Correlation between natural and generated genomic GC content. Scatter plot comparing the natural GC content (%) with the GC content (%) of sequences generated by DeepD under species-specific taxonomic conditioning. Each data point represents a distinct species, color-coded by taxonomic domain (Eukaryote in blue, Prokaryote in red). The coefficient of determination and Spearman's rank correlation coefficient are indicated in the top left. The dashed diagonal line represents the linear regression fit. (D) Log-likelihood shifts upon introduction of premature termination codons (PTCs). Boxplots illustrating the change in per-site log-likelihood after simulated PTC substitutions in the coding sequences (CDS) of representative prokaryotic (left) and eukaryotic (right) species. The red dashed line at y = 0.000 represents the wild-type baseline. Box boundaries denote the interquartile range (IQR), center lines represent the median, and whiskers extend to 1.5 IQR, with outliers plotted as individual points.

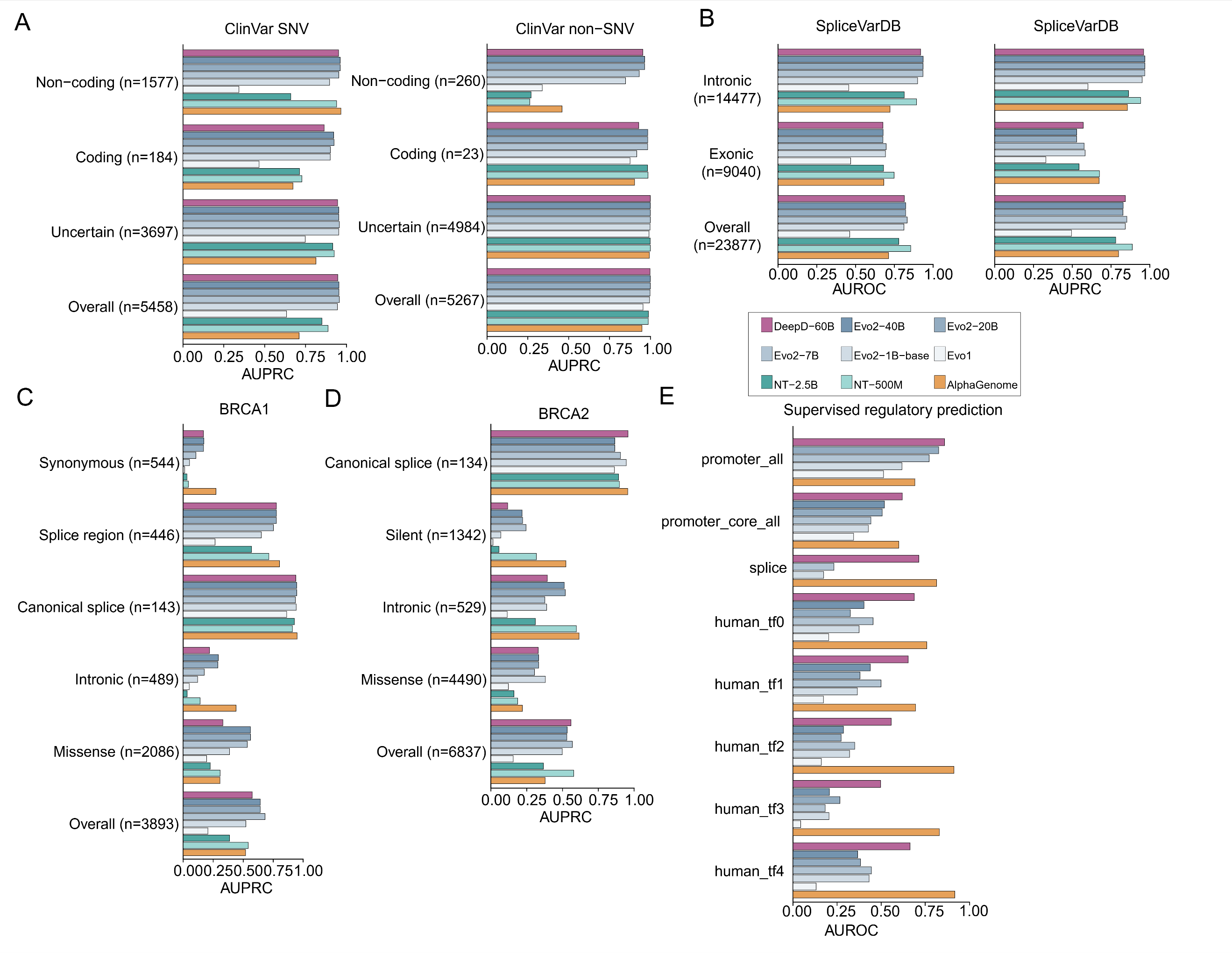

**Figure S3. Human variant-effect and supervised regulatory prediction benchmarks.** (A) Area under the precision–recall curve (AUPRC) for predicting the effects of ClinVar single-nucleotide variants (SNVs) and non-SNVs, stratified by coding annotation. (B) Area under the receiver operating characteristic curve (AUROC; left) and AUPRC (right) for distinguishing splice-altering from control variants in SpliceVarDB, shown overall and for exonic and intronic variants. (C and D) AUPRC for *BRCA1* (C) and BRCA2 (D) variant-effect prediction, shown overall and by variant consequence. (E) AUROC for supervised prediction of human promoters, core promoters, splice sites, and five transcription factor-binding datasets using model representations. Models include DeepD-60B, the Evo series, NT-2.5B, NT-500M, and AlphaGenome. Numbers in parentheses indicate the number of variants in each category.

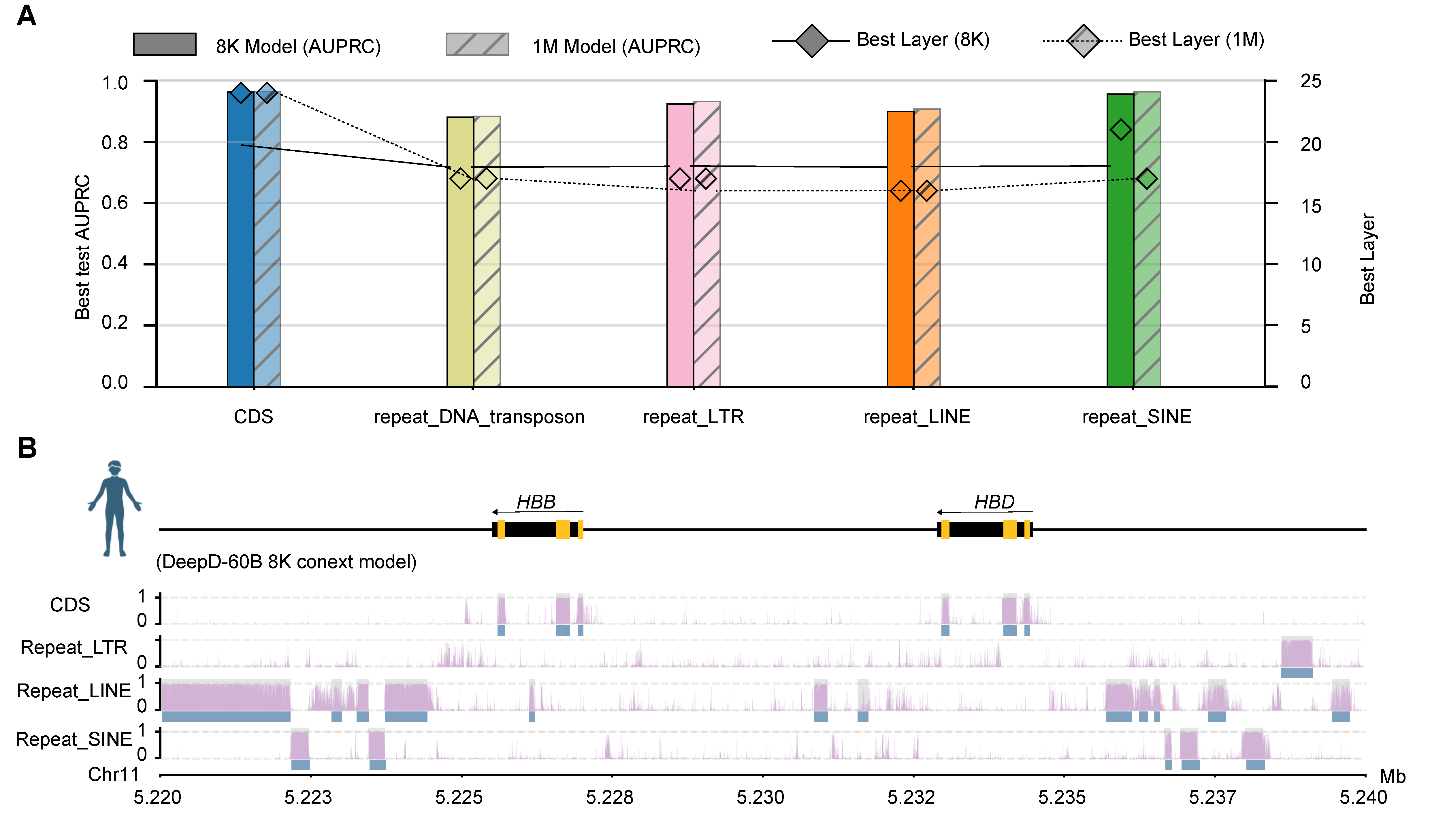

**Figure S4. Layer-wise decoding of human genomic features across context lengths.** (A) Bar plot (left y-axis): test-set AUPRC for the best-performing classifier per annotation category in the human genome. All categories achieve AUPRC > 0.9. Dashed line with diamond markers (right y-axis): optimal extraction layer for each category. Coding features (CDS, exon) are best decoded from the deepest layer (layer 24), while repetitive element classes are optimally represented at middle-to-deep layers (16–17). (B) High-resolution decoding of coding and repetitive features in the human genome with routing weights with DeepD-60B (8K). Predicted probability tracks are displayed in purple and ground-truth annotations are indicated as blue boxes underneath each track.

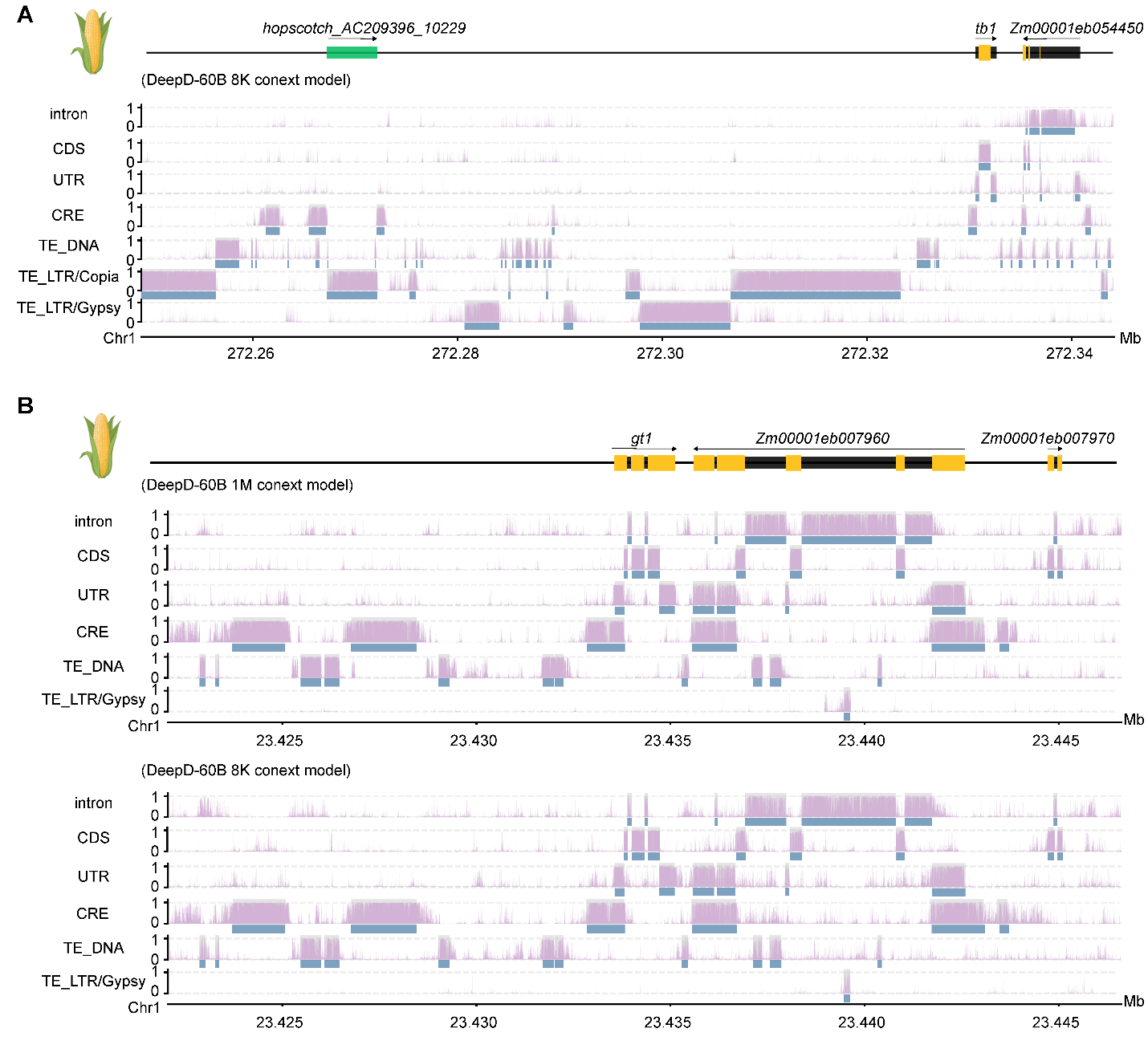
**Figure S5. High-resolution decoding at the maize *tb1* and *gt1* loci.** (A) Decoding of maize genomic features around *tb1* region with routing weights with DeepD-60B (8K). Predicted probability tracks are displayed in purple and ground-truth annotations are indicated as blue boxes underneath each track. The hopscotch LTR/Copia retrotransposon insertion is indicated by a light green box. (B) Decoding of maize genomic features around *gt1* region with routing weights with DeepD-60B (1M) (upper plot) and DeepD-60B (8K) (bottom plot).

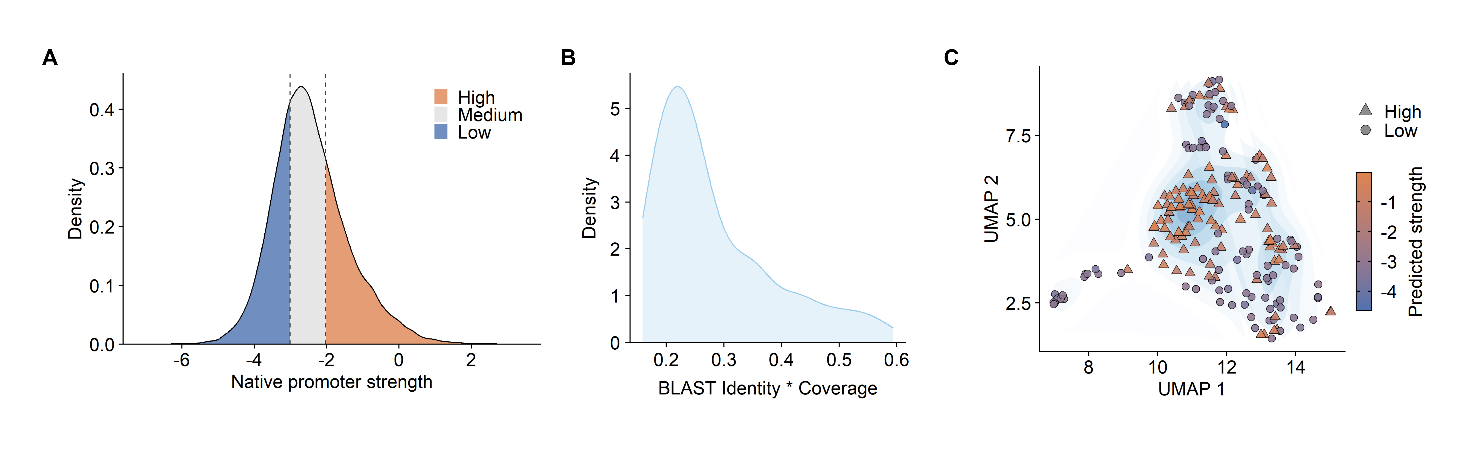

**Figure S6. Native data distributions and predictive evaluation of regulatory elements conditionally generated by DeepD.** (A) Empirical density distribution of native maize STARR-seq core promoter strengths (n = 26,572). Dashed lines indicate the strength thresholds (−3.007 and −2.018) used to categorize the training set into Low (bottom 30%), Med (middle 40%), and High (top 30%) conditioning labels. (B) Density distribution of the Identity × Coverage metric derived from BLAST searches against the NCBI nt database for the selected maize candidate sequences (n = 200). (C) UMAP projection of the selected High (n = 100) and Low (n = 100) candidate sequences based on DeepD embeddings. Point shapes represent the conditioning labels used during generation, and the color gradient reflects the predicted promoter strength.

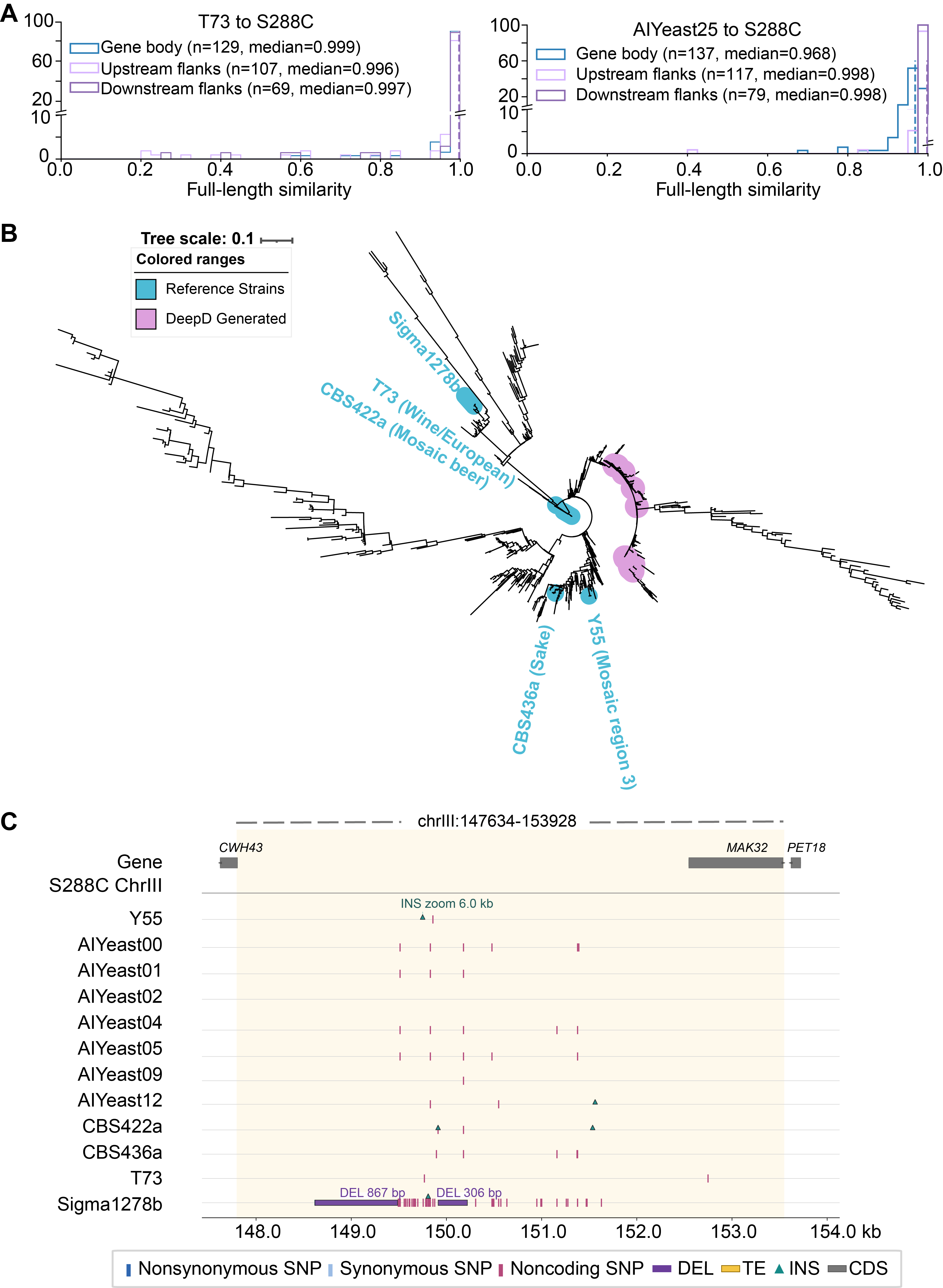

**Figure S7. DeepD-generated ChrIII sequences recapitulate yeast population diversity.** (A) Distributions of full-length sequence similarity to S288C for protein-coding gene bodies and their upstream and downstream flanking regions. The left panel compares the natural isolate T73 with S288C, whereas the right panel compares DeepD25 with S288C. (B) Phylogenetic tree inferred from synonymous variants in natural *S. cerevisiae* chromosome III sequences and DeepD-generated chromosomes. The outer track denotes sample identity and clade assignment: colors indicate natural yeast clades, and black indicates generated chromosomes. (C) Comparative view of sequence variation within the approximately 5-kb non-coding region around 150 kb on ChrIII. DeepD-generated chromosomes and representative natural isolates were aligned to the S288C reference interval (ChrIII:147,634–153,928). Colored marks denote synonymous, nonsynonymous, and non-coding variants; triangles indicate structural-variant insertions and deletions; gray and yellow boxes indicate genes and transposable elements, respectively; and connecting blocks show collinear one-to-one alignments.

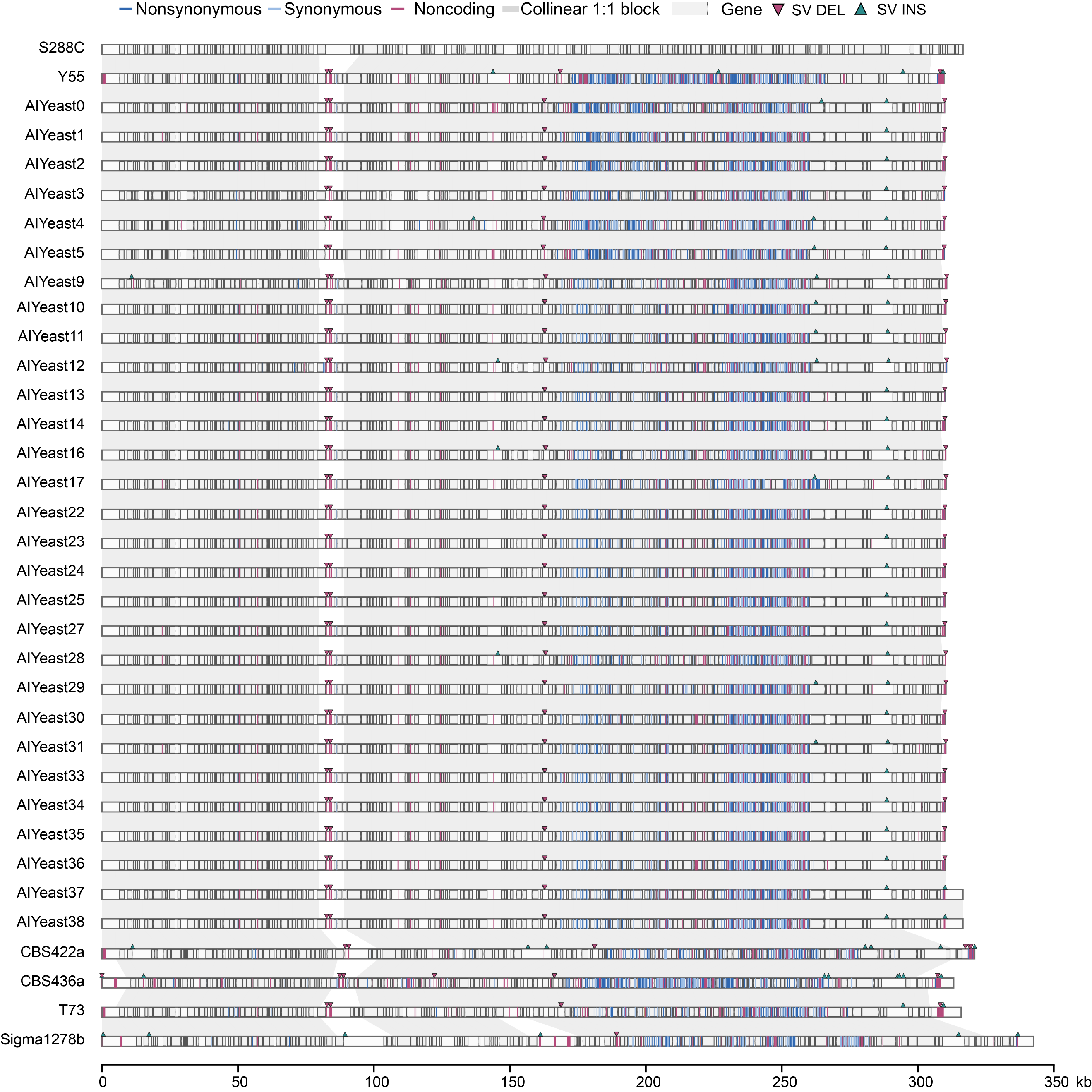

**Figure S8. DeepD-generated ChrIII sequences recapitulate yeast population diversity.**

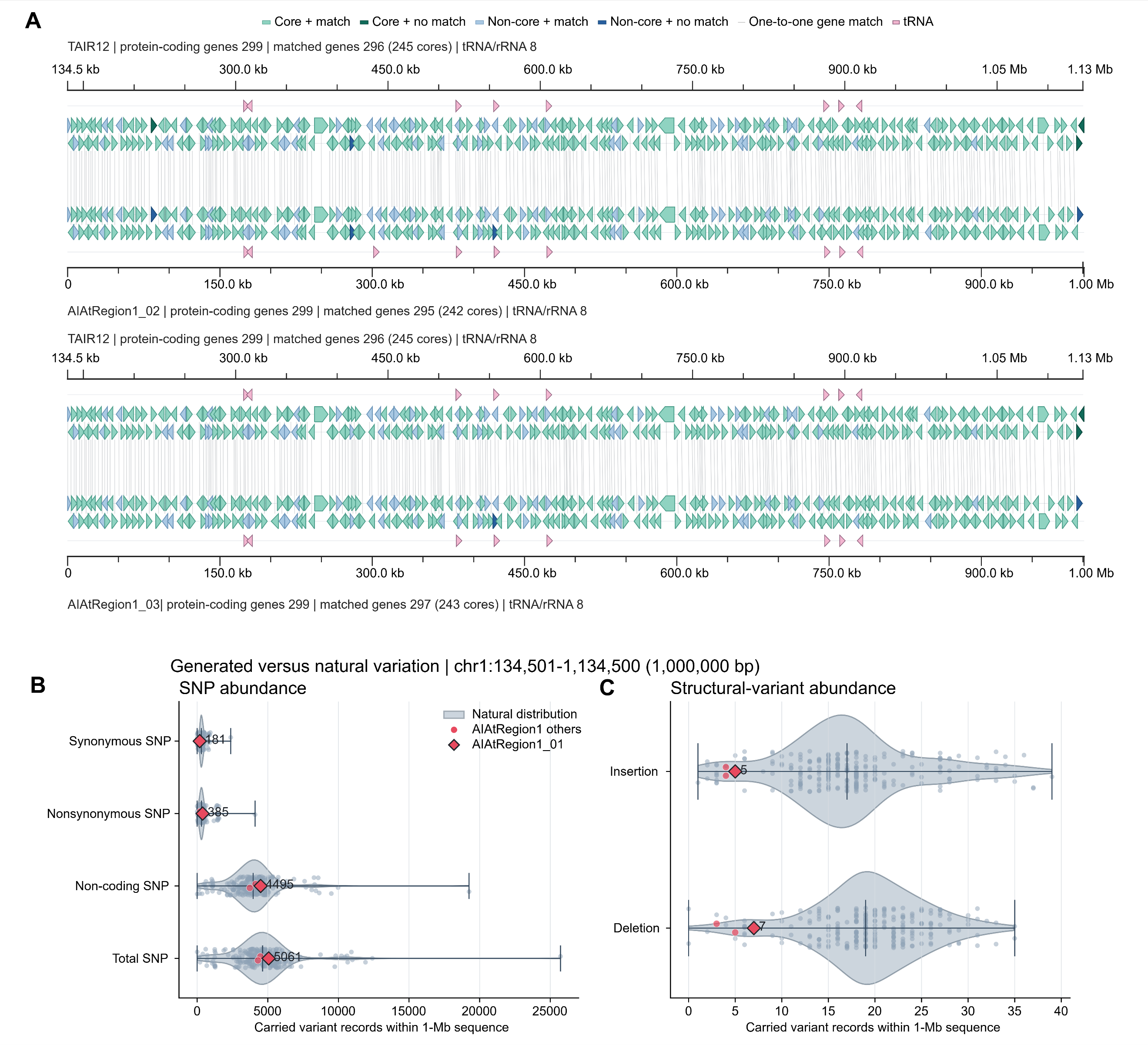

**Figure S9. Gene collinearity and natural-like variation in independently generated *Arabidopsis* sequences.**
(A) Gene-level synteny between TAIR12 and two additional independently generated 1-Mb sequences, AIAtRegion1_02 and AIAtRegion1_03. Core and non-core gene matches, unmatched genes, one-to-one gene correspondences, and tRNA genes are indicated.
(B) Numbers of synonymous, nonsynonymous, non-coding, and total SNPs within the generated 1-Mb region compared with their distributions across 255 natural *A. thaliana* accessions.
(C) Numbers of insertions and deletions in generated sequences compared with the corresponding distributions across natural accessions.
