## Supplementary material for "DeepD enables efficient genome modeling and controllable long-context sequence generation across domains of life": Method Details

### METHODS DETAILS

### 1 Training DeepD

DeepD is trained with next-token prediction on a byte-tokenized, internally curated, annotation-enriched DNA corpus. We adopt a two-stage training strategy: (1) pretraining at an 8,192-token context to learn local functional patterns and sequence grammar, and (2) midtraining with progressive context extension up to 1M tokens, with increased inclusion of whole-genome sequences. We also train and release a smaller DeepD-2B base model at 8,192-token context for 1T tokens. For computational efﬁciency, training uses sequence packing together with distributed sequence parallelism.

#### 1.1 Model architecture

DeepD is a sparse Mixture-of-Experts (MoE) model family with high representational capacity and efﬁcient decoding through sparse activation. For MoE routing, we use a species-aware expert selection strategy in which the gating network takes the concatenation of token representation and species representation as input, allowing experts to capture both sequence-level and species-speciﬁc signals. In the attention pathway, we insert a minimal sigmoid gate immediately after the self-attention output, introducing input-dependent nonlinearity and adaptive sparsity.^1^ This modiﬁcation improves model quality and training stability at scale, reduces attention-sink behavior, and strengthens long-context generalization. To balance local and global dependency modeling, and to preserve the ability to learn both short-range and long-range DNA signals, 75% of attention heads use sliding-window attention (SWA) with window size 256, while the remaining 25% use full attention (FA). Positional encoding uses RoPE with YaRN-based scaling for stable extrapolation to ultra-long contexts.^2^ The same architectural principles are used in both DeepD-60B and DeepD-2B variants. Model-scale MoE conﬁgurations are summarized in Supplementary Table 1.

|  | DeepD-2B | | DeepD-60B |
| --- | --- | --- | --- |
| Total parameters | | 1.8B | 63B |
| Active parameters | | 170M | 1.3B |
| Hidden size | | 768 | 2048 |
| FFN hidden size | | 256 | 768 |
| Total layers | | 24 | 24 |
| Num heads | | 16 | 32 |
| Total experts | | 128 | 576 |
| Activated experts | | 8 | 8 |

**Supplementary Table 1** Model architecture conﬁgurations for DeepD models

#### 1.2 Loss function

DeepD is optimized with a reweighted cross-entropy objective to reduce over-dominance of low-information genomic regions (e.g., repetitive) and improve calibration on functionally informative loci:

$l_{\mathrm{wCE}}=\frac{1}{Z}\sum_{t} \omega_{t}l_{\mathrm{CE}}\left( t \right)$,

Where

$$\omega_{t}=\left\{ \begin{aligned} 0.1, &if position t is repetitive \\ 1, &ifposition t is nonrepetitive \end{aligned} \right.$$

and $Z=0.1N_{\mathrm{repeat}}+N_{\mathrm{nonrepeat}}$. During early pretraining, auxiliary annotation indicators are provided to distinguish repetitive and unannotated sequence from other regions, and loss is masked on species-tag tokens.

#### 1.3 Pretraining infrastructure

The training stack is based on DeepSeekMoE and is optimized for large-scale long-context MoE language modeling.^3^ At the model level, DeepD uses 576 experts (including one shared expert) and activates 8 experts per token. To improve routing stability in shallow layers, the first FFN block is kept dense. We further stabilize expert utilization with a combination of auxiliary-loss-free load balancing and sequence-wise balancing losses.

At the systems level, we use mixed-precision training with DeepSpeed Ulysses sequence parallelism and activation checkpointing to control memory under long contexts.^4^ These components are jointly configured to support efficient scaling from 8k pretraining to 1M-context midtraining.

#### 1.4 Pretraining phase

Pretraining establishes the base DeepD models before long-context extension. We train both DeepD-2B and DeepD-60B with a fixed context length of 8,192 tokens using next-token prediction on the curated genomic corpus. In this phase, DeepD-2B is trained for 1T tokens, while DeepD-60B is trained for 5.69 T tokens. The 8,192-token regime is chosen to prioritize robust learning of local sequence grammar, coding regularities, and short-to-mid range functional dependencies under stable optimization and high hardware throughput. Parameters are shown in Supplementary Table 2. The resulting checkpoints are then used as initialization for the staged midtraining procedure that extends context length to 1M tokens.

|  | DeepD-2B base | DeepD-60B base |
| --- | --- | --- |
| Learning Rate | 5.00e-04 | 3.00e-04 |
| Training Batch | 9.44M | 36.70M |
| Total Iterations | 105,963 | 151,500 |
| Pretraining Tokens | 1.00T | 5.69T |
| Sequence length | 8,192 | 8,192 |

**Supplementary Table 2** Training hyperparameters for DeepD models. Each model uses the Adam optimizer with $\beta_{1}=0.9$, $\beta_{2}=0.95$, and cosine learning rate decay.^5^

#### 1.5 Midtraining phase: Context extension

During midtraining, we extend the context length in stages from 8k to 32k, 128k, 512k, and finally 1M tokens. At each stage, the model is warm-started from the previous checkpoint to preserve short-context capabilities while adapting to longer-range dependencies. We adopt YaRN-based RoPE scaling during this transition so that positional encoding remains stable under long-context extrapolation.^2^ The stage-wise configuration used in practice is shown in Supplementary Table 3.

| Context Length | Training Tokens (60B) | Use YaRN | Max position embeddings | Original max position |
| --- | --- | --- | --- | --- |
| 32k | 100B | Yes | 32,768 | 8,192 |
| 128k | 50B | Yes | 131,072 | 8,192 |
| 512k | 50B | Yes | 524,288 | 8,192 |
| 1M | 50B | Yes | 1,048,576 | 8,192 |
| Total | 250B | - | - | - |

**Supplementary Table 3** Midtraining context extension protocol for DeepD-60B with YaRN-based settings.

#### 1.6 Inference infrastructure

For inference and large-scale evaluation, we deploy DeepD on a vLLM-based serving stack to maximize throughput and memory efficiency.^6^ This setup leverages vLLM’s paged key-value (KV) cache management and continuous batching to support long-context autoregressive decoding under high-concurrency workloads.^6^ We use the same model checkpoints as in training and configure inference with long-context-compatible runtime settings to ensure stable generation quality at extended sequence lengths.

### 2 DeepD training data

DeepD was trained on a total of 5.39 Tb nucleotides from eukaryotic genomes, 452.60 Gb from bacterial genomes, 12.09 Gb from archaeal genomes, and 45.22 Gb nucleotides from viral genomes. The token distribution was designed to reflect the composition of real genomic sequences, rather than being restricted to gene-coding regions alone. In addition to nucleotide sequences, gene annotation tracks were available for 1,215 of 5,434 eukaryotic species and for all prokaryotic species. Furthermore, we downloaded phylogenetic tree information for all species from [Open Tree of Life](https://tree.opentreeoflife.org/about/open-tree-of-life).^7^ Together, these three data modalities-genome sequence, gene annotation, and phylogenetic information-constituted the core inputs used during DeepD training.

#### 2.1 Genome collection and filtering

**Eukaryotic genomes.** Eukaryotic reference genomes were downloaded from NCBI [GenBank assembly summary](https://ftp.ncbi.nlm.nih.gov/genomes/ASSEMBLY_REPORTS/assembly_summary_genbank.txt) on September 4, 2025. To exclude organellar interference, non-nuclear sequences, including mitochondrial and chloroplast genomes, were removed. Initial collection retained 19,880 reference genomes, totaling 14.98 Tb and spanning 2,828 families. To reduce redundancy while preserving phylogenetic diversity, genomes were first deduplicated at the genus level by retaining only the single representative species with the largest contig N50 within each genus, yielding 9,092 genomes. Assemblies with contig N50 < 1 Mb were then removed, leaving 4,538 species. To restore taxonomic representation lost during this filtering step, the species with the highest contig N50 was reintroduced for each missing family, expanding the dataset to 5,716 species and recovering the full set of 2,828 families. In addition, 8 manually selected model and agriculturally important species (*O. sativa, T. aestivum, D. melanogaster, G. max, C. elegans, S. bicolor, S. cerevisiae, O. aries*) were added, resulting in a total of 5,730 species.

To improve sequence continuity, genomes with a high proportion of ambiguous nucleotides (“N”) were then curated at the family level by replacing them with higher-quality species from the same family. Species with a genome-wide N content > 1% were removed. After sequence cleaning, 5,440 eukaryotic genomes were retained. To preferentially use higher-quality annotation standards, the retained species were cross-matched against NCBI [RefSeq assembly summary](https://ftp.ncbi.nlm.nih.gov/genomes/ASSEMBLY_REPORTS/assembly_summary_refseq.txt), and 588 GenBank assemblies were replaced by their corresponding RefSeq versions.

All retained eukaryotic genomes were processed using a sequence-cleaning pipeline. All retained prokaryotic and eukaryotic genomes were processed using a unified sequence-cleaning pipeline. First, ambiguous nucleotides (“N”) at both ends of each contig were trimmed. Contigs were then filtered by length, with minimum retained lengths of 10 kb for eukaryotic genomes and 1 kb for prokaryotic genomes. To ensure data integrity, any contig containing invalid characters was removed in its entirety. To identify and handle local low-quality regions, each sequence was scanned using a 50-kb sliding window. If the proportion of non-standard bases within a window exceeded 10%, the sequence was split at that region, and a special delimiter token (“@”) was inserted between the resulting fragments to indicate that they originated from the same contig and remained adjacent in genomic space. Finally, a stringent global quality filter was applied by removing any contig with a total non-standard base content > 1% across the full sequence. After sequence processing, the eukaryotic dataset comprised 5,434 genomes, totaling 5.39 Tb and representing 2,697 families.

**Prokaryotic genomes.** Prokaryotic genome assemblies were obtained from the [Genome Taxonomy Database](https://gtdb.ecogenomic.org/) (GTDB, release 226), which provides a phylogenetically standardized taxonomy for Bacteria and Archaea.^8^ We selected all species-representative genomes defined by GTDB, yielding 143,614 candidate assemblies, including 136,646 bacterial and 6,968 archaeal genomes. Assembly quality was then filtered using CheckM (v1 and v2), a machine learning–based framework for estimating microbial genome completeness and contamination.^9^ Genomes with completeness ≤ 50.0% or contamination ≥ 10.0% were removed. This excluded 81 low-quality assemblies, resulting in a final set of 143,533 high-quality prokaryotic representative genomes with a total size of 465.03 GB. Among these genomes, 136,578 were bacterial genomes, accounting for 452.90 Gb, and 6,955 were archaeal genomes, accounting for 12.13 Gb. To mask repetitive elements for downstream analyses, all retained genomes were subjected to soft-masking using RepeatMasker v4.2.2 with parameters -species bacteria and -xsmall.^10^

All retained prokaryotic genomes were processed using the same sequence-cleaning pipeline used for eukaryotic genomes. The final prokaryotic dataset comprised 136,485 bacterial genome assemblies totaling 452.60 Gb and 6,932 archaeal genome assemblies totaling 12.09 Gb.

**Virus genomes.** Viral genomes were obtained from the [IMG/VR database](https://img.jgi.doe.gov/vr/) (v7) and restricted to entries annotated as high_confidence. To minimize redundancy, a single representative genome was selected for each viral operational taxonomic unit (vOTU), prioritizing the sequence with the highest reported quality and, in cases of tied quality, the longest genome. This procedure yielded a final dataset of 2,917,521 high-quality viral genomes, totaling 45.22 GB.

#### 2.2 Sequence processing and tokenization

Genomic positions were classified as repetitive or non-repetitive based on sequence masking and genome annotations. For genomes with gene annotation tracks, uppercase bases in the corresponding soft-masked FASTA sequences, together with bases annotated as coding sequences (CDSs) or introns in the GFF files, were classified as non-repetitive. Annotated CDS and intron positions were retained as non-repetitive even when they overlapped lowercase bases. All remaining positions were classified as repetitive. For genomes lacking gene annotation tracks, classification relied solely on the soft-masked FASTA sequences, with uppercase bases classified as non-repetitive and lowercase bases classified as repetitive. Viral genomes were classified in their entirety as non-repetitive.

For pretraining, 8,192 genomic windows are segmented and packed for efficient training, with optional reverse-complement augmentation to improve strand robustness. For long-context midtraining stages, contigs from the same source genome are stitched under controlled rules so that the model observes biologically meaningful long-range context windows. For long-context sequence training, we apply a hierarchy-aware balanced sampling strategy over the phylogenetic tree. We first set target sampling budgets at the kingdom level (e.g., Animalia, Plantae, Fungi, Bacteria, Archaea, and viruses), then recursively allocate sampling mass to lower taxonomic levels (phylum/class/order/family/genus/species). Within each level, species sampling is temperature-scaled by sequence count to up-weight underrepresented taxa. Let $N_{i}$ denote the number of sequences for species $i$; its sampling probability is

$$p_{i}=\frac{N_{i}^{\alpha}}{\sum_{j} N_{i}^{\alpha}}$$

where $\alpha$ controls the balancing strength ($\alpha<1$ increases the relative probability of low-resource species). In all experiments, we set $\alpha=0.3$.

#### 2.3 Phylogenetic tree embedding

To encode hierarchical evolutionary relationships, we compute phylogenetic tree embeddings using the HS-DTE method described in van Spengler and Mettes and its official implementation.^11^ For each tree, we use constructive initialization with optimization-based generation at 32 dimensions (temperature 0.02, one terminal component, float64 precision). We then map the resulting 32-dimensional embeddings to 2,048 dimensions via a fixed projection-plus-expmap conversion pipeline and use the converted vectors as species representations during model training and inference.

### 3 Prediction evaluations

#### 3.1 General model scoring and evaluation framework

To enable consistent evaluation across prediction benchmarks, we defined model-specific scoring procedures according to model architecture and applied task-specific evaluation metrics according to the type of experimental label. For binary classification tasks, model-derived variant-effect scores were used as ranking scores, and performance was quantified using the area under the receiver operating characteristic curve (AUROC) and the area under the precision-recall curve (AUPRC). For deep mutational scanning (DMS) tasks with continuous experimental measurements, performance was quantified using the absolute Spearman rank correlation coefficient between model-derived scores and the reported experimental phenotypes.

DeepD and the Evo-series models^12,13^ were evaluated as autoregressive nucleotide language models using sequence log-likelihood. The sequence log-likelihood was calculated by summing the conditional log-probability of each nucleotide given its preceding sequence context.

$$LL(S)=\sum_{i=1}^{L} logP(s_{i} | s_{<i})$$

For binary variant-effect classification tasks, the model-derived variant-effect score was defined as the difference in sequence log-likelihood between the wild-type and mutant sequences, calculated as wild-type minus mutant.

$$s_{variant}=LL(S_{wt})-LL(S_{mut})$$

Larger values therefore indicate a greater reduction in sequence likelihood caused by the mutation. The resulting score was used directly to calculate AUROC and AUPRC. For DMS tasks with continuous experimental measurements, all variants within each assay were derived from the same wild-type reference sequence. Because subtraction of the same wild-type log-likelihood does not affect the rank ordering of variants within an assay, the sequence log-likelihood of each mutant sequence was used directly as the model-derived score, and prediction performance was evaluated using the absolute Spearman rank correlation coefficient against the experimental phenotype.

AlphaGenome^14^ was evaluated using its official API. For each input sequence, equal numbers of ambiguous nucleotides (N) were symmetrically padded to the 5’ and 3’ ends to reach a fixed input length of 16,384 bp. For each variant, the corresponding wild-type and mutant sequences were provided to the AlphaGenome API to obtain predicted variant-effect scores. The composite_score was used for ClinVar, *BRCA1*, *BRCA2*, and the corresponding human protein DMS benchmarks, whereas the delta_score was used for SpliceVarDB. These scores were used directly for downstream evaluation. For binary classification benchmarks, the corresponding AlphaGenome score was used as the ranking score and performance was quantified using AUROC and AUPRC. For human protein DMS benchmarks with continuous experimental measurements, performance was evaluated using the absolute Spearman rank correlation coefficient between the composite_score and the reported phenotype. For the GUE regulatory benchmarks, AlphaGenome was instead evaluated using task-specific native functional outputs, as described in the corresponding subsection.

Two Nucleotide Transformer (NT) version 2 models^15^ were evaluated as embedding-based zero-shot baselines: NT-2.5B multi-species and NT-500M multi-species. Wild-type and mutant nucleotide sequences were independently converted to upper case, tokenized using the NT 6-mer tokenizer, and passed through the corresponding encoder. A sequence-level representation was obtained by mean-pooling the final-layer hidden states over all non-padding tokens, including the classification token. The variant-effect score was then calculated as the cosine distance between the pooled representations of the wild-type and mutant sequences.

$$s_{NT}=1-\frac{hwt \cdot hmt}{\parallel hwt\parallel\parallel hmt\parallel}$$

Larger cosine-distance values indicate a greater displacement between the wild-type and mutant sequence representations and were interpreted as stronger predicted variant effects. When the tokenized sequence exceeded the model context length, tokens were retained from the 5’ end of the input. After accounting for the classification token, NT-2.5B retained at most 999 6-mer tokens, corresponding to 5,994 bp, whereas NT-500M retained at most 2,047 6-mer tokens, corresponding to 12,282 bp. For binary classification benchmarks, the cosine-distance score was used directly as the ranking score for AUROC and AUPRC calculation. For protein DMS benchmarks with continuous experimental measurements, performance was quantified using the absolute Spearman rank correlation coefficient between the cosine-distance score and the reported phenotype.

#### 3.2 Zero-shot protein fitness prediction

To evaluate the ability of DeepD to perform zero-shot fitness prediction on protein variant datasets, we assembled a benchmark comprising 7 prokaryotic tasks and 10 eukaryotic tasks.

For the prokaryotic benchmark, six tasks were obtained from ProteinGym, each derived from a previously published deep mutational scanning (DMS) study.^16^ These tasks were designated as Firnberg,^17^ Jacquier,^18^ Kelsic,^19^ Ecoli_Weeks,^20^ Rockah,^21^ and Chen.^22^ Weeks and Ecoli_Weeks were derived from the same study but used different nucleotide sequence inputs. Weeks was scored on the native nucleotide sequence reported in the original paper, whereas Ecoli_Weeks was reconstructed by reverse-translating the ProteinGym protein sequence using species-specific high-frequency codons. These two tasks differed in the input sequences used for model evaluation. Because DeepD operates on nucleotide sequences, we reverse-translated the ProteinGym protein sequences into coding DNA sequences using the preferred codons of the corresponding species, and used the resulting nucleotide sequences for scoring.

For the eukaryotic benchmark, we evaluated 10 tasks spanning yeast, mouse, and human datasets. Among these, 9 DMS tasks were obtained from ProteinGym and were originally represented as protein sequences. To make these datasets compatible with DeepD, we applied the same reverse-translation strategy as described above, converting each protein sequence into a nucleotide sequence using the high-frequency codon usage of the corresponding species. These 10 tasks were Yeast,^23^ Human_Faure,^24^ Sun,^25^ MOUSE_1JFM, HUMAN_1V1C, HUMAN_2LO1, HUMAN_3F0M, HUMAN_2O2W, MOUSE_1K1V, and MOUSE_1E0L.^26^ The task Sun was evaluated directly using the original nucleotide sequences reported in the source publication.^25^

For each task, performance was evaluated using the absolute Spearman correlation coefficient between the experimental fitness values and the model scores, following the model-specific scoring procedures described in Section 3.1. For autoregressive models and masked language models, sequence likelihood and pseudo-likelihood were used as scoring criteria, respectively. We performed a cross-model comparison of DeepD against nucleotide models such as Evo1, Evo2 40B, Evo2 7B, Evo2 1B,^12,13^ as well as protein language models, including ESM series,^27,28^ ProGen2 series,^29,30^ and CARP-640M.^31^ AlphaGenome was additionally evaluated on the corresponding human protein DMS tasks for which compatible predictions were available.

#### 3.3 Zero-shot ncRNA fitness prediction

We adopted the same set of seven ncRNA DMS tasks curated in the Evo2 study, including: Zhang (an rRNA mutagenesis study),^32^ Pitt (a ribozyme DMS study),^33^ Hayden (a ribozyme DMS study),^34^ Guy (a tRNA DMS study),^35^ Kobori (a ribozyme DMS study),^36^ Domingo (a tRNA DMS study),^37^ and Andreasson (a ribozyme DMS study).^38^ These datasets provide experimentally measured fitness values for single mutations and combinatorial mutations, enabling a systematic assessment of mutation effects across ncRNA variants.

To quantify prediction accuracy, we calculated the absolute Spearman correlation coefficient between experimental fitness and model-derived scores. DeepD and the Evo-series models were evaluated following the model-specific scoring procedures described in Section 3.1, whereas RiNALMo,^39^ a masked language model, was scored using sequence pseudo-likelihood. We compared DeepD with several nucleotide-based models, including Evo1, Evo2 40B, Evo2 7B, Evo2 1B, and RiNALMo, to assess performance in predicting ncRNA variant effects.

#### 3.4 Genome understanding evaluation benchmark

We used the genome understanding evaluation (GUE) benchmark introduced in the DNABERT-2 study, which provides a standardized multi-species evaluation suite for genomic sequence classification tasks.^40^ All datasets and train/validation/test splits used here were obtained following the DNABERT-2 benchmark release at [Github folder](https://github.com/MAGICS-LAB/DNABERT_2/tree/main#2-model-and-data). From this resource, we selected 15 downstream tasks, including transcription factor prediction for 5 human TFs and 5 mouse TFs, human promoter detection, human core promoter detection, human splice site prediction, and yeast epigenetic marks (H3 and H4) prediction.

For all GUE tasks evaluated with pretrained genome language models, we adopted a unified frozen-backbone probing protocol. Each DNA sequence was first passed through the pretrained genome language model to extract and store token-level, layer-normalized hidden states, while keeping the backbone model fully frozen throughout downstream training. Task-specific prediction was then performed using a lightweight two-layer multilayer perceptron (MLP) classifier with architecture input_dim🡪512🡪128🡪 K, where K denotes the number of output classes. The value of input_dim depended on the pretrained model; for DeepD, input_dim = 2048. Models were trained using cross-entropy loss, with Matthews correlation coefficient (MCC) on the validation set used for model selection and early stopping. Final performance was reported on the independent test set using AUROC.

AlphaGenome was additionally evaluated in a zero-shot setting on five human TF-binding tasks, human splice site prediction, promoter detection, and core promoter detection using its native functional outputs. No GUE training or validation data were used for model fitting. Test sequences were centered within the minimum supported AlphaGenome input length of 16,384 bp and padded with ambiguous nucleotides (N). For TF-binding tasks, CHIP_TF predictions corresponding to the target TF and, when available, the matched biosample were used; when the exact biosample was unavailable, all available tracks for the target TF were retained. The maximum predicted signal over bins overlapping the original GUE sequence and the selected tracks was used as the TF-binding score. For splice site prediction, SPLICE_SITES donor and acceptor probabilities were extracted within ±2 bp of the central splice boundary, and performance across acceptor, donor, and non-splice classes was summarized by macro-averaged one-vs-rest AUROC and AUPRC. For promoter and core promoter prediction, PRO-cap signals within ±5 bp of the expected transcription start site were extracted across all available human tracks, and the maximum signal was used as the promoter score. Binary TF-binding and promoter tasks were evaluated using AUROC and AUPRC.

Potential overlap between the GUE regulatory benchmarks and the AlphaGenome training data was assessed by cross-referencing the provenance of the five human TF-binding, promoter, and splice-site benchmarks used in DNABERT with the AlphaGenome supplementary track metadata. We found no evidence that these exact benchmark datasets were directly included in AlphaGenome training. However, AlphaGenome was trained on more recent datasets measuring the same regulatory outputs and, in some cases, on data from related or overlapping biosamples. Overlap at the level of genomic intervals, biological samples, or regulatory signals could therefore not be fully excluded. Consequently, AlphaGenome’s performance on the GUE regulatory benchmarks may represent an optimistic estimate and should not be interpreted as a strictly out-of-distribution comparison with self-supervised genome language models.

#### 3.5 Phage gene essentiality prediction

We performed zero-shot essentiality prediction for genes from two phage genomes, lambda (NC_001416) and P1 (NC_005856), with the same dataset curated in the Evo2 study, containing 162 genes in total. The genomic sequences used for model input were segments of 8,192 bp, including the gene body and its symmetric flanking sequences. For genes longer than this limit, we truncated the sequence to the first 8,192 bp for computation.

Prediction performance was assessed by calculating the wild-type and mutant log-likelihood difference following the scoring procedure described in Section 3.1. The mutation strategy involved inserting a stop codon 12 nucleotides downstream in the coding sequence. AUROC and AUPRC were used to evaluate gene-essentiality classification performance. We compared the performance of DeepD against Evo1 and Evo2 series to evaluate the relative prediction accuracy.

#### 3.6 ClinVar variant effect prediction

We used the ClinVar release dated December 15, 2024 to assess model performance in recognizing human disease-associated variants.^41^ To construct a high-confidence benchmark, we retained only variants with review status of at least three stars, corresponding to entries in the CLNREVSTAT field annotated as “reviewed by expert panel” or “practice guideline”. For class labels, variants were assigned to the benign category if the CLNSIG field was annotated as “Likely benign,” “Benign,” or “Benign/Likely benign,” and to the pathogenic category if annotated as “Pathogenic,” “Pathogenic/Likely pathogenic,” or “Likely pathogenic.” For each variant, we extracted the variant-centered sequence with 4 kb of flanking sequence on each side.

We formulated ClinVar variant-effect prediction as a binary classification task distinguishing pathogenic variants from benign variants. Variant-effect scores were calculated using the model-specific scoring procedures described in Section 3.1, and performance was evaluated using AUROC and AUPRC. SNVs and non-SNVs were analyzed as separate benchmark groups. The SNV benchmark included 5,458 variants, among which 184 were annotated in coding regions, 1,577 in non-coding regions and 3,697 uncertain regions. The non-SNV benchmark included 5,267 variants, including 23 coding-region variants, 260 non-coding-region variants and 4,984 uncertain variants. Because many variants lacked confident coding/non-coding regional assignment, we first evaluated all variants within each benchmark as an overall task, and then separately evaluated the coding and non-coding subsets.

#### 3.7 SpliceVarDB variant effect prediction

Zero-shot splice-disrupting variant prediction was evaluated using SpliceVarDB,^42^ an experimentally validated database of human splicing variants. We used the same curated benchmark dataset as reported in the Evo2 study to enable direct comparison. Variants were classified as splice-disrupting or neutral using the model-specific variant-effect scores described in Section 3.1, and performance was quantified using AUROC and AUPRC. The benchmark included 23,877 variants, of which 9,040 were annotated as exonic and 14,477 as intronic. Because 360 variants could not be confidently assigned to either category, we evaluated all variants together as the overall benchmark and separately assessed the exonic and intronic subsets.

#### 3.8 *BRCA1/2* zero-shot classification

For the *BRCA1* benchmark,^43^ we used a total of 3,893 variants. Functional-region annotations were extracted directly from the published descriptions. Each variant was assigned to one unique functional region. The seven functional-region categories were 5’ UTR (n = 47), Canonical splice (n = 143), Intronic (n = 489), Missense (n = 2,086), Nonsense (n = 138), Splice region (n = 446), and Synonymous (n = 544). The 5′UTR and Nonsense subsets were not analyzed separately because of their relatively small sample sizes, substantial class imbalance, and unstable classification performance, as reflected by large discrepancies between AUROC and AUPRC. Variant-effect labels were processed following the Evo2 protocol. Specifically, variants labeled as loss-of-function (LOF) or functional/intermediate were used to define the binary variant-effect classification. Thus, variant-effect prediction within each functional-region category was formulated as a binary classification task. Model-specific variant-effect scores were calculated following the procedures described in Section 3.1, and AUROC and AUPRC were computed separately within each analyzed functional-region category.

For the *BRCA2* benchmark,^44^ we used a total of 6,837 variants. Five functional-region categories were Canonical splice (n=134), Intronic (n=529), Missense (n=4,490), Nonsense (n=342), and Silent (n=1,342). The Nonsense subset was not evaluated separately because all variants in this category belonged to the positive class, precluding binary classification performance evaluation. Model-specific variant-effect scores were calculated following the procedures described in Section 3.1, and AUROC and AUPRC were computed for the overall benchmark and the remaining functional-region subsets.

### 4 Analysis of phylogeny-informed genome modeling

#### 4.1 Interpretability analysis of expert activations and GC prediction

We randomly sampled 500 gene fragments, each spanning 8,192 bp and centered on the gene start coordinates defined in the GFF annotation file, from 10 species with a broad range of genomic GC distributions. For each layer in the Mixture-of-Experts (MoE) architecture, we applied average pooling to the activation weights of the experts invoked at G or C nucleotide positions within each sequence. This average-pooled representation characterizes the response intensity of the experts in that layer to GC composition. Subsequently, we trained a Ridge regression model using the expert weights from each layer as input features and the ground-truth GC content of the sequences as target labels. We evaluated the association between the model's internal features and the physical sequence composition by calculating the R^2^ scores and Spearman correlation coefficients across different layers. All activation data were extracted from the forward pass of the model during zero-shot inference. Finally, we selected the top 10 experts with the strongest positive and negative correlations with GC content, designating them as GC-activated and AT-activated experts, respectively.

#### 4.2 GC content generation and hierarchical tag sensitivity analysis

We utilized a conditional generation strategy to validate the capability of species tags in restoring genomic GC content. We selected 107 eukaryotic and prokaryotic species: 91 randomly-selected species and 16 manually selected model and agriculturally important species (*D. melanogaster*, *M. musculus*, *A. thaliana*, *C. elegans*, *S. scrofa*, *S. bicolor*, *G. max*, *C. hircus*, *G. gallus*, *T. aestivum*, *D. rerio*, *Z. mays*, *S. cerevisiae*, *O. aries*, *O. sativa*, *B. taurus*). For each species, we uniformly provided ATG as a prompt to generate 25 sequences, each 500 bp in length, and subsequently calculated the average GC content of the generated sequences. As a control, we implemented an additional generation task that omitted species-specific tags, which was likewise prompted with ATG. Finally, the GC content of the generated sequences was evaluated against the ground-truth genomic GC content distributions of the corresponding species.

#### 4.3 Controllable cross-species codon bias recovery analysis

We conducted experiments utilizing the conditional generation mode of DeepD. Specifically, we randomly extracted forty 50-bp sequences upstream of the gene start sites from the human (GRCh38.p14, NCBI RefSeq assembly) and yeast (*S. cerevisiae* S288C, GCF_000146045.2) genomes to serve as prompts,^45^ setting the generation length to 1,000 bp. Forty sequences were independently generated for each species. Subsequently, we used Pyrodigal v3.7.1 to perform open reading frame (ORF) prediction and protein-coding functional annotation on the generated synthetic sequences.^46^ We then quantified the occurrence frequencies of all 64 codons within the generated sequences to compute their relative synonymous codon usage (RSCU),^47^ which we compared against the native RSCU profiles of the human and yeast genomes.

#### 4.4 Glutamine codon preference analysis

Ten representative eukaryotic and prokaryotic species were selected to span a broad spectrum of genomic GC content and codon usage frequencies. Reference genome sequences for 10 species from NCBI and GTDB database were used:

- *S. cerevisiae* (GCF_000146045.2)
- *O. sativa* (GCF_034140825.1)
- *H. sapiens* (GCF_000001405.40)
- *A. thaliana* (GCF_000001735.4)
- *C. Absconditicoccus praedator* (GCF_021057185.1)
- *N. aerobiophila* (GCF_023169545.1)
- *C. Korarchaeum cryptofilum* (GCF_000019605.1)
- *C. akajimensis* (GCF_000025905.1)
- *G. benzoatilyticus* (GCF_017338855.1)
- *C. Nanohalobia haloferacivicina* (GCF_029232205.1)

For each species, CAG codon sites were randomly sampled from validated CDS regions. Using the third position (G) as the target site, the preceding 4,000 bp sequence was extracted as input context for DeepD. The model generated a conditional probability distribution over the four nucleotides at the target position. The likelihood difference was calculated as

$\Delta Likelihood=\log P\left( x_{i}=A \mid\text{context} \right)-\log P\left( x_{i}=G \mid\text{context} \right)$.

The mean $\Delta Likelihood$ and the standard error of the mean (SEM) were calculated for each species to quantify the model's preference for G to A mutations. The CAG Ratio is defined as the frequency of CAG codons relative to all CA-starting codons (CAG and CAA) in the respective genome.

#### 4.5 Genomic functional region sensitivity analysis

To evaluate the model's sensitivity to diverse genomic features, we partitioned the genome into several functional categories based on species-specific annotation, including coding sequences (CDS), intergenic regions, introns (eukaryotes only), riboswitches (prokaryotes only), and various non-coding RNAs (e.g., rRNA, tRNA, and snoRNA). Reference genome sequences for 9 species from NCBI and GTDB database were used:

- *S. cerevisiae* (GCF_000146045.2)
- *O. sativa* (GCF_034140825.1)
- *H. sapiens* (GCF_000001405.40)
- *A. thaliana* (GCF_000001735.4)
- *T. aotearoense* (GCF_009905255.1)
- *G. benzoatilyticus* (GCF_017338855.1)
- *D. geothermalis* (GCF_000196275.1)
- *S. elongatus* (GCF_003846445.2)
- *R. brasiliensis* (GCF_000165715.2)

Within each functional category, we randomly sampled genomic sites and introduced 10-bp deletion mutations. For each mutation, an 8-kb genomic context centered at the deletion site was extracted.

We employed DeepD to quantify the functional impact of these mutations by calculating the change in mean log-likelihood between the wild-type and mutated sequences. To ensure strand-invariant predictions, we calculated the log-likelihood for both the forward and reverse complement sequences and used their average as the final score. The mean response values across different functional regions were then aggregated for each species to characterize the model's ability to distinguish biological significance across the genome.

#### 4.6 Premature termination codons analysis

We performed *in silico* mutagenesis across four eukaryotic and six prokaryotic species. Specifically, sense codons within validated CDS regions were randomly selected and substituted with one of the three stop codons (TAA, TAG, or TGA). Genomic segments spanning 4,000 bp upstream and downstream of the mutation site were extracted. DeepD was employed to compute the per-site log-likelihood for both wild-type and mutant sequences. The $\Delta Likelihood$ per site was calculated as

$\frac{1}{N}\sum\left( \log P_{\mathrm{mut}}-\log P_{\mathrm{wt}} \right)$,

and where $N$ represents the sequence length. This metric quantifies the model’s sensitivity to the overall structural coherence of the sequence.

### 5 Biological interpretability of expert routing

#### 5.1 Expert feature retrieval from inference

The DeepD model replaces standard feed-forward layers with Mixture-of-Experts (MoE) layers, each containing $E=576$ expert sub-networks. For a given input token at layer $l$, the gating network computes a probability distribution $g^{(l)}\in R^{576}$ over all experts via a softmax function, from which the top-k experts are selected for activation. We extracted the full pre-selection gating weight vector for every token at each MoE layer (layers 2–24; layer 1 is a standard attention layer without MoE) and assembled these into a per-layer gating weight matrix $G^{(l)}\in R^{N\times576}$, where $N$ is the total number of tokens across all analyzed genomic regions. The procedure for constructing $G^{(l)}$ differed between model variants according to context window size.

For DeepD-60B (8K), inference was performed using a sliding window of 8,192 bp with a stride of 4,096 bp. To avoid representing the same genomic position multiple times in overlapping windows, only the gating weights corresponding to the last 4,096 tokens (the non-overlapping portion) were retained from each window; these slices were then concatenated along the

sequence axis within each region, and all regions were further concatenated to form $G^{(l)}$. Analyses were performed on 5 genomic regions in *E. coli* K-12 MG1655 (NCBI RefSeq NC_000913.3,^48^ covering replication origins, the *lac* operon, and CRISPR loci, etc.), 11 regions in human genome (NCBI RefSeq GCF_000001405.40), and 3 regions in maize B73 (NCBI RefSeq GCA_902167145.1) spanning known gene loci (*tb1*, *tga1*, *gt1*) (see Supplementary Table 4)^49-52^. For DeepD-60B (1M), each target genomic region fits within a single inference window, and no sliding-window segmentation was required. Each region, extended by 8,192 bp of upstream flanking sequence, was supplied as one contiguous input.

| ***E. coli*** | |  | |  | ***H. sapiens*** |  |  |
| --- | --- | --- | --- | --- | --- | --- | --- |
| Chrom. | Start | | End | | Chrom. | Start | End |
| U00096.3 | 360,000 | | | 369,000 | chr2 | 176,090,000 | 176,200,000 |
| U00096.3 | 2,877,250 | | | 2,879,000 | chr3 | 180,989,770 | 181,791,029 |
| ***Z. mays*** |  | | |  | chr7 | 27,040,000 | 27,180,000 |
| Chrom. | Start | | | End | chr8 | 127,700,000 | 127,750,000 |
| chr1 | 23,422,054 | | | 23,446,622 | chr11 | 5,200,000 | 5,240,000 |
| chr1 | 272,249,064 | | | 272,344,148 | chr11 | 31,775,000 | 31,825,000 |
| chr4 | 46,636,432 | | | 46,664,396 | chr12 | 6,510,000 | 6,560,000 |

**Supplementary Table 4** Genome regions used for DeepD inference.

#### 5.2 Genomic annotation labels

Each token was assigned to binary annotation labels by intersecting token coordinates with genomic feature intervals. For *E. coli*, gene structure annotations (CDS, intergenic, etc.) were obtained from the genome annotation file accompanying NCBI RefSeq NC_000913.3,^48^ and CRISPR array coordinates were retrieved from the CRISPR-Cas++ database (https://crisprcas.i2bc.paris-saclay.fr). For the human genome, gene structure annotations (exon, CDS, intron, etc.) were obtained from the NCBI RefSeq annotation file corresponding to GCF_000001405.40 (hg38), repetitive element annotations were derived from the RepeatMasker output included in the same NCBI annotation release. For maize, gene structure annotations and repetitive element annotations (including LINE, SINE, LTR, and DNA transposon families) were obtained from the annotation files associated with assembly GCA_902167145.1 (B73v5) deposited in MaizeGDB.^52,53^ Maize *cis*-regulatory elements were defined as accessible chromatin regions identified from published ATAC-seq data for B73 leaf,^52^ with peak calling performed using MACS v2.2.9.1 in narrow-peak mode.^54^

#### 5.3 Balanced subsampling and data splitting

Genomic annotations are inherently imbalanced in proportion. To prevent classifiers from biasing toward the majority class, balanced subsampling was applied prior to model training. For *E. coli*, the minority class size was determined, and an equal number of samples was drawn without replacement from the majority class to yield a 1:1 class ratio. For the human and maize genomes, a fixed-size stratified sampling strategy based on label combinations was applied. Specifically, a total of 70,000 tokens were sampled for the human dataset, comprising 10,000 tokens per target group and 30,000 non-exclusive (i.e., samples not exclusively assigned to any single target category, including unannotated regions or multi-label overlaps) tokens. For the maize dataset, a total of 55,000 tokens were sampled, consisting of 5,000 tokens per target group and 20,000 non-exclusive tokens, utilizing a priority sampling mode that favored CDS, UTR, and intron regions. The balanced dataset was then split into training (80%), test (10%) and validation (10%) subsets with stratification by label.

#### 5.4 Classifier training and evaluation

For each combination of MoE layer $l$ and annotation category, an XGBoost classifier was trained on the 576-dimensional gating weight vector to predict the corresponding annotation labels.^55^ For each combination of MoE layer $l$ (layers 2–24) and annotation category, we trained a binary classifier in a one-vs-rest setting using XGBClassifier from the Python xgboost package, with the 576-dimensional gating-weight vector as input. The classifiers were trained separately for the DeepD-60B (8K) and DeepD-60B (1M) using the following hyperparameters: n_estimators=5000, max_depth=3, learning_rate=0.12, min_child_weight=200, subsample=0.3, colsample_bytree=0.3, reg_alpha=30, reg_lambda=0.5, tree_method="hist", device="auto", random_state=42, and n_jobs=16. The gamma parameter was not specified, and early stopping was disabled. Model performance was evaluated using the AUPRC and log-loss on the test set. AUPRC was selected as the primary metric because it more faithfully reflects classifier performance under class imbalance than AUROC, particularly when the positive class is rare in the original genomic distribution. The layer achieving the highest test set AUPRC for each annotation category was designated the optimal extraction layer.

#### 5.5 Genome-wide prediction and feature importance

After training, each classifier was applied to all tokens in the original unsubsampled dataset, yielding token-level predicted probabilities. These continuous scores were used to generate genome-wide predicted annotation tracks for visualization along genomic coordinates. Gain-based feature importance scores were additionally extracted from each trained XGBoost model; gain importance quantifies the average reduction in loss achieved by splits on each expert feature across all trees, enabling identification of experts with disproportionate contributions to specific annotation predictions.

### 6 Controllable design of *cis*-regulatory elements in maize

#### 6.1 Data processing

For the maize system, we used a previously published dataset of 26,572 maize protoplast core promoter sequences together with their corresponding STARR-seq activity measurements.^56^ To construct the conditioning labels required for controllable generation, continuous promoter activities were discretized into three categories based on the empirical activity distribution. Specifically, sequences in the top 30% of the activity distribution (activity > −2.018) were assigned the label “High”, those in the bottom 30% (activity < −3.007) were assigned the label “Low”, and the remaining 40% were assigned the label “Med”.

#### 6.2 Fine-tuning

For conditional fine-tuning, each training example consisted of the maize species token, an activity label (“High”, “Med”, or “Low”), and the corresponding core promoter sequence. DeepD was fine-tuned using the AdamW optimizer for a total of 50 million tokens with a global batch size of 96. The learning rate ranged from 1e-05 to 1e-04, with a linear warmup over the first 2.5% of total training steps to the maximum learning rate, followed by gradual decay to the minimum learning rate to ensure stable convergence.

#### 6.3 Training a predictive model of promoter activity

To build an auxiliary predictive model for downstream sequence ranking, all promoter sequences were passed through the DeepD-60B (8K) model from the midtraining checkpoint, and the resulting 2,048-dimensional representations were extracted. The dataset was split into training, validation, and test sets at a ratio of 8:1:1. A regression model based on bidirectional convolutional neural networks (bidirectional CNNs) was then trained on these representations using AdamW with an initial learning rate of 1e-04.^57^ Training incorporated early stopping monitored by validation loss to prevent overfitting.

#### 6.4 Sequence design

After fine-tuning, promoter generation was performed using only the activity label (“High”, “Low”), as the prompt. To balance sequence diversity and generation quality, we used a stochastic sampling strategy with temperature = 1.0, together with top-k truncation (k = 4) and top-p sampling (p = 0.99). This procedure enabled activity-conditioned generation of maize core promoter candidates spanning the desired activity classes.

#### 6.5 Filtering and candidate selection

Generated sequences were subjected to a multi-step filtering procedure before experimental validation. First, we performed homology searches against the NCBI web BLASTn service against the NCBI Nucleotide collection (nr/nt) database,^58^ and removed sequences exceeding the predefined similarity thresholds (identity × coverage ≥ 0.6 or full-length homology ≥ 80%), thereby enriching for non-natural or sufficiently novel candidates. The remaining sequences were then scored using the trained regression model, and candidates were ranked by their predicted promoter activity. Finally, the top 100 and bottom 100 sequences were selected for *in vivo* maize protoplast validation experiments.

#### 6.6 Motif enrichment analysis

For motif analysis of the designed maize core promoters, we used MEME v4.11.2 to perform de novo motif discovery on the 100 High candidate sequences selected for in vivo validation. To correct for sequence-composition bias, a zero-order Markov background model was generated directly from the input sequences. MEME was run in ZOOPS (zero or one occurrence per sequence) mode, scanning both DNA strands for motifs with widths ranging from 6 to 12 bp, with a maximum of five motifs to be identified.^59^ The discovered sequence motifs were then annotated using TOMTOM v4.11.2^60^ by comparison against the JASPAR 2026 plant database (parameters: Pearson distance, minimum overlap = 5, E-value threshold < 10.0),^61^ to assess the enrichment and positional distribution of canonical regulatory elements, including the TATA box and Y patch.

#### 6.7 Latent space visualization

For visualization of the generated regulatory elements, we extracted sequence embeddings and projected them into two dimensions using uniform manifold approximation and projection (UMAP).^62^ A single global UMAP manifold was first fitted using embeddings from the full sequence pool to ensure a shared coordinate system across subsets. The high-activity and low-activity generated elements selected for validation (100 each) were then projected onto this global embedding space for comparative visualization. UMAP was run with 15 nearest neighbors, minimum distance = 0.1, and cosine distance.

#### 6.8 Maize protoplast isolation and transformation

Maize seeds of LH244 line were sown in soil and grown in a controlled growth chamber under a 16 h light / 8 h dark photoperiod at 25°C and 50% humidity. After uniform seedling emergence (approximately 3 cm above the soil surface), seedlings were transferred to complete darkness and grown for an additional 7–10 days at 25°C and 50% humidity. Protoplasts were isolated and transformed using a previously established PEG4000–Ca²⁺-mediated protocol.^63^ For each tested construct, 10 μg plasmid DNA was transfected into 1 × 10⁵ protoplasts. After transformation, protoplasts were incubated in the dark at 28°C for 12–16 h, collected by centrifugation at 100 × g (standard gravitational acceleration) for 3 min, and used for subsequent analyses.

#### 6.9 Dual-luciferase reporter assay

Firefly and Renilla luciferase activities were quantified using a SpectraMax i3x microplate reader (Molecular Devices) with the Dual-Luciferase Reporter Assay Kit (Vazyme, DL101), following the manufacturer’s instructions. Briefly, after incubation and collection of transformed protoplasts, samples were lysed by adding 100 μL of 1× cell lysis buffer according to the sample volume and incubated at room temperature for 15 min. An equal volume of firefly luciferase substrate was then added, and after 3 min of incubation, firefly luciferase activity was measured. Subsequently, an equal volume of Renilla luciferase substrate was added to the same sample, and Renilla luciferase activity was recorded.

### 7 Controllable long-context design of eukaryotic chromosomes

Beyond context-extension midtraining, we fine-tuned DeepD-60B for species-specific chromosome-scale sequence design, targeting *S. cerevisiae* chromosome III (316,620 bp) and *A. thaliana* chromosome 1 (~32.64 Mb). Fine-tuning warm-started from the DeepD-60B checkpoint and continued to model sequences at a 32,768-token context. During fine-tuning, we froze the MoE router (gating network) and updated only the attention and expert parameters, which stabilized expert routing while adapting the model to the target design distribution.

#### 7.1 Data preparation of long-sequence fine-tuning

To adapt DeepD for species-specific genome generation, we constructed separate fine-tuning datasets for *S. cerevisiae* and *A. thaliana*.

For yeast genome generation, genome assemblies were obtained from the near telomere-to-telomere *S. cerevisiae* pan-genome dataset containing 1,086 genomes.^64^ We followed the assembly-quality criterion used in the original study for gene-based pangenome construction and retained assemblies with a Merqury quality value (QV) greater than 40. This filter yielded 762 genome assemblies corresponding to 651 isolates. The selected set did not include S288C, which was used as the target backbone for yeast genome generation.

For *A. thaliana* genome generation, 406 candidate genome assemblies were collected from [NGDC](https://ngdc.cncb.ac.cn/) and project-hosted repositories associated with published *A. thaliana* genome-assembly studies, including the 69-accession pan-genome, the pan-genome and local-adaptation dataset, and the 27-genome comparison dataset.^65-67^ Of these assemblies, 141 were accompanied by matched gene annotations. For each assembly lacking a matched annotation, gene models were transferred from the TAIR12 reference annotation (GCA_978657495.1) using Liftoff v1.6.3 with the options -exclude_partial, -s 0.9, and -polish.^68^ Assemblies whose original or Liftoff-generated annotations did not contain explicit CDS features were excluded from subsequent transcript-level quality control.

For each remaining *A. thaliana* genome–annotation pair, CDS features were parsed from the annotation file and grouped by transcript. CDS segments belonging to the same transcript were ordered according to the annotated strand, extracted from the corresponding genome assembly, reverse-complemented when necessary, and concatenated to reconstruct the complete coding sequence. A transcript was considered valid if its reconstructed CDS began with an ATG start codon, ended with an in-frame stop codon (TAA, TAG, or TGA), had a total length divisible by three, and contained no premature in-frame stop codons. For each genome, the CDS pass rate was calculated as the proportion of CDS-containing transcripts satisfying all four criteria. Genomes with a CDS pass rate greater than 90% were retained. Accessions belonging to the Columbia/Col-0 genetic background were then removed to reduce redundancy with the reference background, followed by the exclusion of assemblies that were not resolved at chromosome scale. After these filtering steps, 255 *A. thaliana* genomes were retained for fine-tuning the *A. thaliana* generation model.

As a result, 762 *S. cerevisiae* genomes and 255 *A. thaliana* genomes were obtained. We encoded genomic annotations for these genomes using the same method applied to the pretraining data. Please refer to Methods section 2.2 for more details.

#### 7.2 The Context–Anchor–Body (CAB) formulation

We cast long-sequence design as a fill-in-the-middle (FIM) problem under a Context–Anchor–Body (CAB) layout, which was compatible with strictly left-to-right autoregressive decoding.^69^ This followed the Suffix–Prefix–Middle (SPM) formulation of FIM training, in which the suffix was placed first, the prefix second, and the middle segment was generated autoregressively. Each training window was decomposed into three roles: a downstream flanking Context (C, 8–16 kb; the FIM suffix), a core regulatory/promoter Anchor (A, 8–16 kb; the FIM prefix), and the target Body (B, 8 kb; the FIM middle) to be generated. The Context and Anchor segments were reordered ahead of the Body and delimited by dedicated separator tokens ([suf], [sop]), so that the model conditions on both the surrounding genomic background and the core regulatory element before emitting the target region. Training samples were drawn by cropping 32 kb windows with an 8 kb overlap from source sequences, and the CAB reordering was applied stochastically at a rate of 0.5. The Context and Anchor acted purely as conditioning prefixes and were excluded from the loss, so gradients were computed only on the Body.

#### 7.3 Region-aware perturbation and loss reweighting

To learn robust topological structure without sacrificing fidelity in functional elements, we applied a region-aware combination of input perturbation and loss weighting within the generated region. Gene (coding) regions received conservative treatment, 2% input perturbation with loss weight 1.0, to protect core sequence grammar, whereas intergenic spacer regions received aggressive 15% perturbation with a reduced loss weight of 0.1, which kept the model from entirely abandoning these noisier positions while still encouraging it to prioritize long-range layout. Conditioning segments (C and A) carried zero loss weight. To avoid corrupting reading frames, perturbation respected a codon-protection length of 3 nucleotides.

#### 7.4 Fine-tuning configuration

Fine-tuning inherited the DeepD-60B architecture and was optimized with Adam ($\beta_{1}=0.9, \beta_{2}=0.95$, weight decay 0.1) under a cosine schedule (peak learning rate $2.5 \times{10}^{-5}$, minimum $2.5 \times{10}^{-6}$, 2.5% warmup, gradient clipping 1.0) in bf16 precision with FlashAttention-2 and gated attention, for 20B tokens. Key hyperparameters are summarized in Supplementary Table 5.

| Setting | Value |
| --- | --- |
| Fine-tuning context length | 32,768 |
| FIM/CAB application rate | 0.5 |
| Context (C) length | 8–16k bp |
| Anchor (A) / Body (B) length | 8–16k bp / 8k bp |
| Gene-region perturbation / loss weight | 2%/ 1.0 |
| Spacer-region perturbation / loss weight | 15% / 0.1 |
| Learning rate (min) | 2.5 × 10^-5^(2.5 × 10^-6^) |
| LR schedule / warmup | cosine / 2.5% |
| Fine-tuning tokens | 20B |

**Supplementary Table 5** Long-sequence fine-tuning configuration for DeepD-60B. Optimization uses Adam ($\beta_{1}=0.9, \beta_{2}=0.95$) with bf16 precision, FlashAttention-2, and gated attention.

#### 7.5 Generation strategy: sliding-window relay

At inference, DeepD generates long contigs using the same CAB arrangement. The sequence was produced in fixed-size Body chunks (8 kb per round): after each round the model slides forward by one chunk, taking the tail of the freshly generated sequence as the Anchor (and, when available, downstream reference as Context) for the next round, and repeated until the requested length was reached. This design was motivated by streaming-generation practice, where retaining a compact set of anchoring tokens together with a rolling window of the most recent sequence maintains stable long-range decoding.^70^ In our setting the Anchor plays this anchoring role, providing a stable conditioning prefix for each round while the window advances. The relay mode decouples the total output length from the training context and enables generation of contigs far longer than any single window. Decoding uses nucleus/top-$k$ sampling (temperature 1.0, top-$k$ 4, top-$p$ 0.99).

| Setting | Value |
| --- | --- |
| Generation mode | sliding-window relay |
| Anchor (A) length | 8,192 bp |
| Body (B) chunk length | 8,192 bp |
| Context (C) length | 16,384 bp (0 = *de novo*) |
| Temperature | 1.0 |
| Top-$\boldsymbol{k}$/Top-$\boldsymbol{p}$ | 4 / 0.99 |

**Supplementary Table 6** Long-sequence generation (CAB rolling-FIM) settings.

#### 7.6 Genome annotation of generated *S. cerevisiae* sequences

Generated sequences were converted from raw text to FASTA format. Each generated sequence was annotated using a standardized pipeline. RepeatMasker was run with the Repbase library using NCBI mode (-e ncbi -xsmall) to soft-mask repetitive elements.^10,71^ Transfer RNA (tRNA) genes were annotated using tRNAscan-SE v2.0.12 with eukaryotic parameters (-E).^72^ Ribosomal RNA (rRNA) genes were annotated using barrnap v0.9 (https://github.com/tseemann/barrnap) with the eukaryotic kingdom setting (--kingdom euk).

*De novo* protein-coding gene prediction was performed using BRAKER4 v0.5.0-beta, a Snakemake-based reimplementation of the BRAKER workflow (Krall and Hoff, BRAKER4 GitHub repository; commit c72618baa4631db4a4128391a40c8ba7392df17c).^73^ Because the OrthoDB v12 protein database was supplied as the sole source of extrinsic evidence,^74^ BRAKER4 automatically selected EP mode, which implements the protein-supported BRAKER2 workflow. In this mode, ProtHint v2.6.0 was run iteratively to generate and refine protein-derived hints, followed by GeneMark-EP+ training and AUGUSTUS v3.5.0 prediction using the protein evidence.^75,76^ The BRAKER2 publication was cited for the EP-mode workflow, whereas the BRAKER4 repository was cited for the Snakemake implementation used in this study.

#### 7.7 Functional annotation of generated *S. cerevisiae* sequences

Predicted protein sequences were searched against the Pfam database using DIAMOND BLASTp v2.1.10 with default parameters.^77,78^ Hits were filtered to retain only high-confidence matches with >80% sequence identity, >80% query coverage, and >80% target coverage. Filtered proteins were further subjected to InterProScan v5.76 for comprehensive protein domain and functional site annotation.^79^

#### 7.8 Synteny analysis of generated *S. cerevisiae* sequences

Synteny analysis was conducted to evaluate the conservation of gene order (collinearity) between the generated full-length *S. cerevisiae* sequences and the corresponding reference chromosome. To ensure robust homologous anchor identification, collinearity was assessed at the protein level.

First, protein sequences and their corresponding genomic coordinates were extracted from the gene annotation (GFF3) files of both the generated sequences and the reference genomes. To identify homologous gene pairs, an all-vs-all sequence similarity search was performed between the generated and reference protein sets using DIAMOND BLASTp with a maximum E-value threshold of 1e-5.

The resulting homologous matches and their chromosomal coordinates were formatted and used as input for DAGchainer to identify collinear blocks.^80^ The software was configured to identify both collinear and inverted syntenic regions while ignoring tandem duplications. Synteny blocks were defined by requiring a minimum of 5 aligned gene pairs per block, allowing a maximum distance of 1,000,000 bp between adjacent matches, and applying a gap parameter of 40,000 bp.

Finally, the retained collinearity blocks were visualized as dot plots and ribbon diagrams to illustrate the degree of syntenic conservation and the distribution of conserved genomic segments across the chromosomes.

#### 7.9 Genomic diversity analysis of generated *S. cerevisiae* sequences

Target chromosome III sequences were extracted from each *S. cerevisiae* assembly using sample-specific sequence identifiers or regular-expression rules. Chromosome identifiers in the reference FASTA, query FASTA, and reference GFF3 files were standardized before alignment.

Pairwise reference–query alignments were generated with AnchorWave v1.2.6.^81^ Reference CDS sequences were extracted using anchorwave gff2seq, excluding CDS exons shorter than 20 bp, and aligned to the reference and query chromosomes with minimap2 v2.28 using -x splice -k 12 -a -p 0.4 -N 20 -t 8.^82^ AnchorWave genoAli was run with -m 20 -mi 0 -mi2 0 -B -4 -O1 -4 -E1 -2 -O2 -80 -E2 -1 -t 16. The parameters -w 100000 -fa3 100000 were used for *S. cerevisiae*. Pairwise MAF alignments were converted into sample-level assembly-derived gVCFs using the TASSEL v5.2.96 MAFToGVCFPlugin.^83^

The TASSEL gVCFs were normalized to the reference chromosome identifier and consolidated using GenomicsDBImport and GenotypeGVCFs in GATK v4.6.1.0.^84^ These tools were used for cohort-level consolidation of deterministic assembly-derived genotypes rather than read-based genotype inference. The resulting multisample VCF was normalized against the reference FASTA using BCFtools v1.24.^85^ Multiallelic records were decomposed and left-normalized using bcftools norm -f reference.fa -m -both, followed by coordinate sorting and indexing. SNPs were defined as biallelic records with single-nucleotide REF and ALT alleles and at least one non-reference genotype among the analyzed samples.

Site-level AC, AN, AF, MAF, and F_MISSING were recalculated with BCFtools +fill-tags using all analyzed chromosomes. Sites with MAF <0.05 or F_MISSING >0.50 were removed. Read-depth filtering was not applied because genotype states were inferred from assembled chromosome sequences rather than sequencing reads.

Insertions and deletions of at least 50 bp were identified independently from each reference–query alignment using the TASSEL gVCF and associated assembly-coordinate annotations. Candidate records were initially selected when the REF allele or at least one non-symbolic ALT allele was ≥50 bp. Reference intervals were derived from VCF coordinates and REF-allele lengths, whereas query intervals and alignment orientation were obtained from TASSEL assembly-coordinate fields. Adjacent collinear blocks on the same reference chromosome, query chromosome, and strand were merged. Merged blocks with reference spans <50 bp and query spans ≥50 bp were classified as insertions, whereas blocks with reference spans ≥50 bp and query spans <50 bp were classified as deletions. Events with |SVLEN| < 50 bp were discarded.

Per-sample insertion and deletion VCFs were merged separately with SURVIVOR v1.0.7 using parameters “merge <vcf_list> 20 1 1 1 0 50”.^86^ Events with breakpoints within 20 bp were eligible for clustering when SV type and strand agreed; at least one input callset was required, SV-length-dependent distance scaling was disabled, and the minimum SV length was 50 bp. The merged insertion and deletion sets were combined, coordinate-sorted, and reordered according to the sample manifest.

SNP effects were annotated with snpEff v5.4c using a run-specific database constructed from the normalized reference FASTA, GFF3 annotation, and CDS FASTA used for AnchorWave alignment.^87^ The database was built using snpEff build -gff3 -v. The population-filtered biallelic SNP VCF was annotated using snpEff ann -v. A SNP was classified as synonymous if at least one ANN entry for its ALT allele contained synonymous_variant. Only synonymous SNPs were used for phylogenetic inference.

#### 7.10 Reference-guided annotation and gene-model evaluation of generated *A. thaliana* sequences

The published GFF3 annotation of the TAIR12 Col-0 reference genome was used as the gene-model baseline. Reference annotations were transferred to each generated sequence using Liftoff with the parameters -a 0.50 -s 0.50 -copies -sc 0.90 -polish -flank 0 -overlap 0.10 -d 2.0.^68^ For each protein-coding gene, the transcript with the greatest total CDS length was selected as the representative transcript. Its CDS segments were ordered according to the annotated strand, concatenated, and translated.

CDS structure quality was assessed from the translatability of each concatenated CDS and the properties of its translated product. A model failed CDS structure assessment if its translation contained an internal stop codon, a dot character, or an unsupported amino-acid character. Empty sequences and translations shorter than 1 aa were also excluded. When a model failed CDS structure assessment, the entire gene model—including the gene, representative transcript, and associated CDS features—was excluded from annotation-dependent summaries, Pfam analysis, and protein alignment.

Gene correspondence between the reference and generated sequences was assessed at both the DNA and protein levels. Annotation-dependent DNA comparisons were performed using minimap2.^82^ Candidate alignments were required to have local sequence identity ≥0.50 and both generated- and reference-sequence coverage ≥0.30. Local identity was calculated as the number of matching bases divided by the alignment-block length, and sequence coverage was calculated as the aligned length divided by the full length of the corresponding sequence.

Protein-level homology searches were performed using DIAMOND BLASTp with an E-value threshold of 1e-5.^78^ Candidate pairs were required to have protein identity, query coverage, and reference coverage ≥0.70. DAGchainer was subsequently run with -D 100000 -g 10000 -A 8 to identify candidate relationships supported by gene collinearity.^80^ Final matching was constrained to be one-to-one. Reciprocal best hits were selected first, after which remaining candidates were assigned in descending order of alignment score, matched length, identity, and bidirectional coverage. Thus, each reference gene and each generated gene occurred in no more than one matched pair.

The core-gene set was obtained from the publicly available orthogroup dataset accompanying the pan-genome study of 69 *A. thaliana* accessions.^67^ Orthogroups present in all 69 accessions were considered core gene families. Col-0 members of these families were selected, and their representative CDS sequences were extracted from the corresponding annotation to construct a local dataset containing core ORFs.

Recovery of core ORFs in generated DNA was evaluated using minimap2 with -x splice:hq -uf --secondary=yes -N 100 -p 0.1 -c --eqx --cs=long. A core ORF was classified as DNA recovered when at least one individual alignment within its expected target interval had local identity ≥0.90 and ORF coverage ≥0.90. Partial alignments from different sequence records or distinct genomic loci were not merged. Each core ORF was subsequently aligned against the concatenated CDS sequences of generated annotation models. An annotation was classified as complete when ORF-to-CDS identity, core-ORF coverage, and predicted-CDS coverage were each ≥0.90; the ORF-to-CDS alignment was forward; the predicted-model strand agreed with the genomic DNA-alignment strand; and the recovered DNA locus covered ≥0.90 of the union of the predicted CDS intervals. Strict core-gene recovery therefore required DNA recovery, complete annotation, and a passing CDS structure. A generated gene inherited the core label only when it formed a one-to-one match with a reference core gene.

Pfam-domain support was evaluated by searching the Pfam database^77^ using DIAMOND BLASTp with an E-value threshold of 1e-3. A gene with at least one retained hit was classified as Pfam-supported. When multiple hits were detected, the hit with the lowest E-value and then the highest bit score was selected.

#### 7.11 Pairwise alignment, variant detection, and AIAtRegion1 sequence-divergence analysis in *A. thaliana*

The TAIR12 Col-0 chromosome 1 sequence and its corresponding GFF3 annotation were used as the reference. Pairwise reference–query genome alignments for natural and generated materials were produced using AnchorWave v1.2.6.^81^ Reference CDS sequences were first mapped to each query genome using minimap2 in splice mode with the principal parameters -x splice -k 12 -p 0.4 -N 20. Complete chromosome 1 assemblies from natural accessions were aligned using the AnchorWave genoAli mode, whereas generated sequences containing only the target region were aligned using the proali -R 1 -Q 1 mode. The principal AnchorWave parameters were -m 20 -mi 0 -mi2 0 -w 38000 -fa3 200000, with scoring parameters -B -4 -O1 -4 -E1 -2 -O2 -80 -E2 -1. A pairwise MAF file relative to the reference genome was generated for each material.

Each pairwise MAF file was converted into a single-sample gVCF using the TASSEL 5 MAFToGVCFPlugin with -fillGaps false to prevent unaligned regions from being interpreted as homozygous reference genotypes.^83^ Single-sample gVCFs were imported using GenomicsDBImport and jointly genotyped as diploid samples using GenotypeGVCFs in GATK v4.6.1.0.^84^ The joint VCF was normalized against the reference genome, decomposed at multiallelic sites, sorted, compressed, and indexed using bcftools.^85^ After normalization, records with single-nucleotide REF and ALT alleles were classified as SNPs. Repeat filtering excluded only SNPs overlapping GFF3 features annotated as CENTROMERIC_REPEAT.

Structural variants were extracted independently for each material from its pairwise gVCF together with the reference and query FASTA sequences. Insertions and deletions were identified according to the number of affected bases, with a minimum SV length of 50 bp. Adjacent variant blocks separated by no more than 20 bp were merged, and a 10-kb N-gap inspection window was used to exclude events potentially caused by assembly gaps. Complex replacement alleles were processed using the both policy, under which their insertion and deletion components were retained separately. Per-sample insertion and deletion VCFs were merged separately using SURVIVOR^86^ with a maximum breakpoint distance of 20 bp and a minimum supporting-sample count of one, while requiring concordant SV type and strand. The merged insertion and deletion callsets were subsequently combined into a population-level SV VCF.

The filtered SNP VCF was functionally annotated using snpEff.^87^ A run-specific snpEff database was constructed from the same reference FASTA and GFF3 annotation. This maintained a common reference-coordinate system across sequence alignment, variant localization, and functional annotation.

Sequence-divergence analysis in the main text was restricted to AIAtRegion1 (chr1:134500–1134500; 0-based, half-open). For natural accessions, the region 1 alignment was extracted from the complete chromosome 1 MAF according to reference coordinates. Reference and query sequences were compared column by column. Columns containing N in either sequence and columns containing gaps in both sequences were ignored. A reference gap was counted as an inserted base, a query gap as a deleted base, unequal non-gap nucleotides as SNPs, and identical nucleotides as matches. Insertions and deletions were counted by the number of affected bases rather than by the number of variant events. Sequence divergence was calculated as:

$$D = \frac{N_{\mathrm{SNP}} + N_{\mathrm{INS}} + N_{\mathrm{DEL}}}{N_{\mathrm{MATCH}} + N_{\mathrm{SNP}} + N_{\mathrm{INS}} + N_{\mathrm{DEL}}}.$$

### References

1.	Qiu, Z., Wang, Z., Zheng, B., Huang, Z., Wen, K., Yang, S., Men, R., Yu, L., Huang, F., and Huang, S. (2026). Gated attention for large language models: Non-linearity, sparsity, and attention-sink-free. Advances in Neural Information Processing Systems *38*, 100092-100118.

2.	Peng, B., Quesnelle, J., Fan, H., and Shippole, E. (2024). Yarn: Efficient context window extension of large language models. International Conference on Learning Representations, 31932-31951.

3.	Dai, D., Deng, C., Zhao, C., Xu, R., Gao, H., Chen, D., Li, J., Zeng, W., Yu, X., and Wu, Y. (2024). Deepseekmoe: Towards ultimate expert specialization in mixture-of-experts language models. Proceedings of the 62nd Annual Meeting of the Association for Computational Linguistics (Volume 1: Long Papers), 1280-1297.

4.	Jacobs, S.A., Tanaka, M., Zhang, C., Zhang, M., Aminadabi, R.Y., Song, S.L., Rajbhandari, S., and He, Y. (2024). System optimizations for enabling training of extreme long sequence transformer models. Proceedings of the 43rd ACM Symposium on Principles of Distributed Computing, 121-130.

5.	Kingma, D.P., and Ba, J. (2014). Adam: A Method for Stochastic Optimization. CoRR *abs/1412.6980*.

6.	Kwon, W., Li, Z., Zhuang, S., Sheng, Y., Zheng, L., Yu, C.H., Gonzalez, J., Zhang, H., and Stoica, I. (2023). Efficient memory management for large language model serving with pagedattention. Proceedings of the 29th symposium on operating systems principles, 611-626.

7.	Rees, J.A., and Cranston, K. (2017). Automated assembly of a reference taxonomy for phylogenetic data synthesis. Biodiversity Data Journal *5*, e12581.

8.	Parks, D.H., Chuvochina, M., Rinke, C., Mussig, A.J., Chaumeil, P.-A., and Hugenholtz, P. (2022). GTDB: an ongoing census of bacterial and archaeal diversity through a phylogenetically consistent, rank normalized and complete genome-based taxonomy. Nucleic Acids Research *50*, D785-D794. <https://doi.org/10.1093/nar/gkab776>.

9.	Chklovski, A., Parks, D.H., Woodcroft, B.J., and Tyson, G.W. (2023). CheckM2: a rapid, scalable and accurate tool for assessing microbial genome quality using machine learning. Nature Methods *20*, 1203-1212. <https://doi.org/10.1038/s41592-023-01940-w>.

10.	Tarailo-Graovac, M., and Chen, N. (2009). Using RepeatMasker to Identify Repetitive Elements in Genomic Sequences. Current Protocols in Bioinformatics *25*, 4.10.11-14.10.14. <https://doi.org/10.1002/0471250953.bi0410s25>.

11.	Spengler, M.V., and Mettes, P. (2025). Low-distortion and GPU-compatible Tree Embeddings in Hyperbolic Space. In S. Aarti, F. Maryam, H. Daniel, L.-J. Simon, B. Felix, M. Tegan, W. Kiri, and Z. Jerry, eds. Proceedings of the 42nd International Conference on Machine Learning.

12.	Brixi, G., Durrant, M.G., Ku, J., Naghipourfar, M., Poli, M., Sun, G., Brockman, G., Chang, D., Fanton, A., Gonzalez, G.A., et al. (2026). Genome modelling and design across all domains of life with Evo 2. Nature *652*, 1349-1361. <https://doi.org/10.1038/s41586-026-10176-5>.

13.	Nguyen, E., Poli, M., Durrant, M.G., Kang, B., Katrekar, D., Li, D.B., Bartie, L.J., Thomas, A.W., King, S.H., Brixi, G., et al. (2024). Sequence modeling and design from molecular to genome scale with Evo. Science *386*, eado9336. <https://doi.org/10.1126/science.ado9336>.

14.	Avsec, Ž., Latysheva, N., Cheng, J., Novati, G., Taylor, K.R., Ward, T., Bycroft, C., Nicolaisen, L., Arvaniti, E., Pan, J., et al. (2026). Advancing regulatory variant effect prediction with AlphaGenome. Nature *649*, 1206-1218. <https://doi.org/10.1038/s41586-025-10014-0>.

15.	Dalla-Torre, H., Gonzalez, L., Mendoza-Revilla, J., Lopez Carranza, N., Grzywaczewski, A.H., Oteri, F., Dallago, C., Trop, E., de Almeida, B.P., Sirelkhatim, H., et al. (2025). Nucleotide Transformer: building and evaluating robust foundation models for human genomics. Nature Methods *22*, 287-297. <https://doi.org/10.1038/s41592-024-02523-z>.

16.	Notin, P., Kollasch, A.W., Ritter, D., van Niekerk, L., Paul, S., Spinner, H., Rollins, N., Shaw, A., Weitzman, R., Frazer, J., et al. (2023). ProteinGym: Large-Scale Benchmarks for Protein Design and Fitness Prediction. bioRxiv, 2023.2012.2007.570727. <https://doi.org/10.1101/2023.12.07.570727>.

17.	Firnberg, E., Labonte, J.W., Gray, J.J., and Ostermeier, M. (2014). A Comprehensive, High-Resolution Map of a Gene’s Fitness Landscape. Molecular Biology and Evolution *31*, 1581-1592. <https://doi.org/10.1093/molbev/msu081>.

18.	Jacquier, H., Birgy, A., Le Nagard, H., Mechulam, Y., Schmitt, E., Glodt, J., Bercot, B., Petit, E., Poulain, J., Barnaud, G., et al. (2013). Capturing the mutational landscape of the beta-lactamase TEM-1. Proceedings of the National Academy of Sciences *110*, 13067-13072. <https://doi.org/10.1073/pnas.1215206110>.

19.	Kelsic, E.D., Chung, H., Cohen, N., Park, J., Wang, H.H., and Kishony, R. (2016). RNA Structural Determinants of Optimal Codons Revealed by MAGE-Seq. Cell Systems *3*, 563-571.e566. <https://doi.org/10.1016/j.cels.2016.11.004>.

20.	Weeks, R., and Ostermeier, M. (2023). Fitness and Functional Landscapes of the E. coli RNase III Gene rnc. Molecular Biology and Evolution *40*, msad047. <https://doi.org/10.1093/molbev/msad047>.

21.	Rockah-Shmuel, L., Tóth-Petróczy, Á., and Tawfik, D.S. (2015). Systematic Mapping of Protein Mutational Space by Prolonged Drift Reveals the Deleterious Effects of Seemingly Neutral Mutations. PLOS Computational Biology *11*, e1004421. <https://doi.org/10.1371/journal.pcbi.1004421>.

22.	Chen, J.Z., Fowler, D.M., and Tokuriki, N. (2020). Comprehensive exploration of the translocation, stability and substrate recognition requirements in VIM-2 lactamase. eLife *9*, e56707. <https://doi.org/10.7554/eLife.56707>.

23.	Melamed, D., Young, D.L., Gamble, C.E., Miller, C.R., and Fields, S. (2013). Deep mutational scanning of an RRM domain of the Saccharomyces cerevisiae poly(A)-binding protein. RNA *19*, 1537-1551. <https://doi.org/10.1261/rna.040709.113>.

24.	Faure, A.J., Domingo, J., Schmiedel, J.M., Hidalgo-Carcedo, C., Diss, G., and Lehner, B. (2022). Mapping the energetic and allosteric landscapes of protein binding domains. Nature *604*, 175-183. <https://doi.org/10.1038/s41586-022-04586-4>.

25.	Sun, S., Weile, J., Verby, M., Wu, Y., Wang, Y., Cote, A.G., Fotiadou, I., Kitaygorodsky, J., Vidal, M., Rine, J., et al. (2020). A proactive genotype-to-patient-phenotype map for cystathionine beta-synthase. Genome Medicine *12*, 13. <https://doi.org/10.1186/s13073-020-0711-1>.

26.	Tsuboyama, K., Dauparas, J., Chen, J., Laine, E., Mohseni Behbahani, Y., Weinstein, J.J., Mangan, N.M., Ovchinnikov, S., and Rocklin, G.J. (2023). Mega-scale experimental analysis of protein folding stability in biology and design. Nature *620*, 434-444. <https://doi.org/10.1038/s41586-023-06328-6>.

27.	Rives, A., Meier, J., Sercu, T., Goyal, S., Lin, Z., Liu, J., Guo, D., Ott, M., Zitnick, C.L., Ma, J., and Fergus, R. (2021). Biological structure and function emerge from scaling unsupervised learning to 250 million protein sequences. Proceedings of the National Academy of Sciences *118*, e2016239118. <https://doi.org/10.1073/pnas.2016239118>.

28.	Hayes, T., Rao, R., Akin, H., Sofroniew, N.J., Oktay, D., Lin, Z., Verkuil, R., Tran, V.Q., Deaton, J., Wiggert, M., et al. (2025). Simulating 500 million years of evolution with a language model. Science *387*, 850-858. <https://doi.org/10.1126/science.ads0018>.

29.	Madani, A., Krause, B., Greene, E.R., Subramanian, S., Mohr, B.P., Holton, J.M., Olmos, J.L., Xiong, C., Sun, Z.Z., Socher, R., et al. (2023). Large language models generate functional protein sequences across diverse families. Nature Biotechnology *41*, 1099-1106. <https://doi.org/10.1038/s41587-022-01618-2>.

30.	Nijkamp, E., Ruffolo, J.A., Weinstein, E.N., Naik, N., and Madani, A. (2023). ProGen2: Exploring the boundaries of protein language models. Cell Systems *14*, 968-978.e963. <https://doi.org/10.1016/j.cels.2023.10.002>.

31.	Yang, K.K., Fusi, N., and Lu, A.X. (2024). Convolutions are competitive with transformers for protein sequence pretraining. Cell Systems *15*, 286-294.e282. <https://doi.org/10.1016/j.cels.2024.01.008>.

32.	Zhang, Z.D., Nayar, M., Ammons, D., Rampersad, J., and Fox, G.E. (2009). Rapid in vivo exploration of a 5S rRNA neutral network. Journal of Microbiological Methods *76*, 181-187. <https://doi.org/10.1016/j.mimet.2008.10.010>.

33.	Pitt, J.N., and Ferré-D’Amaré, A.R. (2010). Rapid Construction of Empirical RNA Fitness Landscapes. Science *330*, 376-379. <https://doi.org/10.1126/science.1192001>.

34.	Hayden, E.J., Ferrada, E., and Wagner, A. (2011). Cryptic genetic variation promotes rapid evolutionary adaptation in an RNA enzyme. Nature *474*, 92-95. <https://doi.org/10.1038/nature10083>.

35.	Guy, M.P., Young, D.L., Payea, M.J., Zhang, X.J., Kon, Y., Dean, K.M., Grayhack, E.J., Mathews, D.H., Fields, S., and Phizicky, E.M. (2014). Identification of the determinants of tRNA function and susceptibility to rapid tRNA decay by high-throughput in vivo analysis. GENES & DEVELOPMENT *28*, 1721-1732. <https://doi.org/10.1101/gad.245936.114>.

36.	Kobori, S., Nomura, Y., Miu, A., and Yokobayashi, Y. (2015). High-throughput assay and engineering of self-cleaving ribozymes by sequencing. Nucleic Acids Research *43*, e85-e85. <https://doi.org/10.1093/nar/gkv265>.

37.	Domingo, J., Diss, G., and Lehner, B. (2018). Pairwise and higher-order genetic interactions during the evolution of a tRNA. Nature *558*, 117-121. <https://doi.org/10.1038/s41586-018-0170-7>.

38.	Andreasson, J.O.L., Savinov, A., Block, S.M., and Greenleaf, W.J. (2020). Comprehensive sequence-to-function mapping of cofactor-dependent RNA catalysis in the glmS ribozyme. Nature Communications *11*, 1663. <https://doi.org/10.1038/s41467-020-15540-1>.

39.	Penić, R.J., Vlašić, T., Huber, R.G., Wan, Y., and Šikić, M. (2025). RiNALMo: general-purpose RNA language models can generalize well on structure prediction tasks. Nature Communications *16*, 5671. <https://doi.org/10.1038/s41467-025-60872-5>.

40.	Zhou, Z., Ji, Y., Li, W., Dutta, P., Davuluri, R., and Liu, H. (2024). DNABERT-2: Efficient Foundation Model and Benchmark For Multi-Species Genomes. In B. Kim, Y. Yue, S. Chaudhuri, K. Fragkiadaki, M. Khan, and Y. Sun, eds. International Conference on Learning Representations 2024 (ICLR 2024).

41.	Landrum, M.J., Lee, J.M., Benson, M., Brown, G.R., Chao, C., Chitipiralla, S., Gu, B., Hart, J., Hoffman, D., Jang, W., et al. (2018). ClinVar: improving access to variant interpretations and supporting evidence. Nucleic Acids Research *46*, D1062-D1067. <https://doi.org/10.1093/nar/gkx1153>.

42.	Sullivan, P.J., Quinn, J.M.W., Wu, W., Pinese, M., and Cowley, M.J. (2024). SpliceVarDB: A comprehensive database of experimentally validated human splicing variants. The American Journal of Human Genetics *111*, 2164-2175. <https://doi.org/10.1016/j.ajhg.2024.08.002>.

43.	Findlay, G.M., Daza, R.M., Martin, B., Zhang, M.D., Leith, A.P., Gasperini, M., Janizek, J.D., Huang, X., Starita, L.M., and Shendure, J. (2018). Accurate classification of BRCA1 variants with saturation genome editing. Nature *562*, 217-222. <https://doi.org/10.1038/s41586-018-0461-z>.

44.	Huang, H., Hu, C., Na, J., Hart, S.N., Gnanaolivu, R.D., Abozaid, M., Rao, T., Tecleab, Y.A., Ambrosone, C.B., Yao, S., et al. (2025). Functional evaluation and clinical classification of BRCA2 variants. Nature *638*, 528-537. <https://doi.org/10.1038/s41586-024-08388-8>.

45.	Engel, S.R., Dietrich, F.S., Fisk, D.G., Binkley, G., Balakrishnan, R., Costanzo, M.C., Dwight, S.S., Hitz, B.C., Karra, K., Nash, R.S., et al. (2014). The Reference Genome Sequence of Saccharomyces cerevisiae: Then and Now. G3 Genes|Genomes|Genetics *4*, 389-398. <https://doi.org/10.1534/g3.113.008995>.

46.	Hyatt, D., Chen, G.-L., LoCascio, P.F., Land, M.L., Larimer, F.W., and Hauser, L.J. (2010). Prodigal: prokaryotic gene recognition and translation initiation site identification. BMC Bioinformatics *11*, 119. <https://doi.org/10.1186/1471-2105-11-119>.

47.	Sharp, P.M., and Li, W.-H. (1987). The codon adaptation index-a measure of directional synonymous codon usage bias, and its potential applications. Nucleic Acids Research *15*, 1281-1295. <https://doi.org/10.1093/nar/15.3.1281>.

48.	Riley, M., Abe, T., Arnaud, M.B., Berlyn, M.K.B., Blattner, F.R., Chaudhuri, R.R., Glasner, J.D., Horiuchi, T., Keseler, I.M., Kosuge, T., et al. (2006). Escherichia coli K-12: a cooperatively developed annotation snapshot—2005. Nucleic Acids Research *34*, 1-9. <https://doi.org/10.1093/nar/gkj405>.

49.	Studer, A., Zhao, Q., Ross-Ibarra, J., and Doebley, J. (2011). Identification of a functional transposon insertion in the maize domestication gene tb1. Nature Genetics *43*, 1160-1163. <https://doi.org/10.1038/ng.942>.

50.	Wills, D.M., Whipple, C.J., Takuno, S., Kursel, L.E., Shannon, L.M., Ross-Ibarra, J., and Doebley, J.F. (2013). From Many, One: Genetic Control of Prolificacy during Maize Domestication. PLOS Genetics *9*, e1003604. <https://doi.org/10.1371/journal.pgen.1003604>.

51.	Wang, H., Studer, A.J., Zhao, Q., Meeley, R., and Doebley, J.F. (2015). Evidence That the Origin of Naked Kernels During Maize Domestication Was Caused by a Single Amino Acid Substitution in tga1. Genetics *200*, 965-974. <https://doi.org/10.1534/genetics.115.175752>.

52.	Hufford, M.B., Seetharam, A.S., Woodhouse, M.R., Chougule, K.M., Ou, S., Liu, J., Ricci, W.A., Guo, T., Olson, A., Qiu, Y., et al. (2021). De novo assembly, annotation, and comparative analysis of 26 diverse maize genomes. Science *373*, 655-662. <https://doi.org/10.1126/science.abg5289>.

53.	Woodhouse, M.R., Cannon, E.K., Portwood, J.L., 2nd, Gardiner, J.M., Hayford, R.K., Haley, O., and Andorf, C.M. (2025). Tools and Resources at the Maize Genetics and Genomics Database (MaizeGDB). Cold Spring Harb Protoc *2025*, pdb.over108430. <https://doi.org/10.1101/pdb.over108430>.

54.	Feng, J., Liu, T., Qin, B., Zhang, Y., and Liu, X.S. (2012). Identifying ChIP-seq enrichment using MACS. Nature Protocols *7*, 1728-1740. <https://doi.org/10.1038/nprot.2012.101>.

55.	Chen, T., and Guestrin, C. (2016). XGBoost: A scalable tree boosting system. Proceedings of the 22nd ACM SIGKDD International Conference on Knowledge Discovery and Data Mining. ACM.

56.	Jores, T., Tonnies, J., Wrightsman, T., Buckler, E.S., Cuperus, J.T., Fields, S., and Queitsch, C. (2021). Synthetic promoter designs enabled by a comprehensive analysis of plant core promoters. Nature Plants *7*, 842-855. <https://doi.org/10.1038/s41477-021-00932-y>.

57.	Loshchilov, I., and Hutter, F. (2017). Decoupled Weight Decay Regularization. International Conference on Learning Representations.

58.	Altschul, S.F., Gish, W., Miller, W., Myers, E.W., and Lipman, D.J. (1990). Basic local alignment search tool. Journal of Molecular Biology *215*, 403-410. <https://doi.org/10.1016/S0022-2836(05)80360-2>.

59.	Bailey, T.L., Johnson, J., Grant, C.E., and Noble, W.S. (2015). The MEME Suite. Nucleic Acids Research *43*, W39-W49. <https://doi.org/10.1093/nar/gkv416>.

60.	Gupta, S., Stamatoyannopoulos, J.A., Bailey, T.L., and Noble, W.S. (2007). Quantifying similarity between motifs. Genome Biology *8*, R24. <https://doi.org/10.1186/gb-2007-8-2-r24>.

61.	Ovek Baydar, D., Rauluseviciute, I., Aronsen, D.R., Blanc-Mathieu, R., Bonthuis, I., de Beukelaer, H., Ferenc, K., Jegou, A., Kumar, V., Lemma, R.B., et al. (2026). JASPAR 2026: expansion of transcription factor binding profiles and integration of deep learning models. Nucleic Acids Research *54*, D184-D193. <https://doi.org/10.1093/nar/gkaf1209>.

62.	Healy, J., and McInnes, L. (2024). Uniform manifold approximation and projection. Nature Reviews Methods Primers *4*, 82. <https://doi.org/10.1038/s43586-024-00363-x>.

63.	Gao, L., Shen, G., Zhang, L., Qi, J., Zhang, C., Ma, C., Li, J., Wang, L., Malook, S.U., and Wu, J. (2019). An efficient system composed of maize protoplast transfection and HPLC–MS for studying the biosynthesis and regulation of maize benzoxazinoids. Plant Methods *15*, 144. <https://doi.org/10.1186/s13007-019-0529-2>.

64.	Loegler, V., Thiele, P., Teyssonnière, E., Tsouris, A., Brach, G., Cruaud, C., Payen, E., Engelen, S., Dunham, M.J., Hou, J., et al. (2025). From genotype to phenotype with 1,086 near telomere-to-telomere yeast genomes. Nature *648*, 649-658. <https://doi.org/10.1038/s41586-025-09637-0>.

65.	Igolkina, A.A., Vorbrugg, S., Rabanal, F.A., Liu, H.-J., Ashkenazy, H., Kornienko, A.E., Fitz, J., Collenberg, M., Kubica, C., Mollá Morales, A., et al. (2025). A comparison of 27 Arabidopsis thaliana genomes and the path toward an unbiased characterization of genetic polymorphism. Nature Genetics *57*, 2289-2301. <https://doi.org/10.1038/s41588-025-02293-0>.

66.	Kang, M., Wu, H., Liu, H., Liu, W., Zhu, M., Han, Y., Liu, W., Chen, C., Song, Y., Tan, L., et al. (2023). The pan-genome and local adaptation of Arabidopsis thaliana. Nature Communications *14*, 6259. <https://doi.org/10.1038/s41467-023-42029-4>.

67.	Lian, Q., Huettel, B., Walkemeier, B., Mayjonade, B., Lopez-Roques, C., Gil, L., Roux, F., Schneeberger, K., and Mercier, R. (2024). A pan-genome of 69 Arabidopsis thaliana accessions reveals a conserved genome structure throughout the global species range. Nature Genetics *56*, 982-991. <https://doi.org/10.1038/s41588-024-01715-9>.

68.	Shumate, A., and Salzberg, S.L. (2021). Liftoff: accurate mapping of gene annotations. Bioinformatics *37*, 1639-1643. <https://doi.org/10.1093/bioinformatics/btaa1016>.

69.	Bavarian, M., Jun, H., Tezak, N., Schulman, J., McLeavey, C., Tworek, J., and Chen, M. (2022). Efficient training of language models to fill in the middle. arXiv preprint arXiv:2207.14255.

70.	Xiao, G., Tian, Y., Chen, B., Han, S., and Lewis, M. (2024). Efficient streaming language models with attention sinks. International Conference on Learning Representations, 21875-21895.

71.	Bao, W., Kojima, K.K., and Kohany, O. (2015). Repbase Update, a database of repetitive elements in eukaryotic genomes. Mobile DNA *6*, 11. <https://doi.org/10.1186/s13100-015-0041-9>.

72.	Chan, P.P., and Lowe, T.M. (2019). tRNAscan-SE: Searching for tRNA Genes in Genomic Sequences. In Gene Prediction: Methods and Protocols, M. Kollmar, ed. (Springer New York), pp. 1-14. 10.1007/978-1-4939-9173-0_1.

73.	Brůna, T., Hoff, K.J., Lomsadze, A., Stanke, M., and Borodovsky, M. (2021). BRAKER2: automatic eukaryotic genome annotation with GeneMark-EP+ and AUGUSTUS supported by a protein database. NAR Genomics and Bioinformatics *3*, lqaa108. <https://doi.org/10.1093/nargab/lqaa108>.

74.	Tegenfeldt, F., Kuznetsov, D., Manni, M., Berkeley, M., Zdobnov, E.M., and Kriventseva, E.V. (2025). OrthoDB and BUSCO update: annotation of orthologs with wider sampling of genomes. Nucleic Acids Research *53*, D516-D522. <https://doi.org/10.1093/nar/gkae987>.

75.	Brůna, T., Lomsadze, A., and Borodovsky, M. (2020). GeneMark-EP+: eukaryotic gene prediction with self-training in the space of genes and proteins. NAR Genomics and Bioinformatics *2*, lqaa026. <https://doi.org/10.1093/nargab/lqaa026>.

76.	Stanke, M., Diekhans, M., Baertsch, R., and Haussler, D. (2008). Using native and syntenically mapped cDNA alignments to improve de novo gene finding. Bioinformatics *24*, 637-644. <https://doi.org/10.1093/bioinformatics/btn013>.

77.	Bateman, A., Coin, L., Durbin, R., Finn, R.D., Hollich, V., Griffiths‐Jones, S., Khanna, A., Marshall, M., Moxon, S., Sonnhammer, E.L.L., et al. (2004). The Pfam protein families database. Nucleic Acids Research *32*, D138-D141. <https://doi.org/10.1093/nar/gkh121>.

78.	Buchfink, B., Reuter, K., and Drost, H.-G. (2021). Sensitive protein alignments at tree-of-life scale using DIAMOND. Nature Methods *18*, 366-368. <https://doi.org/10.1038/s41592-021-01101-x>.

79.	Jones, P., Binns, D., Chang, H.-Y., Fraser, M., Li, W., McAnulla, C., McWilliam, H., Maslen, J., Mitchell, A., Nuka, G., et al. (2014). InterProScan 5: genome-scale protein function classification. Bioinformatics *30*, 1236-1240. <https://doi.org/10.1093/bioinformatics/btu031>.

80.	Haas, B.J., Delcher, A.L., Wortman, J.R., and Salzberg, S.L. (2004). DAGchainer: a tool for mining segmental genome duplications and synteny. Bioinformatics *20*, 3643-3646. <https://doi.org/10.1093/bioinformatics/bth397>.

81.	Song, B., Marco-Sola, S., Moreto, M., Johnson, L., Buckler, E.S., and Stitzer, M.C. (2022). AnchorWave: Sensitive alignment of genomes with high sequence diversity, extensive structural polymorphism, and whole-genome duplication. Proceedings of the National Academy of Sciences *119*, e2113075119. <https://doi.org/10.1073/pnas.2113075119>.

82.	Li, H. (2021). New strategies to improve minimap2 alignment accuracy. Bioinformatics *37*, 4572-4574. <https://doi.org/10.1093/bioinformatics/btab705>.

83.	Bradbury, P.J., Zhang, Z., Kroon, D.E., Casstevens, T.M., Ramdoss, Y., and Buckler, E.S. (2007). TASSEL: software for association mapping of complex traits in diverse samples. Bioinformatics *23*, 2633-2635. <https://doi.org/10.1093/bioinformatics/btm308>.

84.	van der Auwera, G., and O'Connor, B.D. (2020). Genomics in the Cloud: Using Docker, GATK, and WDL in Terra (O'Reilly Media, Incorporated).

85.	Danecek, P., Bonfield, J.K., Liddle, J., Marshall, J., Ohan, V., Pollard, M.O., Whitwham, A., Keane, T., McCarthy, S.A., Davies, R.M., and Li, H. (2021). Twelve years of SAMtools and BCFtools. GigaScience *10*, giab008. <https://doi.org/10.1093/gigascience/giab008>.

86.	Jeffares, D.C., Jolly, C., Hoti, M., Speed, D., Shaw, L., Rallis, C., Balloux, F., Dessimoz, C., Bähler, J., and Sedlazeck, F.J. (2017). Transient structural variations have strong effects on quantitative traits and reproductive isolation in fission yeast. Nature Communications *8*, 14061. <https://doi.org/10.1038/ncomms14061>.

87.	Cingolani, P., Platts, A., Wang, L.L., Coon, M., Nguyen, T., Wang, L., Land, S.J., Lu, X., and Ruden, D.M. (2012). A program for annotating and predicting the effects of single nucleotide polymorphisms, SnpEff. Fly *6*, 80-92. <https://doi.org/10.4161/fly.19695>.
